# Species delimitation in an intractable syngameon: Bringing order to the polyphyletic *Heuchera americana* group

**DOI:** 10.64898/2026.02.27.708381

**Authors:** N.J. Engle-Wrye, R.A. Folk

**Affiliations:** Mississippi State University

## Abstract

Species are the fundamental analytical units of evolutionary processes; thus evidence-based species delimitation is a crucial step for understanding species radiations. However, the task of delimiting species is particularly challenging in the context of a syngameon—a group of distinct, but closely related species that have incomplete reproductive isolation and frequently hybridize in nature. This problem is further exacerbated by the presence of cryptic species—species that are phenotypically distinct, though difficult to distinguish with gross morphology alone. *Heuchera* subsect. *Heuchera* comprises both clear and cryptic species within a syngameon that has seen study from morphological, experimental, and phylogenetic aspects. This group has long been recognized for its taxonomic complexity, namely two recognized hybrid zones with extreme morphological variation and persistent non-monophyly among parental populations. Here, we reassess species limits within *Heuchera* subsect. *Heuchera*, focusing on the hybrid complex between *H. americana* and *H. richardsonii* and adjacent *H. americana* populations. We use a multipronged approach with deep population-level sampling to 1) assess the genetic structure of 655 individuals across the geographic range of the *H. americana* group to identify genetic lineages and 2) assess the phenotypic diagnosability of these lineages. Despite extensive admixture and gene tree conflict, we find multiple cohesive lineages with diagnosable phenotypes. We recognize five species and three varieties within the *H. americana* group, one new and four resurrected. Our results demonstrate that even highly reticulate syngameons can be partitioned into meaningful taxonomic units with multiple lines of evidence.

## Introduction

Species delimitation is a difficult task that has continued unabated for nearly 300 years, only recently receiving some relief with the advent of molecular techniques (Bickford et al., 2007). Challenges to species delimitation comprise generally (1) a shortage of primary information on biodiversity that would inform delimitation and (2) challenges in turning primary information measured from specimens into a delimitation decision. Plants are particularly challenging in respect to (2) due to the prevalence of heterogeneous evolutionary processes such as hybridization and polyploidy, which inherently defy discrete hierarchical categorization logic regardless of the data or decision calculus applied to the data. Molecular techniques have been revolutionary for modern species delimitation (Mallet, 2005; De Queiroz, 2007; Soltis and Soltis, 2009; Yang and Rannala, 2010; Fujita et al., 2012), yet molecular information in itself does not simplify a complex reality. The interpretation of molecular data in species problems comes with inherent shortcomings ranging from the inability to directly assess prezygotic isolation barriers (Bickford et al., 2007) to oversplitting species and confusing populations as specific entities due to differences in species concept and statistical model violations (Chan et al., 2017; Jackson et al., 2017; Sukumaran and Knowles, 2017; Chambers et al., 2025). More pointedly, molecular data offer promise in teasing apart complex evolutionary complexes (Folk et al., 2018b), but identifying this complexity brings us no closer to a taxonomic decision without a clear sense of how to recognize taxa and apply ranks to the results.

Leaving aside the (substantial) methodological challenges, two further primary empirical challenges can be identified in application of molecular data to species delimitation in plants: (1) phenotypically cryptic genetic differentiation and (2) lack of completely discontinuous variation in syngameons. In respect to challenge (1), “cryptic species” are evolutionarily independent lineages that, depending on the species criterion, are either phenotypically indistinguishable or distinct but challenging to distinguish in the field. While by definition cryptic species are at best distinct for phenotypes that are difficult for humans to observe in the field, this often proves less so for pollinators and other ecological interactors detecting volatile compounds and other phenotypes beyond direct human sense. Cryptic species only rarely display syntopy (local co-occurrence of species that might readily hybridize) such that they contribute noticeably to regional and global diversity but insignificantly to the diversity of a single site (Delić et al., 2017), a further type of crypsis for locally oriented studies. Cryptic species can be integral to meaningful conservation practice and to basic science, as they (1) could have ecologically distinct function not necessarily in proportion to ease of identification, and (2) could elucidate diversification at the micro/macroevolutionary boundary as the qualities of “good species” accrue (Bickford et al., 2007; Delić et al., 2017; Fišer et al., 2018). Because the ecological niches of cryptic species are often similar but may also differ in environmental responses, they may serve as a functional redundancy allowing ecological communities to absorb the shock of changing inputs and buffer themselves against detrimental environmental changes (Hubbell, 2005; Fišer et al., 2018). Thus phenotypic crypsis is generally unproblematic for species delimitation and improving estimates of diversity, distribution, and conservation status (Delić et al., 2017; Fišer et al., 2018; Vivien et al., 2025), whereas the contrasting case of true lack of ecological role distinction is fundamentally problematic for species as the basis for biodiversity value and management decisions; see (Freudenstein et al., 2016). Unfortunately for present-day conservation purposes, cryptic species are often perceived to have only a negligible impact when generating numerical selection criteria for prioritizing preservation of conservation sites.

In respect to challenge (2), “taxonomically difficult” plant groups often comprise networks of phenotypically distinct species linked by occasional gene flow, famously typified by oaks (Cannon and Petit, 2019); Lotsy first termed these networks “syngameons” (Lotsy, 1925). Syngameons represent a direct challenge to the hierarchical and discrete logic underlying Linnean taxonomy (“linneons”) as Lotsy identified a century ago. Syngameons also fly in the face of the underlying motivation for recognizing biological species as isomorphic with reproductive communities (Cannon and Petit, 2019). Syngameons are incompatible with reproductive barrier logic because “good species” in this context (phenotypically discontinuous lineages with secondary gene flow) and their hybrids are potentially easily diagnosable and ecologically distinct, but Biological Species Concept logic would lead to a lumping decision regardless of the magnitude of distinction or even whether the parents are immediately related (leading also to hierarchy violation). On a more practical level, moderate gene flow levels combined with the presence of stable phenotypic combinations lead to benefits and costs (essentially, sensitivity vs. specificity as this is literally a classification problem) whether we lump or split. Lumping would lead to more easily categorizing and navigating genetic and phenotypic variation in a syngameon, whereas it simultaneously would conceal partially discontinuous variation for basic and applied purposes such as conservation.

*Heuchera* (Saxifragaceae) forms an excellent system for grappling with hybridization problems given evolutionary complexity at various phylogenetic levels (Folk et al., 2016, 2018a, c). *Heuchera americana* in particular has long been known to be a polymorphic complex (Rosendahl et al., 1936; Wells, 1979, 1984) and stands as one of the thorniest delimitation problems in the genus. The first authoritative treatment by Rosendahl et al. (1936) suggested that there were six well marked morphotypes within *H. americana* that warranted varietal ranks. Wells et al. (1984) later reexamined these varieties for geographic, reproductive, and morphological cohesiveness, and while acknowledging the unusually high variation judged them to be too continuous or overlapping in all traits. As a result, she collapsed varietal ranks and left a polymorphic *Heuchera americana*, only recognizing three subspecific taxa in total. Given that two of these recognized taxa involve hybrids with close but non-sister species (*H. pubescens* and *H. richardsonii*), the complex comprises the entire containing subclade, *Heuchera* subsect. *Heuchera.* With the advent of molecular data Folk et al. (2018a) sequenced hundreds of nuclear loci with limited taxon sampling, finding *H. americana* is non-monophyletic and confirming previous hypotheses that *H. americana* has a history of interspecific gene flow (Rosendahl et al., 1936). Given the presence of polymorphism, apparent polyphyly, and widespread gene flow, we hypothesized that *H. americana* was actually a group of cryptic taxa and hybrids whose seemingly intractable morphological and genetic variation are the classic symptoms (Cannon and Petit, 2019; Buck and Flores-Rentería, 2022) of a syngameon.

Using the “phenophyletic view” (Freudenstein et al., 2016) as our guiding principle for dividing a syngameon into manageable units, we seek to map the *Heuchera americana* syngameon and identify lineages within *Heuchera* subsect. *Heuchera* with cohesive phenotypes to delimit species. Based on historical taxonomic treatments and preliminary phylogenetic data, we hypothesized that: 1) There are at least two cryptic lineages within what is currently taxonomically described as *Heuchera americana*, distinct in genetic lineage and phenotype, 2) the cryptic taxa delimited from *H. americana* would be morphologically equivalent to at least two of the widely distributed varieties discussed by Rosendahl et al. (1936); namely *H. americana* var. *brevipetala*, *H. americana* var. *calycosa*, and/or *H. americana* var. “*typica*” (hereafter called *H. americana* var. *americana*), 3) that *Heuchera richardsonii* also contains at least two cryptic lineages related to those discussed by Rosendahl et al. (1936), and that 4) that the widespread and morphologically variable *H. americana* var. *hirsuticaulis* should remain recognized as a hybrid taxon closer to *H. americana*.

## Materials and methods

### Species concept

For the purpose of species delimitation, we chose the “phenophyletic view” of Freudenstein et al. (2016). This concept considers a species to be a lineage or collection of related lineages (here, historically connected metapopulations) that is diagnosable through shared phenotypes. Notably, hybridization is unproblematic under this concept as “lineage” is conceived in population genetic terms and does not correspond to a cladistic concept. This species delimitation criterion is related to the general lineage concept and its use of lineage (De Queiroz 1998) but differs in requiring, rather than allowing as evidence, phenotypic diagnosability.

### Taxon sampling

We conducted both genetic and morphological sampling (methods discussed below), we sought to collect representative materials from all potential species and/or varieties across as much of their ranges as feasible. These materials were obtained through field collections and dried herbarium specimens. Field collections primarily targeted the hybrid zone between *H. americana* and *H. richardsonii* as described by Wells (1984) and geographic regions of morphologically well marked varieties, from both species, as hypothesized by Rosendahl et al. (1936). Genetic sampling ultimately comprised 532 ingroup specimens (those corresponding to *H. americana*, *H. richardsonii*, and hybrids) and 123 outgroup specimens (including other members of *Heuchera* subsect. *Heuchera* as well as taxa outside this clade), for a total of 655 specimens (after application of filtering steps; see below). While many of the varieties mentioned in Rosendahl et al. (1936) have little history of subsequent recognition, we implement these as *a priori* hypotheses of distinct cryptic species. While our genetic sampling was as broad as possible, morphometric sampling targeted a subset of specimens representing morphologies of the varieties mentioned in Rosendahl et al. (1936). Measurements were conducted on 139 herbarium specimens also included in our phylogenetic analyses.

Wells (1984) noted many intermediate morphologies among the taxa recognized in Rosendahl et al. (1936) and we encountered similar difficulties in applying the taxonomic key reported in the latter; as such initial species assignments were made using reported geographic distributions within the “*H. americana* group” (hereafter defined as only *H. americana*, *H. richardsonii*, their varieties and hybrids as recognized by Rosendahl et al (1936)), and morphological identifications for other taxa. We found this to be an acceptable starting point that captures the main taxon gradients and renders genetic and morphological data as independent tests of these *a priori* hypotheses. In addition to these prior hypotheses we delimited a further *a priori* hypothesis comprising *H. americana* individuals from the Ocoee River Gorge and Smoky Mountains of Tennessee with hairy petioles; first pointed out by Wells (1984) and referred to in (Folk and Freudenstein, 2014) as *H. americana* “var. *heteradenia*,” hereafter *H. fumosimontana* (see nomenclatural notes in the Taxonomic Treatment below).

### Phylogenetic construction

We used the ASTRAL phylogeny from BIORXIV/2026/708067 as a guideline for downstream phylogenetic analyses and inferences. This phylogeny was inferred from a customized panel of 277 low-copy nuclear loci specific to *Heuchera* (details following Folk et al. (2015). From the 532 ingroup samples, we collected 297 *H. americana* var. *americana*, 135 *H. americana* var. *hirsuticaulis*, and 100 *H. richardsonii* accessions (according to the delimitation of Wells 1984; accessions were also geographically annotated following Rosendahl et al. 1936). This analysis evinced substantial gene tree conflict; in an attempt to increase support and limit the influence of interspecific gene flow, we then inferred a second ASTRAL phylogeny (https://github.com/njenglewrye/Heuchera-americana-group-species-manuscript/blob/main/Phylogenetics/ASTRAL_pure/ASTRAL_pure.sh) newly reported here, to include only individuals recovered as “pure” (taxon assignment > 90%) by fastSTRUCTURE, see below. According to the system of Wells (1984), this ancestry-filtered phylogeny contained 216 of the focal taxa (101 *H. americana* var. *americana*, 86 *H. americana* var. *hirsuticaulis*, and 29 *H. richardsonii*) and 53 individuals belonging to outgroup taxa; for a grand total of 269 specimens. According to the system of Rosendahl et al. (1936), the 216 individuals comprised 69 *H. americana* var. *hirsuticaulis*, 17 *H. richardsonii* var. *grayana*, 29 *H. richardsonii* var. *richardsonii*, 15 *H. fumosimontana* (see Taxonomic Treatment), 20 *H. americana* var. *calycosa*, 41 *H. americana* var. *brevipetala*, and 25 *H. americana* var. *americana*.

### Population genetics

The same trimmed and filtered sequence read data underlying the phylogeny from BIORXIV/2026/708067 obtained and aligned to an assembly reference based on *H. alba* H63-1 (derived from assemblies reported in (Folk et al., 2016)) using BWA-MEM. BAM files were sorted into mapped and unmapped read groups, duplicate reads marked, and read groups added using GATK and samtools. Per-sample variant calling was performed with GATK HaplotypeCaller in GVCF mode (https://github.com/njenglewrye/Heuchera-americana-group-species-manuscript/blob/main/Phylogenetics/pop_gen/1_gatk_variantcall.sh). Sample GVCFs were then jointly genotyped using gatk. Given that unfiltered multilocus data violate independence assumptions, genome-wide linkage disequilibrium (LD) then was estimated on the unpruned SNP dataset using PopLDdecay, calculating pairwise LD (*R*²) as a function of physical distance. The resulting LD decay curves were visualized and used to guide SNP thinning. Linkage disequilibrium (LD) decays rapidly in this population genetic dataset (Fig. S1); a typical pruning benchmark is 50% decay from the initial value and LD drops from an (r^2^ ∼ 0.07) to (r^2^ ∼ 0.035) at a distance of ∼50 bp. We then used vcftools to restrict variants to SNPs, filtered them for a minor allele frequency ≥ 0.05, maximum missing data ≤ 50%, and thinned one SNP for every 50 bp to minimize linkage disequilibrium, resulting in a set of 6,733 SNPs from the initial 158,683. We then assessed population structure via fastSTRUCTURE, using the default admixture model with correlated allele frequencies. Analyses were conducted for *K* = 2–13 (*K* = genetic cluster, with the maximum of 13 representing the number of *a priori* taxa for *Heuchera* subsect. *Heuchera*) and then the optimal *K* was determined (*K =* 11) using the chooseK.py script. The *K* genetic clusters represent hypothetical ancestral populations from which samples derive through unique or mixed ancestry; we will refer to these units as “ancestral populations” hereafter. Ancestry coefficients (*Q* values) were visualized using pophelper in R (version 4.5.0). Individuals with a maximum single ancestry assignment ≥ 90% were considered “pure,” whereas those below this threshold were considered admixed.

A principal component analysis (PCA) was also run on the pruned SNP data to further visualize grouping in the data using the adegenet package in R. Population genetic differentiation was also quantified in R using the hierfstat package. Pairwise fixation indices (*F*_st_), which measure allelic fixation among *a priori* populations were computed according to (Weir and Cockerham, 1984) and pairwise *F*_st_ matrices were visualized as heat maps using pheatmap.

### Bayesian estimates of lineages

Species delimitation via BPP (Yang and Rannala, 2010) was used as a model-based approach to estimate lineage distinctness. Species delimitation was performed using the reversible-jump MCMC (rjMCMC) algorithm (Yang, 2015) while jointly estimating the species tree. Thirteen putative species were included, as in STRUCTURE and other population genetic analysis, and an initial guide tree reflecting the relationships inferred from phylogenomic analyses was specified. We used pruned alignments from BIORXIV/2026/708067 and analyzed the sequence data directly without removal of ambiguous sites. Analyses used a prior of uniform rooted trees with population size parameters (θ) assigned using an inverse-gamma prior with shape α = 3 and scale β = 0.04. Divergence times (τ) were also assigned shape α = 3 while scale was set to β = 0.2 following developer suggestions. For internal divergence times, a Dirichlet prior was used with a beta prior on the root age. Markov chain Monte Carlo analyses were set to automatically finetune and run for 100,000 samples, sampling every two iterations following a burn-in of 10,000 iterations. We implemented four identical runs and examined posterior probabilities of species delimitations to assess convergence and suitable rjMCMC mixing behaviour. Posterior probabilities were also used to evaluate support for alternative species hypotheses.

### Morphometrics

High resolution imagery was obtained from MEM, MISSA, UNA, UNC, and TEX for initial comparisons and morphometric measurements using ImageJ. Additionally, visits were made to each herbarium to examine specimens in person, take leaf samples for phylogenetic analyses, and take supplemental measurements to corroborate the accuracy of ImageJ measurements. In total 338 specimens were imaged and measured, but only 139 specimens had floral parts measurable using ImageJ while also at the prescribed developmental stage for morphological analyses (in full anthesis and not yet in fruiting with intact leaves). It is important for consistency to measure flowers in full anthesis because *Heuchera* flowers are protandrous and highly dynamic in size, morphology, color, and attitude throughout their developmental stages (Wells, 1984). For a list of vouchers used see (https://github.com/njenglewrye/Heuchera-americana-group-species-manuscript/tree/main/Morphometrics). The morphological traits measured included: longest petiole hair length, shortest petiole hair length (as petiole hairs in *Heuchera* often show two distinct length classes; see (Fernald, 1942), primary tooth lengths of secondary lobes, primary tooth widths of secondary lobes, flower lengths, adaxial hypanthium lengths, abaxial hypanthium lengths, hypanthium widths at ovary, stamen exsertion lengths past sepal margins, pedicel lengths, and cymule branch lengths (these lengths are each given definitions in Engle-Wrye (2023). Three replicates of each metric were taken from distinct structures on a specimen when possible; in total 5,301 individual measurements were made. Using these metrics, we implemented linear discriminant analyses (LDAs) across all taxa and within a subset of taxa exhibiting extensive morphological overlap, followed by a jackknifed classification success test. LDA was used rather than PCA to render the ordination problem as hypothesis-oriented. Multivariate differentiation among taxa was further evaluated using MANOVA, followed by pairwise PERMANOVA (adonis) tests and univariate boxplot analyses.

## Data availability

Code and datasets to perform the analyses noted above are available at https://github.com/njenglewrye/Heuchera-americana-group-species-manuscript/tree/main.

## Results

### Curated phylogeny

The original ASTRAL phylogeny of BIORXIV/2026/708067 rendered a monophyletic *H. richardsonii*, if assigned hybrids are excluded, and a paraphyletic *H. americana*; while the hybrid taxa had typical hybrid topologies, often split between and nesting in subclades of parental taxa. Among the subclades there was low to high support (LPP > 0.5). The backbone had low support (LPP < 0.5), reflecting high conflict in the data. Our new ASTRAL phylogeny (Fig. 1), pruned to include only individuals recovered as “pure” by fastSTRUCTURE, improved overall backbone and subclade support (LPP > 0.5). Clades corresponding to *Heuchera richardsonii* var. *richardsonii*, *H. richardsonii* var. *grayana*, *H. americana* var. *hispida* and *H. americana* var. *calycosa* formed moderately to well supported (LPP > 0.5) monophyletic clades. All other putative taxa were non-monophyletic or unresolved and formed mostly well supported subclades.

**Figure 1.**
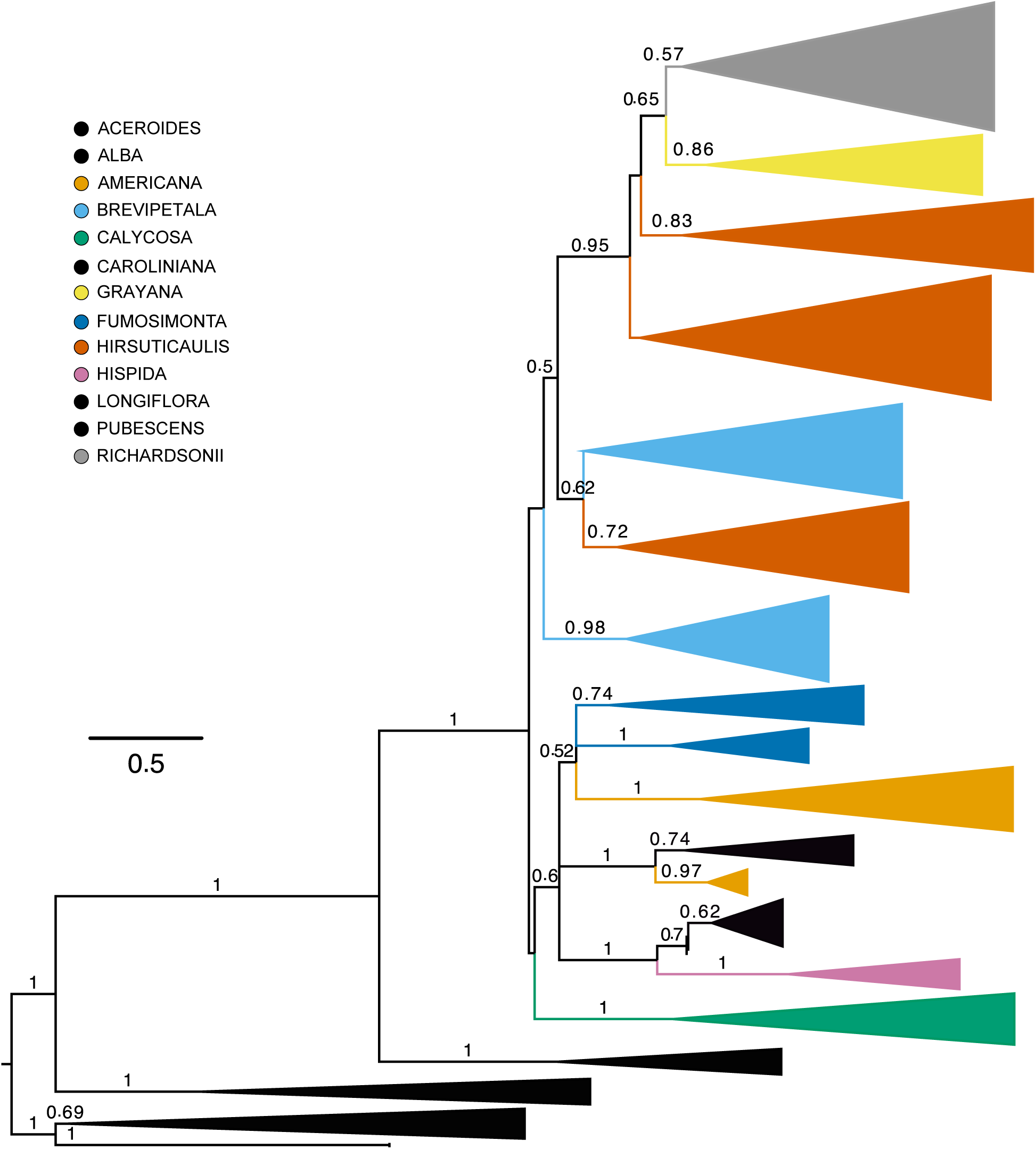
ASTRAL phylogeny of “pure” taxa. ASTRAL phylogeny inferred using a subset of genetic data from BIORXIV/2026/708067, to include only individuals recovered as “pure” (>90% assignment probability) by fastSTRUCTURE. Low support values (LPP < 0.5) were removed. Names in the legend reflect the highest-resolution epithet, either infraspecific (varietal) or specific

### Population genetics

fastSTRUCTURE reveals a complex genetic landscape with high levels of admixture and few “pure” individuals (Fig. 2). fastSTRUCTURE decisively supported 11 distinct ancestral populations underlying the data; considering a pure assignment cutoff as 90%, 44.4% of the individuals are pure and 55.6% are mixed among the ancestral populations. Among the members of *Heuchera* sect. *Heuchera*, *H. alba, H. richardsonii, H. caroliniana,* and *H. fumosimontana* consisted almost entirely of genetically pure individuals. Taxa comprising the remainder of the complex show at least some degree of admixture. This includes all included individuals of *H. longiflora* and *H. pubescens,* both morphologically well-marked taxa with more than a century of recognition, as well as the remaining *H. americana* segregates (Fig. 2). Three segregate taxa recognized here correspond to Wells’ (1984) hybrid zone complexes: *H. americana* var. *hirsuticaulis* and *H. richardsonii* var. *grayana* correspond to those members of the *H. americana* × *H. richardsonii* hybrid zone that more closely resemble the *H. americana* or *H. richardsonii* parent respectively (Rosendahl et al., 1936; see also morphological discussion in Wells, 1984). These were found to correspond to three separate ancestral populations in fastSTRUCTURE (Fig. 2, orange, dark blue, red); interestingly, these ancestral populations are not found except as minor admixture in the parental taxa, suggesting the hybrids are genetically distinct. *H. americana* var. *hispida*, the remaining hybrid, was considered by Wells to represent *H. americana* × *H. pubescens* and was recovered as intermixed with no evidence of a unique gene pool. Further ancestral populations contributed uniquely to *H. americana* var. *americana*, *H. americana* var. *brevipetala*, and *H. americana* var. *calycosa* (Fig. 2, green, yellow, pink, and brown) but do not perfectly delimit these.

**Figure 2.**
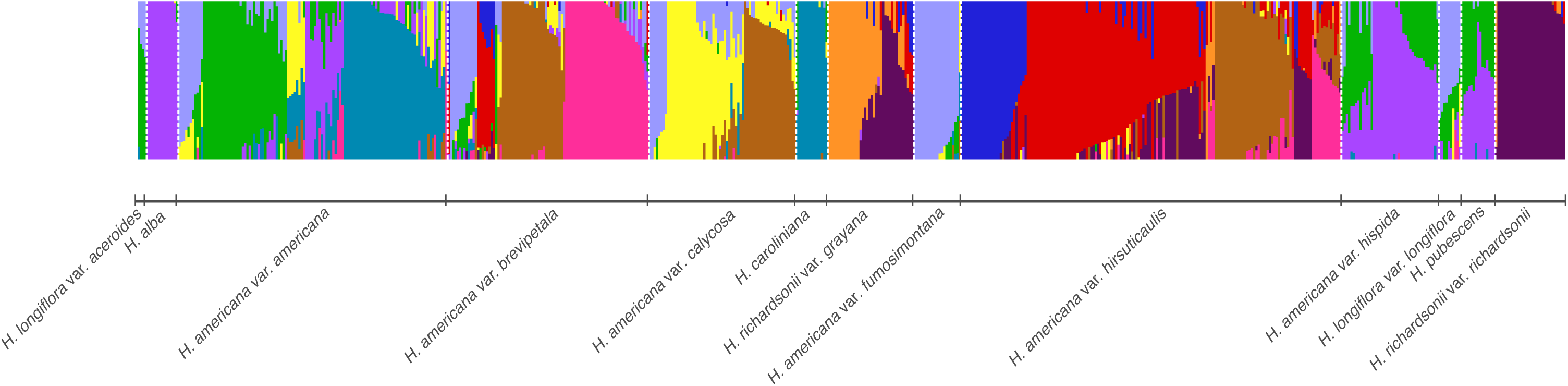
fastSTRUCTURE ancestral population assignments. fastSTRUCTURE results showing ancestral population assignments (optimal *K =* 11), delimited by tick marks on the x-axis, of individuals with each individual column being one specimen, and proportion of shared genetic clusters or inferred ancestral population indicated by shared colors and proportion of the y-axis each color occupies. Taxon assignments of individuals follow *a priori* assignments.

The principal component analysis (PCA) of pruned SNP data (Fig. 3) shows two primary clusters within the *H. americana* group. One centers on *H. richardsonii* var. *richardsonii* and its putative hybrid segregate *H. richardsonii* var. *grayana*, while the other centers on *H. americana* and all other members of the complex. *Heuchera americana* var. *hirsuticaulis*, the putative hybrid segregate of *H. americana*, and *H. americana* var. *brevipetala* occupy a line connecting the majority of *H. americana* and *H. richardsonii*, much as morphology would suggest (Rosendahl et al., 1933, 1936). Pairwise comparisons of fixation indices (Fig. 4) show varying degrees of population structure and gene flow among taxa, particularly delimiting *H. richardsonii* and its hybrid derivatives. Notable pairwise comparisons are interpreted below in the Discussion.

**Figure 3.**
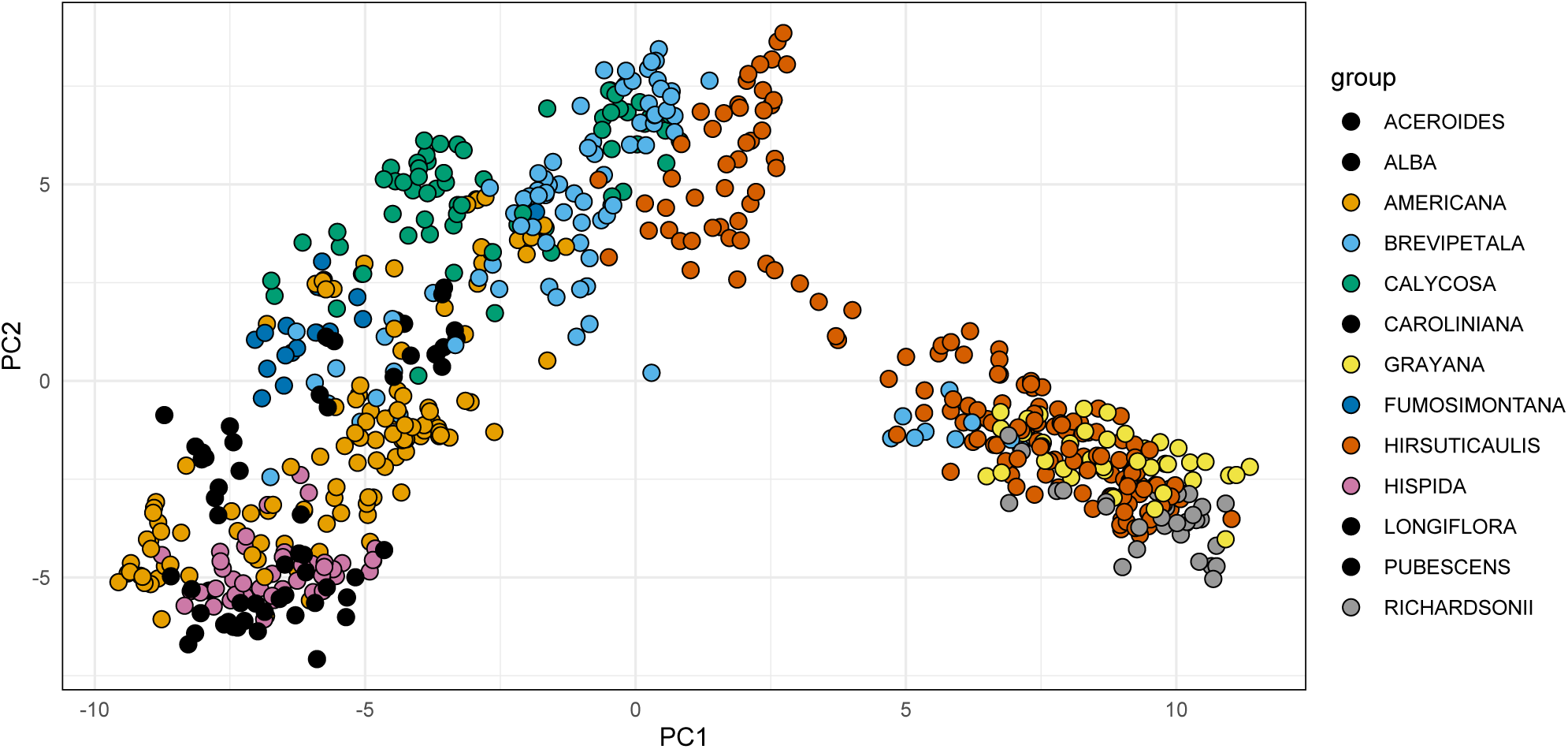
Principal component analysis of genetic variation. Principal component analysis (PCA) of genetic variation among individuals. Each point represents an individual colored according to its *a priori* assigned species in the legend on the right. Names in the legend reflect the highest-resolution epithet, either infraspecific (varietal) or specific.

**Figure 4.**
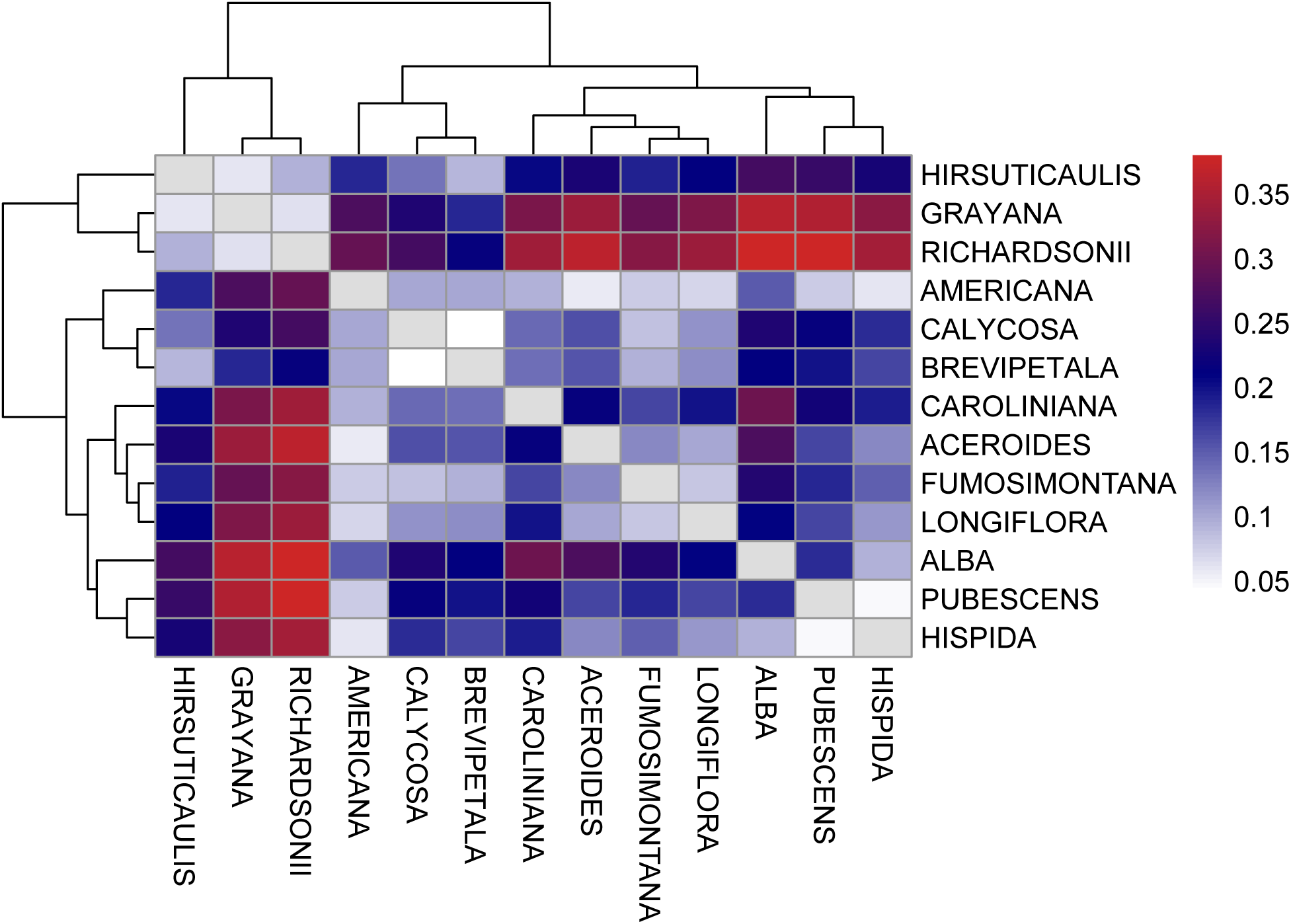
Pairwise comparisons of fixation indices (F_st_) among populations. Names on the axes reflect the highest-resolution epithet, either infraspecific (varietal) or specific. Values between 0-0.05 indicate very little differentiation or high degrees of geneflow, values between 0.05-0.15 indicate moderate differentiation with some population structure and geneflow, values between 0.15-0.25 indicate strong differentiation with clear population structure, and values > 0.25 indicate very strong differentiation of highly divergent or isolated populations. Taxonomic assignments are *a priori.* Dendrograms on the left-hand and top sides indicate genetic distance.

### Bayesian estimates of lineages

Two independent MCMC chains run in 130,438 generations together. The best model predicted five species (Fig. 5) with a posterior probability (PP) of 0.507536. *Heuchera fumosimontana* was the only taxa not lumped with another and had the highest support (PP = 0.97). The estimated species with the second-best support, involving focal members of the *H. americana* group, included *H. richardsonii*, *H. richardsonii* var. *grayana*, and *H. americana* var. *hirsuticaulis* (PP = 0.85). Finally, *H. americana* var. *americana* and *H. caroliniana* were assigned as one species (PP = 0.51) and *H. americana* var. *brevipetala* and *H. americana* var. *calycosa* were assigned as another (PP = 0.51).

**Figure 5.**
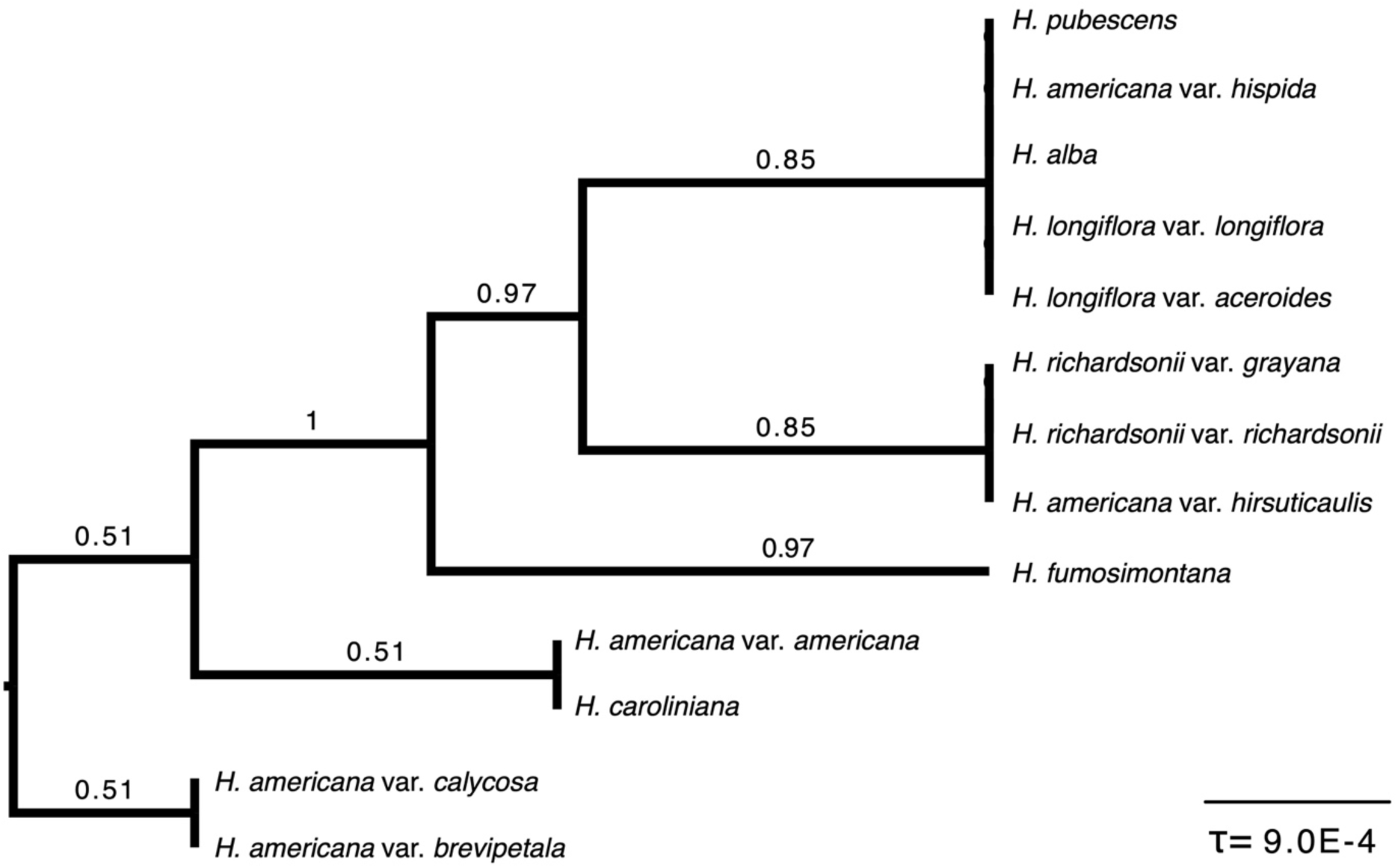
BPP inferred species limits. Best supported model of species limits inferred by BPP; posterior probability of 0.507536 for five species. Divergence times (τ) for each node are reported in substitutions per site.

### Morphometrics

The foregoing molecular analyses were performed on *a priori* species hypotheses; hereafter assignments were corrected to reflect *a posteriori* hypotheses so as to provide a direct test of morphological distinction of units delimited by molecular data (i.e., the phenophyletic view). This was done by consulting the population assignment probability table output by fastSTRUCTURE and adjusting taxon membership. Conflict of geographic designations with *a priori* hypotheses was mostly minor for available specimens, but one individual of *H. richardsonii* var. *grayana* was reassigned to *H. richardsonii* var. *richardsonii*, one *H. americana* var. *brevipetala* individual was reassigned to *H. americana var. hispida,* and two *H. americana* var. *calycosa* were reassigned to *H. americana* var. *brevipetala*.

A linear discriminant analysis of all measured taxa was implemented to confer a direct test of morphological distinctness. Much like the molecular PCA, the morphological LDA showed strong separation of *H. americana* var. *hirsuticaulis, H. richardsonii* var. *richardsonii,* and *H. richardsonii* var. *grayana*, forming the main axis of separation in the complex. *Heuchera fumosimontana* also exhibited good separation while the remaining members of the *H. americana* group and *H. americana* var. *hispida* showed substantial overlap (Fig. 6). A jackknifed classification success test had an accuracy of 0.70, suggesting the linear discriminant had some success distinguishing several of the *H. americana* complex taxa. When subsetted to the overlapping members of the *H. americana* group and *H. americana* var. *hispida* (a hybrid variety of *H. americana*), an LDA showed better separation (Fig. 7), but resulted in a lower jackknifed classification success of 0.65; suggesting that *H. fumosimontana*, *H. richardsonii*, *H. richardsonii* var. *grayana*, and *H. americana* var. *hirsuticaulis* are easily distinguished, but the remaining segregates of *H. americana* exhibit overlap as measured. However, a MANOVA (which tests for mean difference rather than implementing a classification model) indicated that species differed significantly in overall morphology (Pillai = 2.63, p < 2.2 × 10^-16^) and investigation of individual predictors in the MANOVA indicate that all response variables were significant (Table 1). Species comparisons via pairwise PERMANOVA (adonis) (Table 2) indicated significant differences among all species pairs except for *H. americana* var. *hispida* vs. *H. richardsonii*, *H. richardsonii* var. *grayana* vs. *H. richardsonii* var. *richardsonii*, and all comparisons involving *H. americana* var. *americana*, except for *H. americana* var. *hirsuticaulis* vs. *H. americana* var. *americana*. Univariate boxplots (Fig. 8) illustrate overlap and divergences of key morphological traits; the lengths of flowers, stamens, petiole hairs, and upper laminar hairs provided the most discrimination information among the taxa.

**Figure 6.**
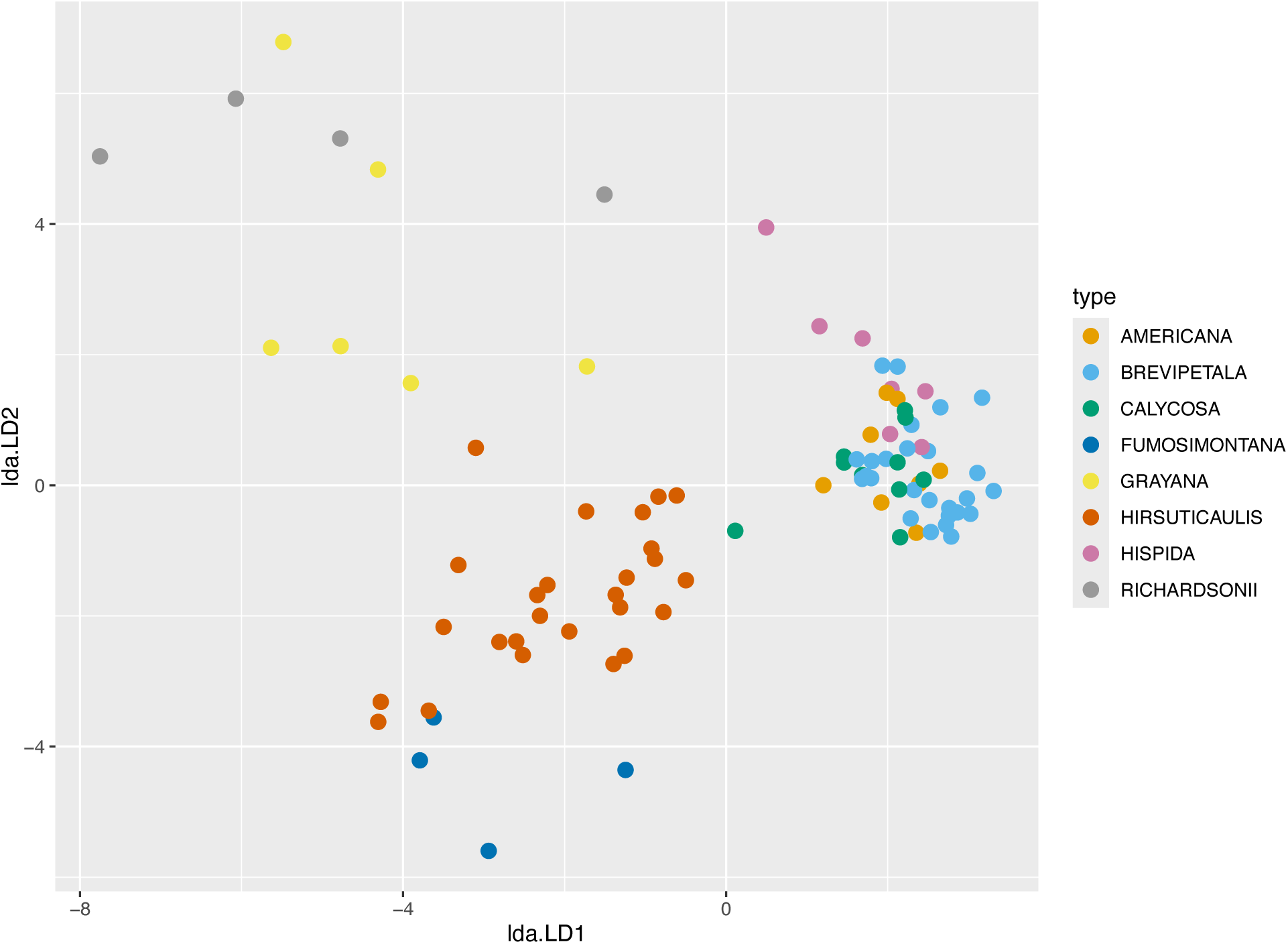
Morphological linear discriminant analysis of all measured taxa. Labels in the legend follow *a posteriori* (post-molecular) reassignments to directly assess the hypothesis of morphological distinction. Names in the legend reflect the highest-resolution epithet, either infraspecific (varietal) or specific.

**Figure 7.**
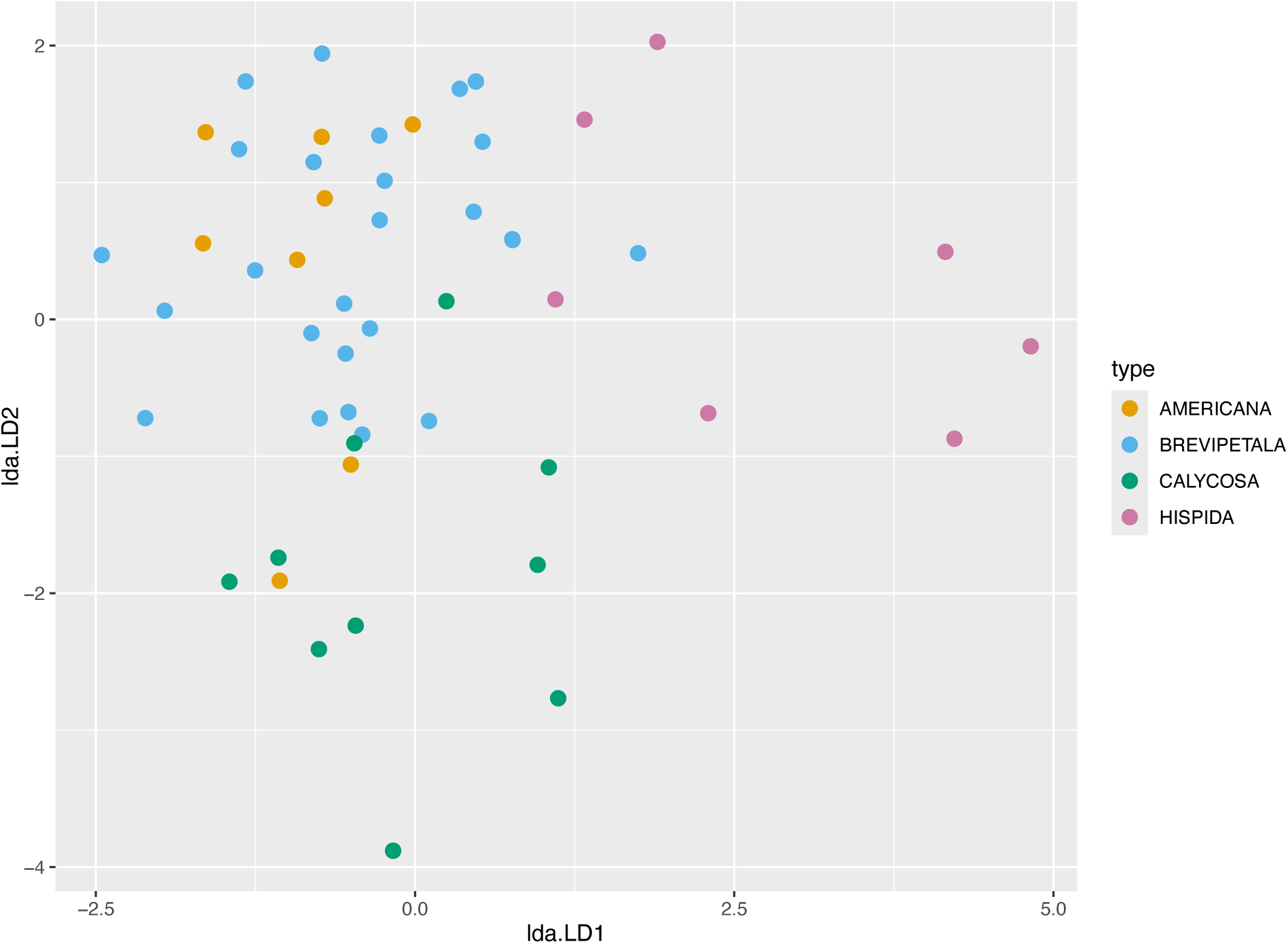
Morphometric linear discriminant analysis of *H. americana* varieties. Linear discriminant analysis of morphological data subsetted to *H. americana* varieties; details otherwise follow Fig. 6.

**Figure 8.**
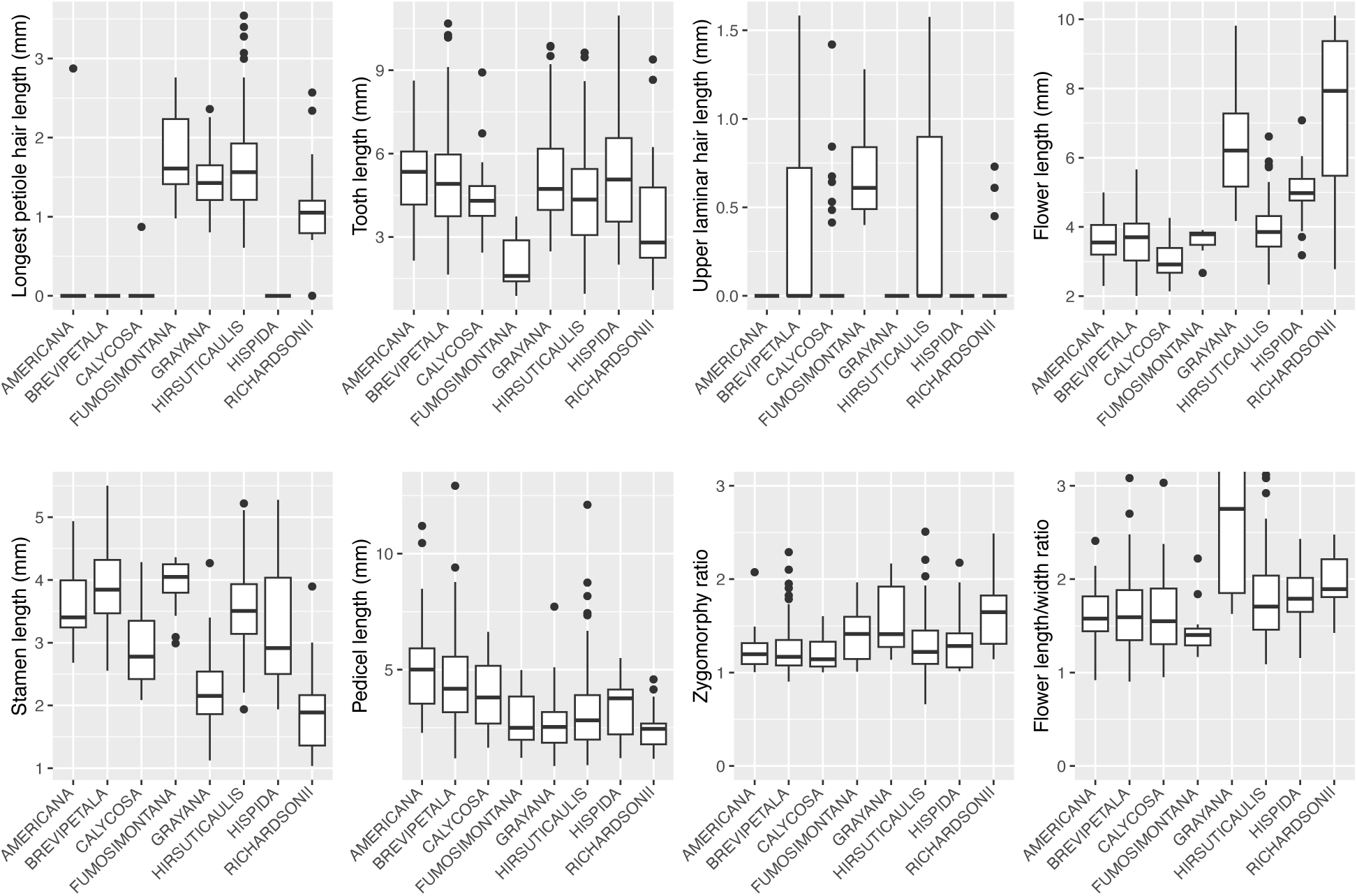
Univariate boxplots of morphological characteristics. Each boxplot has the species on the x-axis and the morphological trait on the y-axis. Names on the x-axes reflect the highest-resolution epithet, either infraspecific (varietal) or specific.

**Table 1.**
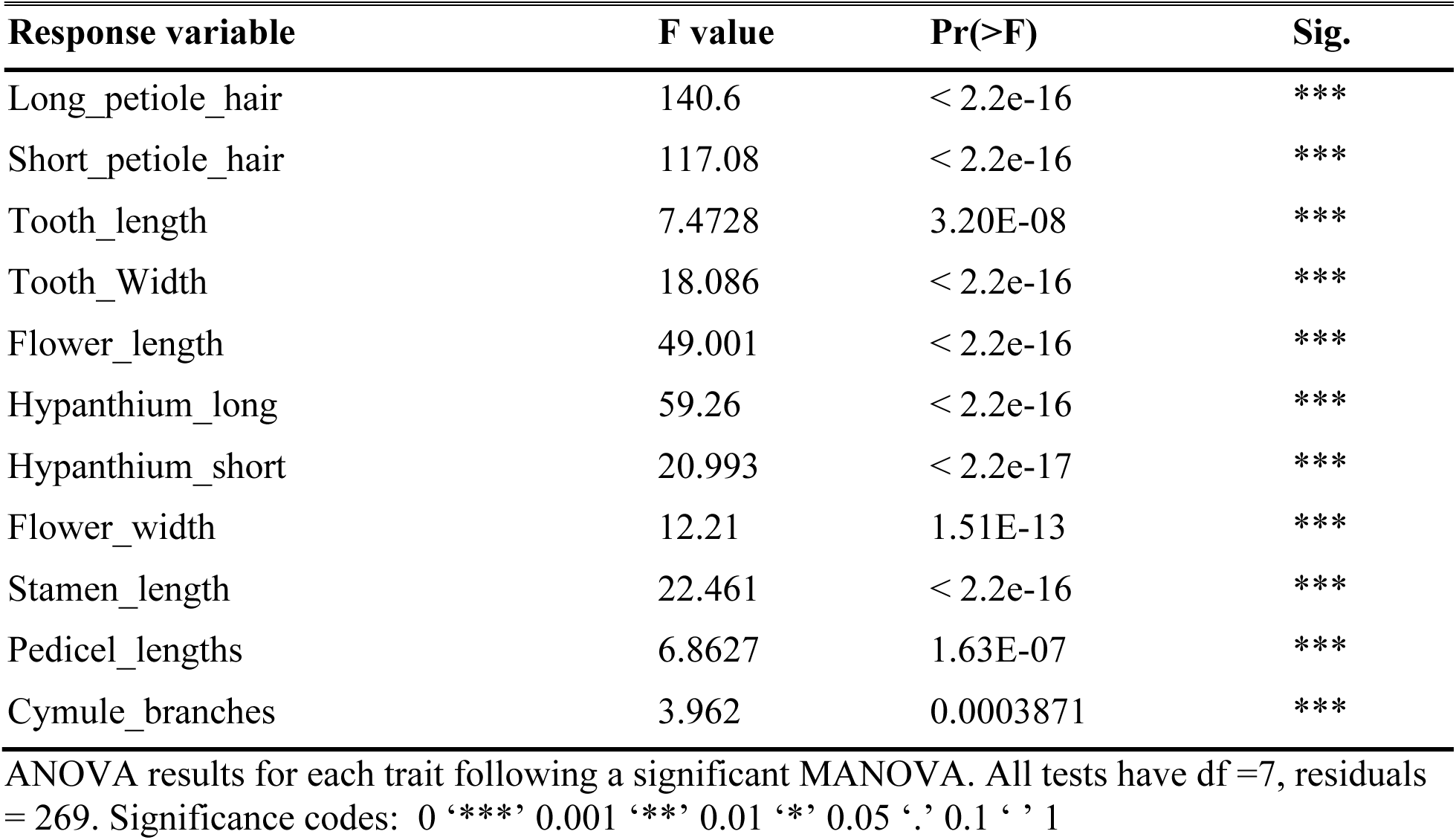
Univariate ANOVAs of morphological metrics.

**Table 2.**
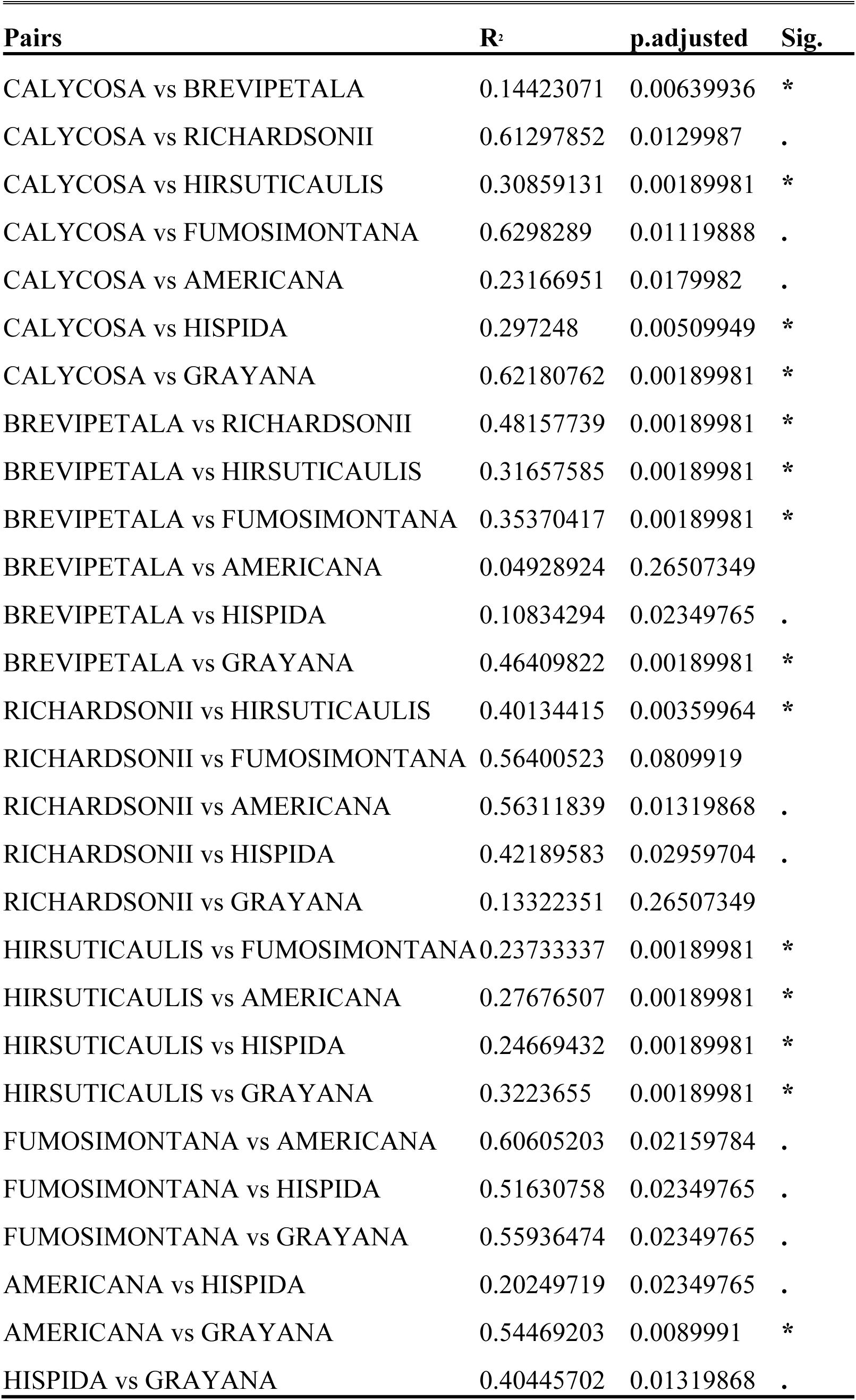
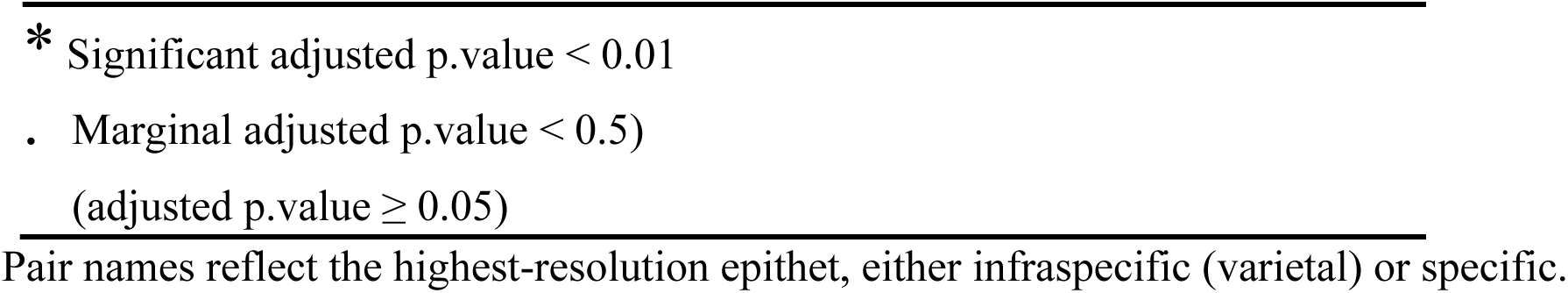
Pairwise comparisons of morphology via pairwise PERMANOVA (adonis).

| **Pairs** | **R<sup>2</sup>** | **p.adjusted** | **Sig.** |
| --- | --- | --- | --- |
| CALYCOSA vs BREVIPETALA | 0.14423071 | 0.00639936 | * |
| CALYCOSA vs RICHARDSONII | 0.61297852 | 0.0129987 | . |
| CALYCOSA vs HIRSUTICAULIS | 0.30859131 | 0.00189981 | * |
| CALYCOSA vs FUMOSIMONTANA | 0.6298289 | 0.01119888 | . |
| CALYCOSA vs AMERICANA | 0.23166951 | 0.0179982 | . |
| CALYCOSA vs HISPIDA | 0.297248 | 0.00509949 | * |
| CALYCOSA vs GRAYANA | 0.62180762 | 0.00189981 | * |
| BREVIPETALA vs RICHARDSONII | 0.48157739 | 0.00189981 | * |
| BREVIPETALA vs HIRSUTICAULIS | 0.31657585 | 0.00189981 | * |
| BREVIPETALA vs FUMOSIMONTANA | 0.35370417 | 0.00189981 | * |
| BREVIPETALA vs AMERICANA | 0.04928924 | 0.26507349 |  |
| BREVIPETALA vs HISPIDA | 0.10834294 | 0.02349765 | . |
| BREVIPETALA vs GRAYANA | 0.46409822 | 0.00189981 | * |
| RICHARDSONII vs HIRSUTICAULIS | 0.40134415 | 0.00359964 | * |
| RICHARDSONII vs FUMOSIMONTANA | 0.56400523 | 0.0809919 |  |
| RICHARDSONII vs AMERICANA | 0.56311839 | 0.01319868 | . |
| RICHARDSONII vs HISPIDA | 0.42189583 | 0.02959704 | . |
| RICHARDSONII vs GRAYANA | 0.13322351 | 0.26507349 |  |
| HIRSUTICAULIS vs FUMOSIMONTANA | 0.23733337 | 0.00189981 | * |
| HIRSUTICAULIS vs AMERICANA | 0.27676507 | 0.00189981 | * |
| HIRSUTICAULIS vs HISPIDA | 0.24669432 | 0.00189981 | * |
| HIRSUTICAULIS vs GRAYANA | 0.3223655 | 0.00189981 | * |
| FUMOSIMONTANA vs AMERICANA | 0.60605203 | 0.02159784 | . |
| FUMOSIMONTANA vs HISPIDA | 0.51630758 | 0.02349765 | . |
| FUMOSIMONTANA vs GRAYANA | 0.55936474 | 0.02349765 | . |
| AMERICANA vs HISPIDA | 0.20249719 | 0.02349765 | . |
| AMERICANA vs GRAYANA | 0.54469203 | 0.0089991 | * |
| HISPIDA vs GRAYANA | 0.40445702 | 0.01319868 | . |

\* Significant adjusted p.value < 0.01
. Marginal adjusted p.value < 0.5)
(adjusted p.value ≥ 0.05)
Pair names reflect the highest-resolution epithet, either infraspecific (varietal) or specific.

## Discussion

### A syngameon of five species

The phenophyletic view of species requires 1) a lineage *sensu* de Queiroz (1998) (i.e. an ancestral descent sequence of metapopulation[s]), to 2) be diagnosable through a cohesive phenotypic profile to be considered as a species. The use of lineages *sensu* de Queiroz (1998) is a pragmatic response to realworld geneflow found in some taxa that have inherently low barriers to genetic isolation; as is the case for *Heuchera*. More explicitly, this species concept enables us to delimit species when faced with a seemingly intractable syngameon as in *Heuchera* subsect. *Heuchera*. Wells (1984) too grapples with how to map species concepts on a syngameon; in her generic revision of *Heuchera* section *Heuchera*, *H. americana*, *H. caroliniana*, *H. longiflora*, *H. pubescens*, and *H. richardsonii* “constitute a closely related, natural group of species that are interfertile and vegetatively similar although distinctive in floral characters”. In particular, she noted that it is “necessary to recognize and distinguish between biological and taxonomic species in order to resolve the dilemma. In terms of breeding relationships, *H. americana*, *H. pubescens*, and *H. richardsonii* could be regarded as different ecotypes of one biological species. However, the magnitude of floral differences [including floral symmetry, normally quite conserved], the differences in geographical range, and the inability of the putative hybrids to migrate beyond the area of range overlap support the recognition of each of the three taxa as separate species within a syngameon (Grant, 1981). In *H. longiflora* and *H. caroliniana* the biological and taxonomic species concepts coincide reasonably well.”

While Rosendahl et al. (1936), the preceding authority, may have over-recognized species due to morphological differences, Wells (1979 & 1984) underrecognized species due to subjective categorical boundaries in morphology and a wish to impose the Biological Species Concept in an actively hybridizing radiation. Both monographic treatments lacked direct genetic data to test if lineage units correlate to morphology or even if gradients in morphology correspond at all to gradients in geneflow, with the caveat that (Wells, 1979) assessed hybrid morphology after controlled interspecific crosses and verified that the morphology of hybrid *H.* ×*hirsuticaulis* (hereafter at specific rank; see Taxonomic Treatment) and *H. americana* var. *hispida* are reproducible in synthetic hybrids. We chose to reexamine species limits for those putative taxa immediately related to *H. americana* and *H. richardsonii* using both genetic and morphological criteria. Similar to Wells (1984), our investigation finds a troubling combination of phenotypic cohesiveness among multiple potential taxa, combined with profuse gene flow creating seeming exceptions to every morphological rule. The same case Wells (1984) made for syngameons involving three of the species can now be extended to every member of *H.* subsect. *Heuchera* as most pointedly revealed by the fastSTRUCTURE analysis. Genetic patterns suggest strong but incomplete differentiation without complete reproductive isolation and in some cases morphological intergradation, a situation traditionally captured by botanists using subspecific rank (Rosendahl, 1949; Folk et al., 2018a). This no doubt is why Wells chose to eliminate several varieties previously recognized, since in her system varieties were used in the context of hybrids only.

We instead implement varieties as a designation for putative incipient speciation events that have not yet broken off into distinct groups (Folk et al., 2018a), aligning with the historical sense that the criteria of “good species” accumulate with time (Fig. 1 in de Queroz 2007; his “gray zone”). Whether intergrades are a failure of complete primary separation or secondary backcrossing is not directly relevant to decisions of rank (Freudenstein et al., 2017) although both fastSTRUCTURE and our “pure” ASTRAL analysis strongly suggests secondary gene flow among ancestrally distinct lineages given strong structure recovered in both. From this point forward, revised nomina and circumscriptions will be used as described hereafter. With these criteria in mind, three taxa among the focal *Heuchera americana* group meet the criteria to be recognized as species: *Heuchera americana*, *Heuchera richardsonii*, and *Heuchera fumosimontana* sp. nov. (see Taxonomic Treatment). The former two have been recognized for two centuries; the last is treated below. Two further taxa in the complex, *Heuchera×hirsuticaulis* and *Heuchera ×grayana* (see Taxonomic Treatment), are of hybrid origin based on morphology, artificial crosses, and phylogenomic data analyzed by multiple methods; interestingly they are also strictly homoploid (Folk and Freudenstein 2014 and references therein). Their recognition as taxa is justified by their stability over a wide geographic range and complete replacement of both parents across this entire range (Wells 1984; Folk et al. 2016; BIORXIV/2026/708067). Their recognition at specific rank is primarily based on the logic of applying hierarchical classification to networks. While the two hybrid taxa have a less discrete distinction from their parents and morphologically would merit varietal rank (Rosendahl et al., 1936; Wells, 1984), the Botanical Code strongly discourages the practice of treating hybrids under one of the parents at inferior rank (Madrid Code, H.5.1); under this recommendation these taxa are also treated at species rank.

It should be noted that for each species, we frequently found it necessary to use multiple analyses to individually assess distinctiveness and apply delimitation criteria. Together with limits in each analysis regarding decisiveness in individual cases, analytical limitations and assumptions of each analysis were used to weight decisions made when considering them as a whole. For example, fastSTRUCTURE is used to estimate individual ancestry proportions in the presence of gene flow while BPP is not meant for simultaneous species delimitation and gene flow estimation; see (Flouri et al., 2020). This can be problematic when trying to identify unique genetic lineages in a syngameon. As such, while STRUCTURE-like analysis are more commonly used to generate *a priori* units for subsequent species delimitation, no published model-based methods completely accommodate our taxa in scale or model complexity. We primarily deferred to fastSTRUCTURE for our unique lineage estimates and used BPP, ASTRAL, PCA, geography, and morphology to assess units recovered by fastSTRUCTURE. As an example, fastSTRUCTURE predicts two distinct pure lineages in *H.* ×*hirsuticaulis* (red and blue, Fig. 2), which have distinctive distributions (Fig. 9), but BPP lumps it as one species with *H. richardsonii*, a taxon from which it is easily diagnosed using morphology, and which has a distinct phylogenetic placement in ASTRAL. So our decision was primarily informed by fastSTRUCTURE and morphology over BPP in the distinctness of *H.* ×*hirsuticaulis* from *H. richardsonii*; but the analyses suggest that trusting fastSTRUCTURE at face value would oversplit *H.* ×*hirsuticaulis*. In our morphological diagnostics we weighted visually obvious traits as giving better diagnosability information than statistical analyses of morphometrics when clear separating traits were available. For example, *H. americana* var. *americana* and *H. americana* var. *brevipetala* had significant overlap in discriminant analyses (Figs. 6, 7) of morphological traits and a pairwise PERMANOVA (adonis) indicates a lack of significant difference between them (p = 0.48805119), but *H. americana* var. *americana* has a glabrous adaxial leaf lamina while *H. americana* var. *brevipetala* is diagnosible by a hirsutulous upper lamina over most of its wide range.

**Figure 9.**
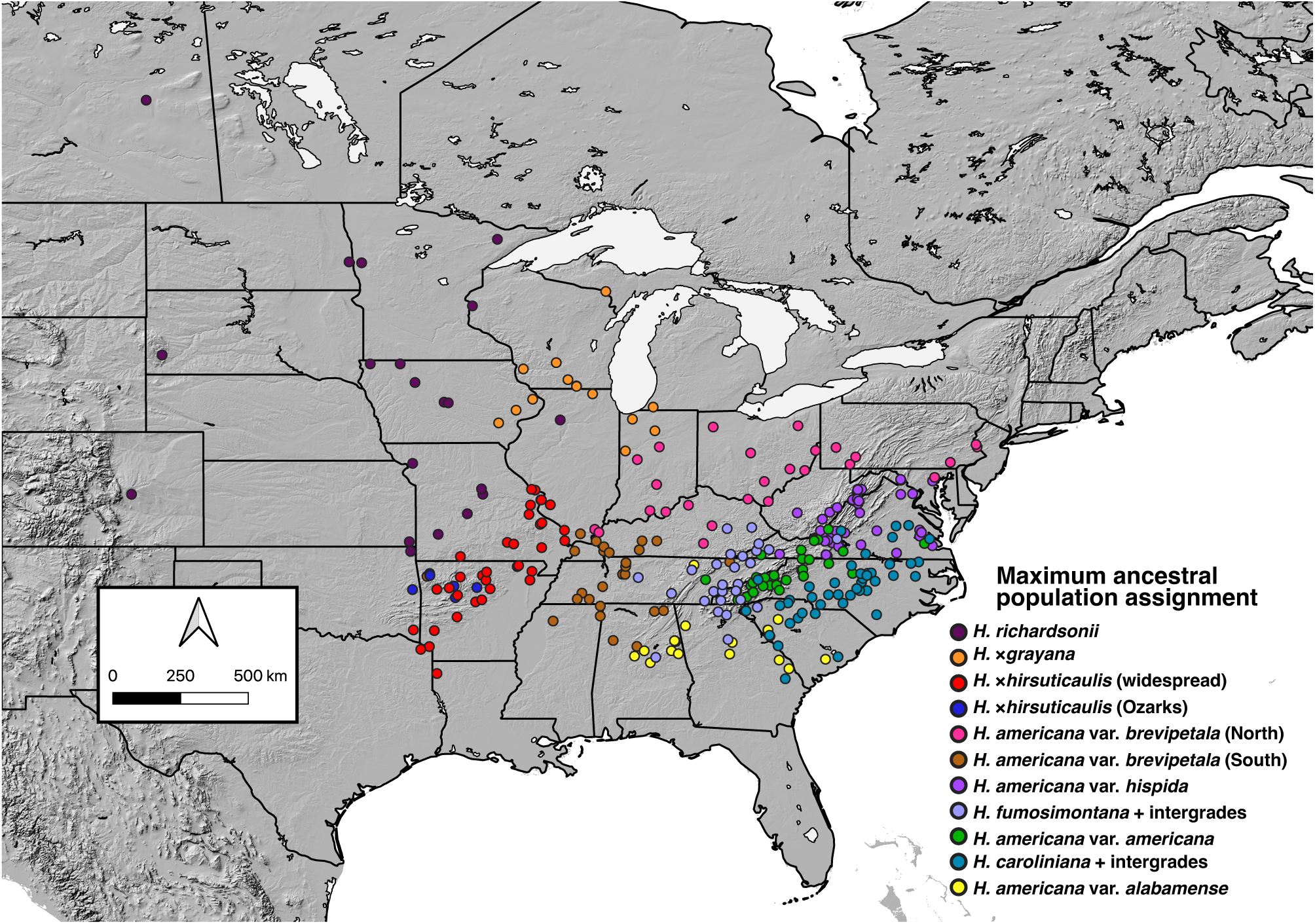
Ancestral population assignment map. Map of fastSTRUCTURE assignments (587 distinct samples shown at 309 distinct georeferenced localities). Colors and legend match Fig. 2. Descriptive titles of hypothetical ancestral populations accompany Discussion text.

### Species in the *Heuchera americana* group

Unsurprisingly, due to its obvious diagnosability in morphological features and geographic distance from the eastern hotspot of interspecific gene flow (BIORXIV/2026/708378), *Heuchera richardsonii* remains at the rank of species. These morphological traits include flowers much larger and zygomorphic than those of *H. americana* or any of its segregates, with larger length to width ratios than all but *H. ×grayana*, stamens predominantly included to level with sepal tip stamens, and hairy petioles (Fig. 10). This obvious morphological distinction is supported by univariate box plots (Fig. 8), a morphometric discriminant analysis (Fig. 6), and pairwise PERMANOVA (adonis) analysis (Table 2). The LDA (Fig. 6) showed clear morphological distinction with only some overlap with *H. ×grayana* and the pairwise PERMANOVA (adonis) analyses (Table 2) identified significant differences in multivariate morphology among all pairwise comparisons with *H. richardsonii* with the exception of *H. fumosimontana* and *H. ×grayana*. Genetic lineages were identified through cohesive clades in ASTRAL (Fig. 1), unique inferred ancestral population (purple) corresponding to taxon assignment in fastSTRUCTURE (Fig. 2), good clustering in an ordination of genetic variation via PCA (Fig. 3), strong (F_st_ 0.15-0.25) to very clear (F_st_ > 0.25) population structure among all pairwise comparisons of fixation index (Fig. 4) of taxa (except for with the hybrid species *H. ×hirsuticaulis* (F_st_ = 0.09) and *H. ×grayana* (F_st_ = 0.06)), BPP results corroborating fixation indices (Fig. 5), and a distinct geographic range only partially overlapping with *H. ×grayana* (Fig. 9). The higher degree of geneflow, indicated by fixation index, with the hybrid species is unsurprising in that *H. richardsonii* has long been thought to be a parental taxon.

**Figure 10.**
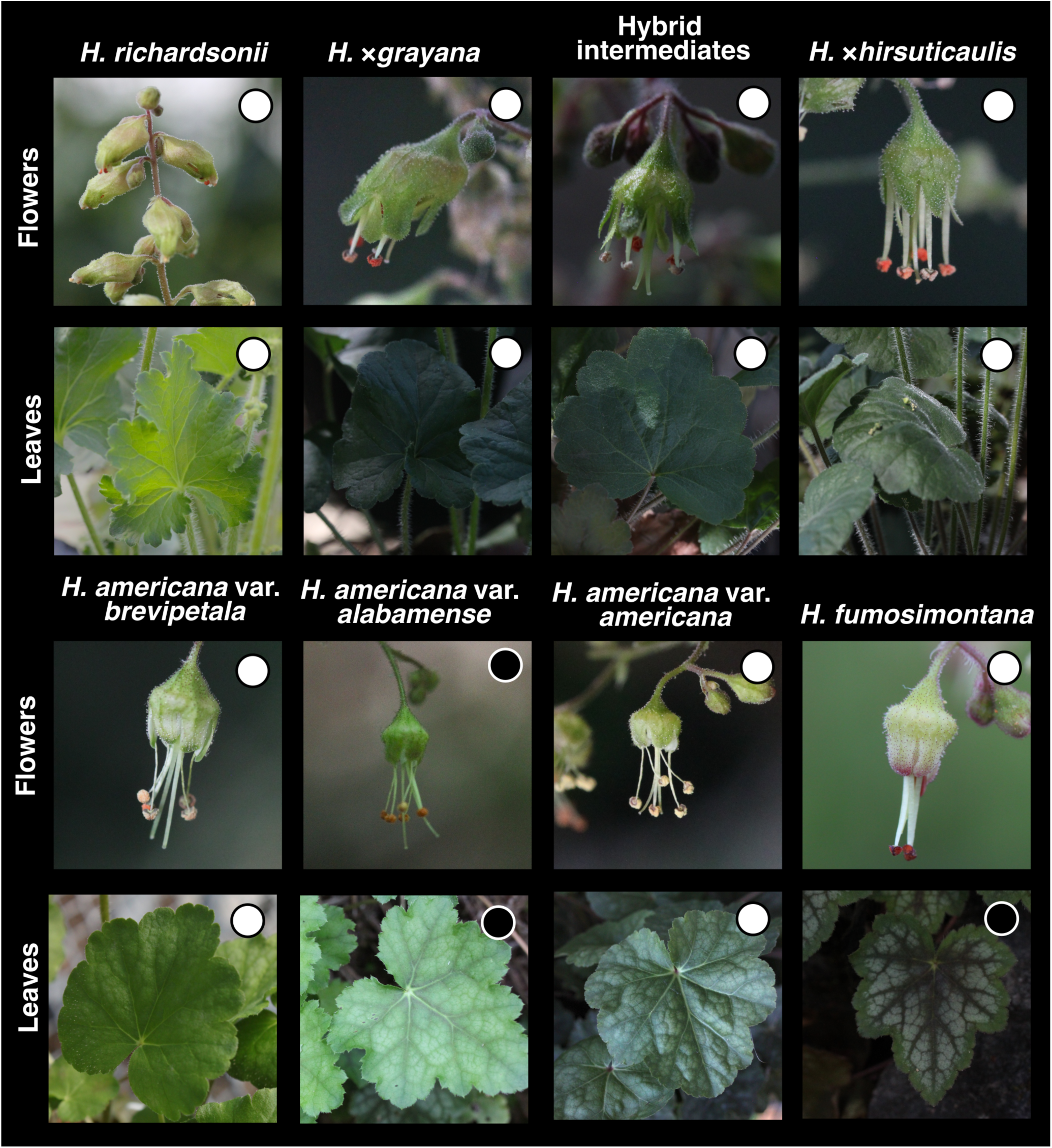
*Heuchera americana* group photoplate. Photoplate showing flowers and leaves of the taxa treated here. White dots indicate photos taken in common greenhouse conditions; black dots mark wild plant photos. Note differences in zygomorphy, exertion of the sexual parts, incurvation of free sepals, indumentum, floral attitude, inflorescence density, leaf lobe shape, and variegation

Concerning the remainder of the *H. americana* group (members of *Heuchera* subsect. *Heuchera* with exserted stamens), only three taxa were clearly distinguishable from *H. americana* as detectable lineages with diagnosable morphology: the hybrid species *Heuchera ×hirsuticaulis* and *Heuchera ×grayana*, and *Heuchera fumosimontana* (previously treated as a hairy-petioled aberrant *H. americana*). In Wells (1984), *H.* ×*hirsuticaulis* was treated as a variety of *H. americana* that had hirsute petioles, exserted stamens, and radially to zygomorphically symmetrical hypanthia. This description encompasses both *H. ×hirsuticaulis* and *Heuchera ×grayana* (more often treated in recent works as *H. richardsonii*). The hybrid zone that comprises these taxa and their parents, *Heuchera richardsonii* and *H. americana* var. *brevipetala*, has seen nearly a century of study with a variety of methodologies (Rosendahl et al., 1933, 1936; Wells, 1979; Folk et al., 2016); see also BIORXIV/2026/708067 and BIORXIV/2026/708378).

In accordance with their homoploid (Folk and Freudenstein, 2014) hybrid origin (Rosendahl et al., 1936; Wells, 1979; Folk et al., 2016), the hybrid taxa behave somewhat as varieties due to a gradient in distinct morphology characteristic of either parent and and distinct hybrid placements in phylogenetic analyses nearer to the geographically closer parent. Ordinated genetic analyses likewise plot these taxa as a smear between *H. richardsonii* and *H. americana* var. *brevipetala* (Fig. 3). However, while we expected these taxa to be genetically admixed, fastSTRUCTURE revealed many distinct “pure” individuals (taxon assignment > 90%) belonging to solely *H. ×hirsuticaulis* (red or blue in Fig. 2) or *H.* ×*grayana* (orange in Fig. 2). This finding was unexpected, and suggests that these taxa are the product of ancient hybridization events predating more recent gene flow, as long suspected (Rosendahl et al. 1936). Thus species rank is concordant with the lineage criterion. As an additional justification for the treatment at species rank, the *Madrid Code* (H.5.1) discourages the traditional practice of treating named hybrids at inferior rank under one of the parents. While BPP (Fig. 5) did lump *H.×grayana* with *H. ×hirsuticaulis* and *H. richardsonii*, this is interpreted to reflect an artifact gene flow, which the model as implemented does not accomodate; alternatively fastSTRUCTURE recovers these three taxa as entangled backcrosses deriving from four distinct but admixed ancestral populations and the “pure” ASTRAL topology recovers these taxa in separate clades (Fig. 1). Leaving aside statistically identified recent intergrades, the taxa considered here at species rank have consistent phenotypic diagnosability.

For *H. ×hirsuticaulis*, pairwise comparisons of fixation indices show moderate to strong differentiation in population structure (0.1 < F_st_ < 0.25) between all comparisons, except for with *H. ×grayana* (F_st_ = 0.06), *H. richardsonii* (F_st_ = 0.09), and *H. americana* var. *brevipetala* (F_st_ = 0.09), for which there is moderate differentiation and some gene flow. This is expected as the low F_st_ reflects geographic adjacency and suspected parental taxa. Again, this was a pattern reflected in BPP (Fig. 5); however morphometric analyses support the distinction of *H. ×hirsuticaulis*. A discriminant analysis shows almost no overlap with any other taxa (Fig. 6) and a pairwise PERMANOVA (Table 2) shows significant differentiation from all taxa. As noted above, *Heuchera ×hirsuticaulis* contains ancestry from two unique ancestral populations, one covering the range of the taxon and a second restricted to the southern Ozarks (red and blue respectively in Fig. 9). *Heuchera ×hirsuticaulis* primarily differs from *H. ×grayana* in having smaller 2.34–5.3 [-6.61] mm flowers vs 4.17–9.81 mm, radially symmetrical hypanthia, more prominent stamen exsertion [1.94-] 2.21–5.11 [-5.22] mm vs 1.13–3.40 [-4.27] mm, frequently having adaxial laminar hairs, and lower overall floral length to width ratios (Fig. 8).

*Heuchera ×grayana* had a much higher degree of population structure across all pairwise comparisons (Fig. 4) and lesser admixture with other taxa when compared with *H. ×hirsuticaulis* (F_st_ = 0.06) and *H. richardsonii* (F_st_ = 0.06). Lower fixation indices between *H. ×grayana* with *H. ×hirsuticaulis* and *H. richardsonii* is corroborated by BPP (Fig. 5) and intuitive due to *H. ×grayana* residing geographically between them, in a diagonal band ranging from Oklahoma to Wisconsin and Illinois (Fig. 9). However, fastSTRUCTURE (Fig. 2) highlights pure lineages primarily belonging to *H. grayana* (orange) and that most admixture is with *H. richardsonii*. This is supported by ASTRAL (Fig. 1) with a well supported clade of *H. ×grayana* (LPP = 0.86) sister to *H. richardsonii*. In addition to diagnosable lineages, a pairwise PERMANOVA (Table 2) between *H. ×hirsuticaulis* and *H. ×grayana* supports significant (p = 0.00189981) morphological differentiation between them and a morphological discriminant analysis (Fig. 6) shows no overlap between them.

*Heuchera fumosimontana* could have understandably been mistaken for *H. ×hirsuticaulis* in that it has the same morphological characteristics as *H. americana*, with the exception of hairy petioles. However, *H. fumosimontana* is: 1) is restricted to the Tennessee side of the Smoky Mountains and the Ocoee River Gorge and (importantly) is far-disjunct geographically from the range of *H. ×hirsuticaulis*, 2) has a generally hirsute rather than villous indumentum on its petioles, and 3) morphologically differs in other respects such as its smaller flowers, lesser leaf dentation, and longer upper laminar hairs (Fig. 8). Pairwise comparisons of fixation index (Fig. 4) between *H. fumosimontana* and *H. ×hirsuticaulis* supports strong differentiation (0.15 < F_st_ < 0.25) while other pairwise comparisons suggest *H. fumosimontana* is least differentiated from geographically adjacent *H. americana* varieties (F_st_ < 0.1), discussed below, and the *H. longiflora* group (F_st_ < 0.125). A morphological discriminant analysis shows clustering and separation *H. fumosimontana*, and separation from other taxa, only slightly overlapping with *H. ×hirsuticaulis* and pairwise PERMANOVA comparisons show significant morphological difference from all taxa. *Heuchera fumosimontana* differs most from *H. ×hirsuticaulis* in smaller tooth lengths 0.88–3.73 mm vs 0.96–8.61 [-9.63] mm, always having adaxial laminar hairs rather than sometimes having them, slightly longer median stamen exsertion 4.05 mm vs 3.506 mm, and smaller length to width ratios (Fig. 8). We conclude that this lineage restricted to the Smokey Mountains and Ocoee Gorge is significantly distinct from all described taxa in lineage and morphology and merits species rank. Furthermore, *H. fumosimontana* is a narrowly endemic species and completely replaces all other taxa of *Heuchera* subsect. *Heuchera* in its small range, a typical pattern in the genus (Wells, 1984). For these reasons, *H. fumosimontana* is easily identified using a geographically informed key (below).

### *Heuchera americana* as a polytypic species

The remainder of taxa treated here as *Heuchera americana* are collectively paraphyletic (Fig. 1) with high gene flow (Figs. 2, 4) and some morphological overlap (Fig. 7). The species is diagnosable as having glabrous petioles (but see notes to *H. fumosimontana* in the Taxonomic Treatment, below), pendent flowers, actinomorphic or nearly actinomorphic hypanthia, and strongly exserted stamens (Figs. 8, 10). Morphotypes in this polymorphic species correspond well to geographic distributions and display distinct phylogenetic placement, and thus recognition of segregates relieves somewhat the paraphyly of the whole. Yet as noted by all previous workers (Rosendahl et al. 1936, Wells 1984), the morphotypes intergrade where they meet, likely as a result of secondary gene flow after divergence, and are best treated at subspecific rank. This is the most confounding region of the *H. americana* group because it likely captures most of the geneflow history dating from the Pleistocene to the present; see BIORXIV/2026/708378.

Incipient speciation and widespread gene flow lead to high morphological and genetic variation paired with frequent intermediates, rendering both lumping and splitting unsatisfying. In these instances, the varietal rank is a useful handle for acknowledging evolutionary history while also recognizing incomplete distinctness from other members of the species to which the variety belongs. By contrast, preferring the single rank of species could be seen as miscommunicating hierarchical distinctness, and a lumping decision results in unacceptable polyphyly (Freudenstein et al., 2016) given our other delimitation decisions.

Among the focal taxa of the *H. americana* group we find that *H. americana* var. *alabamense* (formerly *H. americana* var. *calycosa*; see Taxonomic Treatment), *H. americana* var. *brevipetala*, and *H. americana* var. *americana* are assigned varietal rank. Each of these has several significantly different morphological characteristics with exceptions, forming an overall notably different gestalt, and has a characteristic geographic distribution. Phylogenetic placements are also distinctive although phylogenetic exclusivity is only seen in *H. americana* var. *alabamense.* While our descriptions and key below reflect the most distinct and typical versions of each variety, geography and morphology need to be considered collectively when identifying these varieties; especially near borders of distributions or in the absence of genetic data.

A set of populations is phenotypically diagnosable over a wide range in the lower Midwest by its clear upper laminar hairs and longer median stamen lengths. It morphologically and genetically overlaps with other *Heuchera*, yet with essentially no geographic overlap (Fig. 9). This regional taxonomic unit appears incipient and is suitably treated at varietal rank as *Heuchera americana* var. *brevipetala* Rosend., Butters & Lakela. *Heuchera americana* var. *brevipetala* is the western-most *H. americana* variety; it is characteristic of the lower Midwest, ranging from Indiana and Pennsylvania to Arkansas, increasingly sporadic east to Mississippi, Tennessee, Ohio, Pennsylvania, and lower New England. Analyses decisively point to this taxon as the immediate parent of hybrids with *H. richardsonii.* Like *H. ×hirsuticaulis*, *H. americana* var. *brevipetala* contains ancestry from two unique ancestral populations, one northern in distribution and one southern in distribution with the dividing line from southern Illinois through the center of Kentucky (Fig. 1); these regions are not morphologically distinctive. *Heuchera americana* var. *brevipetala* has the longest median stamen lengths (3.85 mm) of *H. americana*, and differs from *H. americana* var. *americana* primarily in having upper laminar hairs up to 1.5 mm (Fig. 8). fastSTRUCTURE suggests that *H. americana* var. *brevipetala* has many pure individuals (pink and brown [north and south respectively] in Figs. 2, 9) but a majority of individuals are of mixed assignment or shared assignment with *Heuchera ×hirsuticaulis* (brown and red, Fig. 2) and *H. fumosimontana* (pale blue, Fig. 2). While *H. americana* var. *brevipetala* and *H. americana* var. *americana* are most similar in morphology and share a wide geographic border, their population genetic distinctness and lack of evident gene flow accords with distinct phylogenetic placement (Fig. 1). Pairwise comparisons of fixation index largely mirror those of *H. americana* var. *alabamense* (below) in all pairwise comparisons except for low fixation with respect to *H. americana* var. *alabamense* (F_st_ = 0.04). The molecular PCA (Fig. 3) is an ordinated reflection of this trend, showing higher genetic overlap with other varieties of *H. americana* and no overlap with *H. ×grayana,* and *H. richardsonii*. The ASTRAL phylogeny (Fig. 1), pairwise comparisons of fixation index (Fig. 4), PCA of genetic variation (Fig. 3), fastSTRUCTURE (Fig. 2), and BPP (Fig. 5) all support *H. americana* var. *brevipetala* to varying degrees.

The remaining members of *Heuchera americana* reside to the east of the range of *H. americana* var. *brevipetala* in the broader Appalachian complex, spanning from southeastern Pennsylvania down into Georgia and Mississippi, and from the central Cumberland district east to the Coastal Plain. *Heuchera americana* var. *alabamense*, the most southerly in the species, is restricted in range to primarily the Alabama Cumberland Escarpment, sporadically northward in the Tennessee Cumberland and eastward to Georgia and South Carolina in the general area of the Fall Line. This taxon generally has the smallest flowers (2.14 - 4.26 mm from base to sepal tip) and stamen lengths (2.09–4.28 mm exserted length) of *H. americana* (Fig. 8), albeit with overlap in size with other *H. americana* varieties. The terminal leaf lobe is also commonly elongated. While fastSTRUCTURE (Fig. 2) assigns a significant portion of *H. americana* var. *alabamense* as pure individuals (yellow) and *H. americana* var. *alabamense* is a well supported clade in ASTRAL (LPP = 1, Fig. 1), pairwise comparisons of fixation indices suggest it has indistinguishable to low population structure and high to moderate gene flow when compared with populations of *H. americana* var. *brevipetala*, *H. americana* var. *americana*, and *H. fumosimontana* (F_st_< 0.1), Fig. 4. This set of circumstances is unexpected in that *H. americana* var. *alabamense* is the first-diverging named taxon in *Heuchera* subsect. *Heuchera* (Fig. 1) but compatible with an ancestrally distinct lineage with a high degree of secondary gene flow. Additionally, discriminant analyses reflect a broad overlap of morphology between *H. americana* var. *alabamense*, *H. americana* var. *brevipetala*, and *H. americana* var. *americana* (Figs. 6, 7), though significant differences among all morphological pairwise comparisons of *H. americana* var. *alabamense* (Table 2) indicate a distinct morphospace for this variety. These circumstances militate against species rank and justify treatment at the varietal level.

The remaining *H. americana* populations centering on the Crystalline Appalachians and Piedmont are most similar to *H. americana* var. *brevipetala* in floral morphology, but are similar to *H. americana* var. *alabamense* in that they predominantly lack upper leaf laminar hairs (Fig. 8). A PERMANOVA found significant morphological difference among all pairwise comparisons with *H. americana* var. *americana* except for with *H. americana* var. *brevipetala*, indicating imperfect morphological distinction from *H. americana* var. *brevipetala*, which is not its closest relative. Likewise, discriminant analyses (Figs. 7, 8) show strong morphological overlap of *H. americana* var. *americana* with *H. americana* var. *brevipetala* and *H. americana* var. *alabamense*. ASTRAL (Fig. 1) shows *H. americana* var. *americana* as intermixed with other members of the *Heuchera* syngameon, but with two well supported subclades (LPP = 1 and LPP = 0.97 respectively). FastSTRUCTURE reveals the highest level of admixture in *H. americana* var. *americana* compared to all other taxa, with most individuals being of mixed assignment. *Heuchera americana* var. *americana* contains a unique ancestral population (green in Figs. 2, 9), and a morass of intergrades involving the geographically adjacent taxa *H. caroliniana* (cerulean blue, Figs. 2, 9) and *H. fumosimontana* (light blue, Figs. 2, 9). The former account for *H. americana* accessions from North Carolina that placed with *H. caroliniana* in the ASTRAL phylogeny (Fig. 1). Strangely, these specimens do not at all morphologically intergrade with either *H. caroliniana* (see also Wells 1984, as this was her main reasoning for species rank) or *H. fumosimontana* (see Taxonomic Treatment below). *Heuchera americana* var. *americana* is, in summary, morphologically similar to other adjacent taxa, shows the most profuse introgression, and is the least easily diagnosable of the *Heuchera americana* varieties except by exclusion. In the absence of sufficient data to further divide the variety, and partly in view of avoiding polyphyly, this taxon is recognized as comprising those taxa not assignable to the remaining varieties. The variety is diagnosable as lacking upper laminar hairs, sharing floral size and morphology with *H. americana* var. *brevipetala*, but with shorter median stamen lengths (mean ∼3.4 mm [Fig. 8]), and distribution along the Appalachian mountain range and its peripherals from southeastern Pennsylvania into northern Georgia; it is the only taxon in the complex encountered in the southern and central Blue Ridge and most of the Piedmont. *Heuchera americana* var. *hispida* is provisionally maintained as a fourth variety pending forthcoming work on this taxon, which appears to have complex ancestry.

### Taxonomic Synopsis of the *H. americana* group

#### Heuchera richardsonii

*Heuchera richardsonii* R.Br. TYPE: British N. Am. *Hooker s.n.* [Franklin’s Expedition] (holotype: CGE; isotype: GH photo!).

= *Heuchera richardsonii* var. *hispidior* Rosend., Butters & Lakela. TYPE: Minnesota: Ft. Snelling, June 14, 1931 O. Lakela 532. (holotype: MIN photo!; isotypes: GH photo!, NY photo!, OSC photo!).

= *Heuchera richardsonii* var. *typica* Rosend., Butters & Lakela nom. illeg.

#### Heuchera ×grayana

*Heuchera ×grayana* (Rosend., Butters & Lakela) Engle-Wrye & R.A. Folk, **comb. nov.** TYPE: Illinois: Augusta, May 1843, *Mead s.n.* (holotype: MO [barcode: MO-247815 photo!]; isotypes: WIS, NY [barcode: NY00186000 photo!]).

≡*Heuchera richardsonii* var. *grayana* Rosend., Butters & Lakela, Rhodora 35: 117 (1933).

=*Heuchera richardsonii* var. *affinis* Rosend., Butters & Lakela, Minnesota Stud. Pl. Sci. 2: 124 (1936).

*Note—Heuchera richardsonii* var. *affinis* was used by Rosendahl, Butters & Lakela for hybrid individuals that have more of the morphology of *H. ×hirsuticaulis*, where *H. richardsonii var. grayana* was used for hybrid individuals more like *H. richardsonii*. The morphological difference between these can be marked, but a clear break was not evident in our genetic data.

#### Heuchera ×hirsuticaulis

*Heuchera ×hirsuticaulis* (Wheelock) Rydb., Man. Fl. N. States [Britton], 841 (1901). TYPE: Missouri: Rocks S of St. Louis, May 1865, *Engelmann s.n.* (holotype: GH; isotype: NY).

≡*Heuchera hispida* var. *hirsuticaulis* Wheelock, Bull. Torrey Bot. Club 17(8): 199 (1890).

≡*Heuchera americana var. hirsuticaulis* (Wheelock) Rosend., Butters & Lakela, Minnesota Stud. Pl. Sci. 2: 60 (1936).

=*Heuchera americana* var. *interior* Rosend., Butters & Lakela, Minnesota Stud. Pl. Sci. 2: 60 (1936). TYPE: 1 mi. South of North Salem, Hendricks Co. C.C. Dean 10858

### Heuchera fumosimontana

*Heuchera fumosimontana* Engle-Wrye & R.A. Folk, **sp. nov.** TYPE: United States of America. Tennessee: Polk County, on the roadside cliff along Hwy 64 near a turnout in the Ocoee River Gorge east of Lake Ocoee, 35°05’58.6” N, 84°33’19.8” W. May 10, 2022. *N.J. Engle-Wrye & R.A. Folk A55.* (holotype: MISSA! [barcode: MISSA037100]; isotype: MISSA! [barcode: MISSA037099]).

### Diagnosis

A *Heuchera hirsuticaulis* indumento petiolorum hirsuto, dentatione foliorum reductiore, pilis laminae adaxialis longioribus differt; a *H. americana* petiolorum hirsuto differt; atque distributione ad Montes Fumosos et Fauces Ocoee tantum restricta, ab *H. hirsuticaulis* geographice disiuncta.

### Description

Subcaulescent from a single thick caudex. **Petioles** hirsute throughout, 5.8–17.9 cm long. **Leaf** blades 4.9–7.9 cm wide, 5.5–9.2 cm long, ovate, deeply cordate with a wide sinus, moderately divided into 5 crenate (Ocoee Gorge) to triangular (Smoky Mountains) lobes, terminal lobe sometimes elongate; teeth crenate to triangular and prominently mucronate; upper lamina sparsely to moderately pubescent, lower lamina hispid predominantly along the veins; margins glabrous to ciliate; lamina strongly variegated in Smoky Mountain material. **Inflorescences** thyroid, 61–85.9 cm interrupted, primary axis puberulent; cymules 4–12 flowered, 2.4–4.3 cm long (measured to base of most distant flower), lax, monochasial, either trichasial or compressed monochasial the proximal node. **Flowers** 1.72–3.35 mm wide (at the junction of the hypanthium and ovary), [2.67-] 3.32–3.91 mm long, pale green with green to dark reddish purple to nearly black sepal tips, becoming darker towards fruit, campanulate, outer surfaces (of sepal, hypanthium, and ovary) glandular-puberulent, the glands reddish; free hypanthia 0.76–1.75 mm long on the longest side, 0.51–1.35 mm long on the shortest side, thus scarcely zygomorphic; free sepals rounded; petals included to approximately level with sepal tips, white; stamens exserted [2.99-] 3.43–4.36 mm past sepal margins, filaments filiform, anthers orbicular; closed true styles subulate proximally, filiform distally, exserted 2.74–5.59 mm past sepal margins, stigmas inconspicuously capitate. **Capsules** 3.28–3.88 mm wide, 4.9–6 mm long, ovoid, exceeding the accrescent calyx. **Seeds** not seen.

### Phenology

*Heuchera fumosimontana* flowers in the spring.

### Geographic distribution

Ocoee River Gorge (Polk County, Tennessee) and the Great Smoky Mountains, so far only known on the Tennessee side (Sevier County). The two areas of occurrence, while genetically similar, are morphologically differentiated, with the Ocoee River Gorge material comprising large plants with plain green leaves and low round leaf lobation, and the Smokies material comprising smaller plants with strongly variegated leaves and obviously triangular leaf lobation.

### Etymology

This species is named for the Great Smoky Mountains, its primary area of occurrence.

## Note

This lineage requires a new name despite the long history of observations of hairy-petioled *H. americana* far from the range of *H. ×hirsuticaulis* (Wells 1984) that seemed unlikely to be *H. richardsonii-*derived hybrids. A single specimen in our dataset from York County, Virginia (accession E1514, specimen barcode NCU00181130) represents Fernald’s concept of *H. americana* var. *heteradenia* (Fernald, 1942), a name we have previously used for Ocoee Gorge material (Folk and Freudenstein, 2014). A Tidewater Virginia aberration characterized by shortly hirsute petioles, the hairs in populations corresponding to Fernald’s taxon are mostly disposed to the distal end of the petiole. Our fastSTRUCTURE analysis assigns a probability for this sample of < 0.0001 to the unique *H. fumosimontana* ancestral population, thus ruling out this assignment completely. A similar result was found for hairy material from Screven County, Georgia (E1541, specimen barcode NCU00092996, also reviewed by Wells) which together with the lack of geographic and ecoregional commonality with Smokies material definitively rules out any available *Heuchera* nomen for the Smoky Mountain and Ocoee Gorge material. The Tidewater Virginian material is better assigned to *H. americana* var. *brevipetala*; the Georgia material is *H. americana* var. *alabamense*. While the odd distribution of hairy petioles in the *H. americana* complex could be confounding for identification, it is likely under simple genetic control and the exceptions are very narrowly distributed (two counties in Virginia, one in Georgia). These taxa are allopatric with *H. fumosimontana* and otherwise differ morphologically, particularly in flower size, dentation of the leaves, and upper laminar hairs (Fig. 8).

## Additional materials cited

United States of America: **Tennessee:** Sevier County: *N.J. Engle-Wrye & R.A. Folk A62*, (MISSA; barcode: MISSA037106), *N.J. Engle-Wrye & R.A. Folk A63*, (MISSA; barcode: MISSA037105).

### Heuchera americana

*Heuchera americana* L. Sp. Pl. 1: 226 (1753). TYPE: U.S.A. *Clayton ?*. (BM, UNC).

*Heuchera americana* L. var. *americana*.

=*Heuchera americana* var. *typica* Rosendahl, Butters & Lakela. nom. illeg. Minnesota Stud. Pl. Sci. 2: 55 (1936).

=*Heuchera scapifera* Moench, Methodus (Moench) 674 (1794). TYPE: unknown.

=*Heuchera cortusa* Michx., Fl. Bor.-Amer. (Michaux) 1: 171 (1803). TYPE: “Illinoensi regione, Carolina,” (P).

=*Heuchera glauca* Rafinesque, Med. Fl. 1: 244 (1828). TYPE: unknown.

≡*Heuchera americana* var*. glauca* Rosend., Bot. Jahrb. Syst. 37(2, Beibl. 83): 79 (1905).

=*Heuchera curtisii* Torr. & A.Gray ex A.Gray, Amer. J. Sci. Arts 42: 15 (1842). TYPE: North Carolina: Rocks, Buncombe, June 1818, *Curtis s.n.* (NY).

=*Heuchera americana* var. *calycosa* (Small) Rosend., Butters & Lakela, Minnesota Stud. Pl. Sci. 2: 58 (1936). TYPE: Mts. of Georgia, *Chapman s.n.*(holotype NY [barcode: 186000 photo!]). v

=*Heuchera americana* var. *subtruncata* Fernald, Rhodora 44: 401, tab. 721, fig. 1 (1942). TYPE: Virginia: Henrico Co. Rich wooded slope by James River, W of Varina. 6 June 1940.

*Fernald & Long 13031* (holotype: GH; isotype: PH).

### Heuchera americana *var.* brevipetala

*Heuchera americana* var. *brevipetala* Rosendahl, Butters & Lakela. Minnesota Stud. Pl. Sci. 2: 57 (1936). TYPE: New Jersey: Burlington Co., Ellisdale, *Long 25689* (holotype: PH).

=*Heuchera americana* var. *brevipetala* f. *angustipetala* Rosendahl, Butters & Lakela. Minnesota Stud. Pl. Sci. 2: 58 (1936). TYPE: Pennsylvania: Elwyn, 8 June 1890, *Brinton s.n.* (holotype: MIN [barcode: 122764 photo!]).

=*Heuchera americana* var. *heteradenia* Fernald. Rhodora 44: 400. (1942). TYPE: Virginia: Isle of Wight Co. Seeping calcareous wooded bluffs by James River, W of old Fort Boykin. 14 and 16 June 1941, *Fernald & Long 13027* (holotype: GH [barcodes: GH00056305 photo!, GH00042731 photo!]; isotypes: MIN, PH [barcodes: PH00014439 photo!, PH00014440 photo!])

=*Heuchera roseola* Rydb. in Britton, Man. Fl. N. States [Britton] 481 (1901). TYPE: Pennsylvania: York Furnace. 29 May 1892. *Brinton s.n*. (NY).

*=Heuchera lancipetala* Rydb. in Britton, Man. Fl. N. States [Britton] 482 (1901). TYPE: Kentucky: *Short 1840* (NY).

### Heuchera americana *var.* hispida

*Heuchera americana* var. *hispida* (Pursh) E.F.Wells, Rhodora 81: 576 (1979). TYPE: Virginia: Craig Co., High mountains between Fincastle and the Sweet Springs and some other similar places, *F. Pursh s. n.*, (authentic specimen of Pursh, PH)

≡*Heuchera hispida* Pursh, Fl. Amer. Sept. (Pursh) 1: 188 (1813).

### Heuchera americana *var.* alabamense

*Heuchera americana* var. *alabamense* Engle-Wrye & R.A. Folk, **var. nov.** TYPE: Alabama: Cleburne County: W of Pine Glen Campground and FS Rd 500, along Shoal Creek and in small ravine S of creek and N of FS Rd 531, ca. 5 air mi N of Heflin. 1989-05-07. *S.L. Orzell & Steve E.L. Bridges 9505* (holotype NCU! [barcode: NCU00181138]; isotype: TEX!).

### Diagnosis

A *Heuchera americana* var. *brevipetala* et *H. americana* var. *americana* floribus minoribus, staminibus minus exsertis, pilis laminae adaxialis plerumque absentibus sed interdum praesentibus differt.

### Description

Subcaulescent from a single thick caudex. **Petioles g**labrous, 5.7–13.8 cm long. **Leaf** blades 2.9–6.4 cm wide, [3.2-] 4.5–6.3 [-73] cm long, ovate with the terminal lobe commonly elongate, deeply cordate with a wide sinus moderately divided into 5 crenate to triangular lobes; teeth crenate to triangular and prominently mucronate; upper lamina most frequently glabrous, with sessile glands, occasionally hirsutulous; lower lamina glabrous rarely pubescent along veins; margin glabrous, sometimes subciliate. **Inflorescence** thyroid, 48.2–96.2 cm, interrupted, primary axis glabrous; cymules 4–7 [-8] flowered, 2.2–4.2 cm long (measured to base of most distant flower), lax, monochasial, trichasial or compressed monochasial at the proximal node. **Flowers** 1.09–3.12 mm wide (at the junction of the hypanthium and ovary), 2.14–4.26 mm long, pale green becoming darker with tints of purple towards fruit, campanulate to turbinate, outer surface minutely sessile-glandular, the glands reddish; free hypanthia 0.49–1.65 mm long on the longest side, 0.47–1.61 mm long on the shortest side, thus very nearly actinomorphic; free sepals rounded, petals included; stamens exserted 2.09–4.28 mm past sepal margins, filaments filiform, anthers orbicular; closed true styles subulate proximally, filiform distally, exserted 2.05–4.28 mm past sepal margins, stigmas inconspicuously capitate. **Capsules** 2.64–4.07 [-4.86] mm wide, 3.91–5.24 [-6.58] mm long, ovoid, exceeding the accrescent calyx. **Seeds** not seen.

### Phenology

*Heuchera americana* var. *alabamense* flowers in the spring.

### Geographic distribution

Most prevalent in the Alabama Cumberland Escarpment, also sporadically seen in adjacent the Tennessee Cumberland and the Piedmont and Coastal Plain of Georgia and South Carolina in the vicinity of the Fall Line; absent from the Crystalline Appalachians.

### Etymology

This variety is named for the state of Alabama, which harbors the least introgressed populations.

## Note

This lineage corresponds in large part to Rosendahl et al.’s morphological concept of *Heuchera americana* var. *calycosa* (Rosendahl et al., 1936), comprising individuals in the southern range of *H. americana* with small flowers. However, the holotype collection (Chapman *s.n.*, barcode NY00186000) is from the Appalachians of northern Georgia and does not have particularly small flowers, indicating it is assignable to *H. americana* var. *americana*, leaving this lineage without a name.

## Additional materials cited

United States of America—**Alabama:** Cleburne, *Folk A1*, (MISSA; barcode: MISSA034877); Clay, *Folk A2*, (MISSA; barcode: MISSA034880); Coosa, *Folk A3*, (MISSA; barcode: MISSA034881); Shelby, *Bowers 13612*, (UNA; barcode: UNA00034487); Clay, *Bussey 614*, (NCU; barcode: NCU00181137); Randolph, *McVaugh 8610*, (TEX); **South Carolina:** Abbeville, *Radford 220808*, (NCU; barcode: NCU00180859); Edgefield, *Radford 22536*, (NCU; barcode: NCU00180866); McCormick, *Radford 22388*, (NCU; barcode: NCU00180851); Pickens, *Knox 100*, (NCU; barcode: NCU00180854); **Georgia:** Screven, *Parrish 40*, (NCU; barcode: NCU00092996); Screven, *Park s.n.*, (NCU; barcode: NCU00092992); Morgan, *Hill 349*, (NCU; barcode: NCU00181139); Burke, *Pyron 2492*, (TEX); Floyd, *Kral 63437*, (TEX).

### Dichotomous key for members belonging to the *H. americana* group

NB: Floral morphological traits should only be assessed when flowers are in full anthesis and not yet in fruit.

1. Large flowers (free hypanthia > 2 mm) appearing strongly zygomorphic (adaxial free hypanthium obviously longer than abaxial hypanthium) ……………….. 2
2. Stamens included to approximately level with the sepal tips, in life the flowers more or less closed at the distal end with sepal tips disposed inwards; broadly distributed from Minnesota west to Colorado, and from western Missouri north to as Fort St. John, British Columbia, Canada; disjunct in the Wisconsin Driftless Region *H. richardsonii* 2. Stamens obviously but moderately exserted past sepal margins [0.5-] 1.13–3.4 [-4.27] mm, in life the flowers open at the distal end with the sepal tips disposed forwards; distributed strictly to the southeast of the range of *H. richardsonii* in a band from northeastern Oklahoma to Wisconsin (excluding the Driftless) and Illinois ……………….. *H. ×grayana* Smaller (free hypanthia < 2mm) flowers appearing more actinomorphic than zygomorphic (adaxial and abaxial hypanthium of similar length) ……………….. 3
3. Stamens exserted ≤ 1.5 mm beyond sepal tips ……………….. *H. caroliniana* 3. Stamens exserted > 1.5 mm beyond sepal tips ……………….. 4
4. Petioles hairy ……………….. 5
5. Petioles hirsute (longest petiole hairs 0.98–2.76 mm), primary tooth of secondary lobes often wider than long (median length and width of ∼ 1.59 *×* 4.84 mm), endemic to the Tennessee side of Smoky Mountains and the Ocoee River Gorge ……………….. *H. fumosimontana* 5. Petioles villous (longest petiole hairs 0.61–2.76 [-3.54] mm), primary tooth of secondary lobes often longer than wide (median length and width of ∼ 4.34 *×* 1.41 mm), broadly distributed from the Ozarks to Indiana and extreme northwestern Ohio; absent in the Appalachian system ……………….. *H. ×hirsuticaulis* 4. Petioles glabrous (with sessile glands) ……………….. 6
6. Free hypanthium 1.5-1.9 mm long; petals purple to brick-red and dentate ……………….. *Heuchera americana* var. *hispida* 6. Hypanthium less than 1.5 mm long; petals green, white, or somewhat pink and entire or with minute teeth ……………….. 7
7. Leaf adaxial lamina hirsutulous, broadly distributed in the American Midwest and adjacent regions from Indiana to Pennsylvania, to Arkansas, Mississippi, and Tennessee……*H. americana* var. *brevipetala* 7. Leaf adaxial lamina glabrous (with sessile glands; sometimes hirsutulous in *H. americana* var. *calycosa*) ……………….. 8 8. Flowers larger (2.3–5 mm), broadly distributed from Georgia to southeastern Pennsylvania, most prevalent along the Appalachian Mountain range and its peripherals …. *H. americana* var. *americana* 8. Flowers very small (2.14–4.26 mm), distributed in the Alabama Cumberland Escarpment, more sporadically in adjacent Tennessee and along the Fall Line in Georgia and South Carolina……………………. *H. americana* var. *alabamense*

## Conclusions

We bring a degree of order to the *Heuchera americana* group, but *Heuchera* section *Heuchera* remains an unruly syngameon. Our work provides a path to using reciprocal illumination (Hennig, 1966) among multiple lines of evidence when inferring evolutionary histories in the presence of widespread gene flow. We use a phenophyletic view emphasizing evidence for lineage and phenotypic diagnosability to delimit field-identifiable units, resurrecting four taxa from half a century of synonymy (*H. grayana, H. hirsuticaulis, H. americana* var. *alabamense,* and *H. americana* var. *brevipetala*) and recognizing one new to science (*H. fumosimontana*). As part of the challenge of dissecting a syngameon is that it defies the hierarchical logic of standard Linnean systematics, our integration of phylogenetic and population genetic methods with morphometric analysis provides for field-identifiable taxa that improve on the dissatisfactory combination of intergradation with paraphyly and strong geographic-morphological units. Distinguishing two levels of diagnosability and lineage independence, we use both specific and subspecific rank as previously suggested (Folk et al. 2018) to best communicate distinct and apparently incipient taxa.

## Acknowledgements

This work was funded by NSF DEB-2337784. We would like to thank the curators at MEM, TEX, UNA, and NCU for allowing us to visit and destructively sample herbarium specimens. We also thank Sofia Nail and Hawinet Adugna for their assistance in taking morphological measurements.

