## supplement for "Species delimitation in an intractable syngameon: Bringing order to the polyphyletic *Heuchera americana* group"

### Supplemental Tables

**Table S1** | Accession table of *Heuchera* specimens used for phylogenetic and morphological analyses.

| Accession | Morphological assignment | A priori species | fastSTRUCTURE assignment | A posteriori species | Collector + number | Provider acronym | Provider identifier | Country | State | County | Latitude | Longitude |
|---|---|---|---|---|---|---|---|---|---|---|---|---|
| **A1-1** | *Heuchera americana var. americana* | CALYCOSA | calycosa_yellow | *Heuchera americana var. alabamense* | Folk A1 | MISSA | MISSA034877 | USA | AL | Cleburne | 33.725 | -85.600833 |
| **A1-2** | *Heuchera americana var. americana* | CALYCOSA | calycosa_yellow | *Heuchera americana var. alabamense* | Folk A1 | MISSA | MISSA034877 | USA | AL | Cleburne | 33.725 | -85.600833 |
| **A1-2** | *Heuchera americana var. americana* | CALYCOSA | calycosa_yellow | *Heuchera americana var. alabamense* | Folk A1 | MISSA | MISSA034877 | USA | AL | Cleburne | 33.725 | -85.600833 |
| **A1-3** | *Heuchera americana var. americana* | CALYCOSA | calycosa_yellow | *Heuchera americana var. alabamense* | Folk A1 | MISSA | MISSA034877 | USA | AL | Cleburne | 33.725 | -85.600833 |
| **A1-4** | *Heuchera americana var. americana* | CALYCOSA | calycosa_yellow | *Heuchera americana var. alabamense* | Folk A1 | MISSA | MISSA034877 | USA | AL | Cleburne | 33.725 | -85.600833 |

Table S1: Continued

| | | | | | | | | | | | | |
|---|---|---|---|---|---|---|---|---|---|---|---|---|
| **A10-1** | *Heuchera americana var. americana* | BREVIPETALA | brevipetala_pink | *Heuchera americana var. brevipetala* | Folk A10 | MISSA | MISSA034513 | USA | IN | Floyd | 38.2152778 | -85.906944 |
| **A10-2** | *Heuchera americana var. americana* | BREVIPETALA | brevipetala_pink | *Heuchera americana var. brevipetala* | Folk A10 | MISSA | MISSA034513 | USA | IN | Floyd | 38.2152778 | -85.906944 |
| **A10-3** | *Heuchera americana var. americana* | BREVIPETALA | brevipetala_pink | *Heuchera americana var. brevipetala* | Folk A10 | MISSA | MISSA034513 | USA | IN | Floyd | 38.2152778 | -85.906944 |
| **A10-4** | *Heuchera americana var. americana* | BREVIPETALA | brevipetala_pink | *Heuchera americana var. brevipetala* | Folk A10 | MISSA | MISSA034513 | USA | IN | Floyd | 38.2152778 | -85.906944 |
| **A10-5** | *Heuchera americana var. americana* | BREVIPETALA | brevipetala_pink | *Heuchera americana var. brevipetala* | Folk A10 | MISSA | MISSA034513 | USA | IN | Floyd | 38.2152778 | -85.906944 |
| **A10-6** | *Heuchera americana var. americana* | BREVIPETALA | brevipetala_pink | *Heuchera americana var. brevipetala* | Folk A10 | MISSA | MISSA034513 | USA | IN | Floyd | 38.2152778 | -85.906944 |
| **A10-7** | *Heuchera americana var. americana* | BREVIPETALA | brevipetala_pink | *Heuchera americana var. brevipetala* | Folk A10 | MISSA | MISSA034513 | USA | IN | Floyd | 38.2152778 | -85.906944 |
| **A10-8** | *Heuchera americana var. americana* | BREVIPETALA | brevipetala_pink | *Heuchera americana var. brevipetala* | Folk A10 | MISSA | MISSA034513 | USA | IN | Floyd | 38.2152778 | -85.906944 |

Table S1: Continued

| | | | | | | | | | | | |
|---|---|---|---|---|---|---|---|---|---|---|---|
| **A11-1** | *Heuchera americana var. americana* | BREVIPETALA | brevipetala_pink | *Heuchera americana var. brevipetala* | Folk A11 | No voucher | USA | IN | Crawford | 38.2591667 | -86.461389 |
| **A11-2** | *Heuchera americana var. americana* | BREVIPETALA | brevipetala_pink | *Heuchera americana var. brevipetala* | Folk A11 | No voucher | USA | IN | Crawford | 38.2591667 | -86.461389 |
| **A11-3** | *Heuchera americana var. americana* | BREVIPETALA | brevipetala_pink | *Heuchera americana var. brevipetala* | Folk A11 | No voucher | USA | IN | Crawford | 38.2591667 | -86.461389 |
| **A11-4** | *Heuchera americana var. americana* | BREVIPETALA | brevipetala_pink | *Heuchera americana var. brevipetala* | Folk A11 | No voucher | USA | IN | Crawford | 38.2591667 | -86.461389 |
| **A11-5** | *Heuchera americana var. americana* | BREVIPETALA | brevipetala_pink | *Heuchera americana var. brevipetala* | Folk A11 | No voucher | USA | IN | Crawford | 38.2591667 | -86.461389 |
| **A11-6** | *Heuchera americana var. americana* | BREVIPETALA | brevipetala_pink | *Heuchera americana var. brevipetala* | Folk A11 | No voucher | USA | IN | Crawford | 38.2591667 | -86.461389 |
| **A11-7** | *Heuchera americana var. americana* | BREVIPETALA | brevipetala_pink | *Heuchera americana var. brevipetala* | Folk A11 | No voucher | USA | IN | Crawford | 38.2591667 | -86.461389 |
| **A11-8** | *Heuchera americana var. americana* | BREVIPETALA | brevipetala_pink | *Heuchera americana var. brevipetala* | Folk A11 | No voucher | USA | IN | Crawford | 38.2591667 | -86.461389 |

Table S1: Continued

| | | | | | | | | | | | | |
|---|---|---|---|---|---|---|---|---|---|---|---|---|
| **A13-1** | *Heuchera americana var. hirsuticaulis* | HIRSUTICAULIS | brevipetala_pink | *Heuchera hirsuticaulis* | Folk A13 | MISSA | MISSA036288 | USA | IL | Saline | 37.60485 | -88.384667 |
| **A13-2** | *Heuchera americana var. hirsuticaulis* | HIRSUTICAULIS | brevipetala_pink | *Heuchera hirsuticaulis* | Folk A13 | MISSA | MISSA036288 | USA | IL | Saline | 37.60485 | -88.384667 |
| **A13-3** | *Heuchera americana var. hirsuticaulis* | HIRSUTICAULIS | brevipetala_pink | *Heuchera hirsuticaulis* | Folk A13 | MISSA | MISSA036288 | USA | IL | Saline | 37.60485 | -88.384667 |
| **A13-4** | *Heuchera americana var. hirsuticaulis* | HIRSUTICAULIS | brevipetala_pink | *Heuchera hirsuticaulis* | Folk A13 | MISSA | MISSA036288 | USA | IL | Saline | 37.60485 | -88.384667 |
| **A13-5** | *Heuchera americana var. hirsuticaulis* | HIRSUTICAULIS | brevipetala_pink | *Heuchera hirsuticaulis* | Folk A13 | MISSA | MISSA036288 | USA | IL | Saline | 37.60485 | -88.384667 |
| **A13-6** | *Heuchera americana var. hirsuticaulis* | HIRSUTICAULIS | brevipetala_pink | *Heuchera hirsuticaulis* | Folk A13 | MISSA | MISSA036288 | USA | IL | Saline | 37.60485 | -88.384667 |
| **A13-7** | *Heuchera americana var. hirsuticaulis* | HIRSUTICAULIS | brevipetala_pink | *Heuchera hirsuticaulis* | Folk A13 | MISSA | MISSA036288 | USA | IL | Saline | 37.60485 | -88.384667 |
| **A13-8** | *Heuchera americana var. hirsuticaulis* | HIRSUTICAULIS | brevipetala_pink | *Heuchera hirsuticaulis* | Folk A13 | MISSA | MISSA036288 | USA | IL | Saline | 37.60485 | -88.384667 |

Table S1: Continued

| | | | | | | | | | | | | |
|---|---|---|---|---|---|---|---|---|---|---|---|---|
| **A13-9** | *Heuchera americana var. hirsuticaulis* | HIRSUTICAULIS | brevipetala_pink | *Heuchera hirsuticaulis* | Folk A13 | MISSA | MISSA036288 | USA | IL | Saline | 37.60485 | -88.384667 |
| **A14** | *Heuchera americana var. hirsuticaulis* | HIRSUTICAULIS | hirsuticaulis_red | *Heuchera hirsuticaulis* | Folk A14 | MISSA | MISSA036286 | USA | IL | Jersey | 38.9730556 | -90.464444 |
| **A15-10** | *Heuchera richardsonii* | RICHARDSONII | hirsuticaulis_red | *Heuchera grayana* | Folk A15 | MISSA | MISSA036290 | USA | MO | Camden | 38.1491167 | -92.825311 |
| **A15-11** | *Heuchera richardsonii* | RICHARDSONII | hirsuticaulis_red | *Heuchera grayana* | Folk A15 | MISSA | MISSA036290 | USA | MO | Camden | 38.1491167 | -92.825311 |
| **A15-12** | *Heuchera richardsonii* | RICHARDSONII | hirsuticaulis_red | *Heuchera grayana* | Folk A15 | MISSA | MISSA036290 | USA | MO | Camden | 38.1491167 | -92.825311 |
| **A15-13** | *Heuchera richardsonii* | RICHARDSONII | hirsuticaulis_red | *Heuchera grayana* | Folk A15 | MISSA | MISSA036290 | USA | MO | Camden | 38.1491167 | -92.825311 |
| **A15-15** | *Heuchera richardsonii* | RICHARDSONII | richardsonii | *Heuchera grayana* | Folk A15 | MISSA | MISSA036290 | USA | MO | Camden | 38.1491167 | -92.825311 |
| **A15-2** | *Heuchera richardsonii* | RICHARDSONII | hirsuticaulis_red | *Heuchera grayana* | Folk A15 | MISSA | MISSA036290 | USA | MO | Camden | 38.1491167 | -92.825311 |
| **A15-3** | *Heuchera richardsonii* | RICHARDSONII | hirsuticaulis_red | *Heuchera grayana* | Folk A15 | MISSA | MISSA036290 | USA | MO | Camden | 38.1491167 | -92.825311 |
| **A15-4** | *Heuchera richardsonii* | RICHARDSONII | hirsuticaulis_red | *Heuchera grayana* | Folk A15 | MISSA | MISSA036290 | USA | MO | Camden | 38.1491167 | -92.825311 |
| **A15-5** | *Heuchera richardsonii* | RICHARDSONII | hirsuticaulis_red | *Heuchera grayana* | Folk A15 | MISSA | MISSA036290 | USA | MO | Camden | 38.1491167 | -92.825311 |

Table S1: Continued

| | | | | | | | | | | | | |
|---|---|---|---|---|---|---|---|---|---|---|---|---|
| **A15-6** | *Heuchera richardsonii* | RICHARDSONII | richardsonii | *Heuchera grayana* | Folk A15 | MISSA | MISSA036290 | USA | MO | Camden | 38.1491167 | -92.825311 |
| **A15-7** | *Heuchera richardsonii* | RICHARDSONII | richardsonii | *Heuchera grayana* | Folk A15 | MISSA | MISSA036290 | USA | MO | Camden | 38.1491167 | -92.825311 |
| **A15-8** | *Heuchera richardsonii* | RICHARDSONII | richardsonii | *Heuchera grayana* | Folk A15 | MISSA | MISSA036290 | USA | MO | Camden | 38.1491167 | -92.825311 |
| **A15-9** | *Heuchera richardsonii* | RICHARDSONII | hirsuticaulis_red | *Heuchera grayana* | Folk A15 | MISSA | MISSA036290 | USA | MO | Camden | 38.1491167 | -92.825311 |
| **A16-1** | *Heuchera richardsonii* | GRAYANA | grayana_orange | *Heuchera grayana* | Folk A16 | MISSA | MISSA036549 | USA | WS | Sauk | 43.4175 | -89.726944 |
| **A16-10** | *Heuchera richardsonii* | GRAYANA | grayana_orange | *Heuchera grayana* | Folk A16 | MISSA | MISSA036549 | USA | WS | Sauk | 43.4175 | -89.726944 |
| **A16-11** | *Heuchera richardsonii* | GRAYANA | grayana_orange | *Heuchera grayana* | Folk A16 | MISSA | MISSA036549 | USA | WS | Sauk | 43.4175 | -89.726944 |
| **A16-2** | *Heuchera richardsonii* | GRAYANA | grayana_orange | *Heuchera grayana* | Folk A16 | MISSA | MISSA036549 | USA | WS | Sauk | 43.4175 | -89.726944 |
| **A16-3** | *Heuchera richardsonii* | GRAYANA | grayana_orange | *Heuchera grayana* | Folk A16 | MISSA | MISSA036549 | USA | WS | Sauk | 43.4175 | -89.726944 |
| **A16-4** | *Heuchera richardsonii* | GRAYANA | grayana_orange | *Heuchera grayana* | Folk A16 | MISSA | MISSA036549 | USA | WS | Sauk | 43.4175 | -89.726944 |
| **A16-6** | *Heuchera richardsonii* | GRAYANA | grayana_orange | *Heuchera grayana* | Folk A16 | MISSA | MISSA036549 | USA | WS | Sauk | 43.4175 | -89.726944 |
| **A16-8** | *Heuchera richardsonii* | GRAYANA | grayana_orange | *Heuchera grayana* | Folk A16 | MISSA | MISSA036549 | USA | WS | Sauk | 43.4175 | -89.726944 |

Table S1: Continued

| | | | | | | | | | | | | |
|---|---|---|---|---|---|---|---|---|---|---|---|---|
| **A16-9** | *Heuchera richardsonii* | GRAYANA | grayana_orange | *Heuchera grayana* | Folk A16 | MISSA | MISSA036549 | USA | WS | Sauk | 43.4175 | -89.726944 |
| **A16-9** | *Heuchera richardsonii* | GRAYANA | grayana_orange | *Heuchera grayana* | Folk A16 | MISSA | MISSA036549 | USA | WS | Sauk | 43.4175 | -89.726944 |
| **A17-1** | *Heuchera richardsonii* | RICHARDSONII | richardsonii | *Heuchera richardsonii* | Folk A17 | MISSA | MISSA036545 | USA | WS | Polk | 45.3975 | -92.648056 |
| **A17-2** | *Heuchera richardsonii* | RICHARDSONII | richardsonii | *Heuchera richardsonii* | Folk A17 | MISSA | MISSA036545 | USA | WS | Polk | 45.3975 | -92.648056 |
| **A17-3** | *Heuchera richardsonii* | RICHARDSONII | richardsonii | *Heuchera richardsonii* | Folk A17 | MISSA | MISSA036545 | USA | WS | Polk | 45.3975 | -92.648056 |
| **A17-4** | *Heuchera richardsonii* | RICHARDSONII | richardsonii | *Heuchera richardsonii* | Folk A17 | MISSA | MISSA036545 | USA | WS | Polk | 45.3975 | -92.648056 |
| **A17-5** | *Heuchera richardsonii* | RICHARDSONII | richardsonii | *Heuchera richardsonii* | Folk A17 | MISSA | MISSA036545 | USA | WS | Polk | 45.3975 | -92.648056 |
| **A17-6** | *Heuchera richardsonii* | RICHARDSONII | richardsonii | *Heuchera richardsonii* | Folk A17 | MISSA | MISSA036545 | USA | WS | Polk | 45.3975 | -92.648056 |
| **A17-7** | *Heuchera richardsonii* | RICHARDSONII | richardsonii | *Heuchera richardsonii* | Folk A17 | MISSA | MISSA036545 | USA | WS | Polk | 45.3975 | -92.648056 |
| **A17-8** | *Heuchera richardsonii* | RICHARDSONII | richardsonii | *Heuchera richardsonii* | Folk A17 | MISSA | MISSA036545 | USA | WS | Polk | 45.3975 | -92.648056 |
| **A17-9** | *Heuchera richardsonii* | RICHARDSONII | richardsonii | *Heuchera richardsonii* | Folk A17 | MISSA | MISSA036545 | USA | WS | Polk | 45.3975 | -92.648056 |

Table S1: Continued

| | | | | | | | | | | | | |
|---|---|---|---|---|---|---|---|---|---|---|---|---|
| **A2-3** | *Heuchera americana var. americana* | CALYCOSA | calycosa_yellow | *Heuchera americana var. alabamense* | Folk A2 | MISSA | MISSA034880 | USA | AL | Clay | 33.3705556 | -85.713056 |
| **A2-3** | *Heuchera americana var. americana* | CALYCOSA | calycosa_yellow | *Heuchera americana var. alabamense* | Folk A2 | MISSA | MISSA034880 | USA | AL | Clay | 33.3705556 | -85.713056 |
| **A2-4** | *Heuchera americana var. americana* | CALYCOSA | calycosa_yellow | *Heuchera americana var. alabamense* | Folk A2 | MISSA | MISSA034880 | USA | AL | Clay | 33.3705556 | -85.713056 |
| **A2-4** | *Heuchera americana var. americana* | CALYCOSA | calycosa_yellow | *Heuchera americana var. alabamense* | Folk A2 | MISSA | MISSA034880 | USA | AL | Clay | 33.3705556 | -85.713056 |
| **A21-1** | *Heuchera richardsonii* | RICHARDSONII | richardsonii | *Heuchera richardsonii* | Folk A21 | MISSA | MISSA036546 | USA | MN | Lake | 47.7205556 | -91.777778 |
| **A21-10** | *Heuchera richardsonii* | RICHARDSONII | richardsonii | *Heuchera richardsonii* | Folk A21 | MISSA | MISSA036546 | USA | MN | Lake | 47.7205556 | -91.777778 |
| **A21-11** | *Heuchera richardsonii* | RICHARDSONII | richardsonii | *Heuchera richardsonii* | Folk A21 | MISSA | MISSA036546 | USA | MN | Lake | 47.7205556 | -91.777778 |
| **A21-2** | *Heuchera richardsonii* | RICHARDSONII | richardsonii | *Heuchera richardsonii* | Folk A21 | MISSA | MISSA036546 | USA | MN | Lake | 47.7205556 | -91.777778 |
| **A21-3** | *Heuchera richardsonii* | RICHARDSONII | richardsonii | *Heuchera richardsonii* | Folk A21 | MISSA | MISSA036546 | USA | MN | Lake | 47.7205556 | -91.777778 |
| **A21-4** | *Heuchera richardsonii* | RICHARDSONII | richardsonii | *Heuchera richardsonii* | Folk A21 | MISSA | MISSA036546 | USA | MN | Lake | 47.7205556 | -91.777778 |

Table S1: Continued

| | | | | | | | | | | | | |
|---|---|---|---|---|---|---|---|---|---|---|---|---|
| **A21-5** | *Heuchera richardsonii* | RICHARDSONII | richardsonii | *Heuchera richardsonii* | Folk A21 | MISSA | MISSA036546 | USA | MN | Lake | 47.7205556 | -91.777778 |
| **A21-6** | *Heuchera richardsonii* | RICHARDSONII | richardsonii | *Heuchera richardsonii* | Folk A21 | MISSA | MISSA036546 | USA | MN | Lake | 47.7205556 | -91.777778 |
| **A21-7** | *Heuchera richardsonii* | RICHARDSONII | richardsonii | *Heuchera richardsonii* | Folk A21 | MISSA | MISSA036546 | USA | MN | Lake | 47.7205556 | -91.777778 |
| **A21-8** | *Heuchera richardsonii* | RICHARDSONII | richardsonii | *Heuchera richardsonii* | Folk A21 | MISSA | MISSA036546 | USA | MN | Lake | 47.7205556 | -91.777778 |
| **A21-9** | *Heuchera richardsonii* | RICHARDSONII | richardsonii | *Heuchera richardsonii* | Folk A21 | MISSA | MISSA036546 | USA | MN | Lake | 47.7205556 | -91.777778 |
| **A22-1** | *Heuchera glomerulata* | | not included | *Heuchera glomerulata* | Folk A22 | MISSA | MISSA034909 | USA | AZ | Graham | 32.6499094 | -109.81976 |
| **A26-1** | *Heuchera sanguinea* | | not included | | Folk A26 | MISSA | MISSA036297 | USA | AZ | Graham | 32.6359861 | -109.82353 |
| **A28-1** | *Heuchera americana var. americana* | CALYCOSA | brevipetala_brown | *Heuchera americana var. brevipetala* | Folk A28 | MISSA | MISSA036285 | USA | MS | Tishomingo | 34.58071 | -88.192537 |
| **A28-2** | *Heuchera americana var. americana* | CALYCOSA | brevipetala_brown | *Heuchera americana var. brevipetala* | Folk A28 | MISSA | MISSA036285 | USA | MS | Tishomingo | 34.58071 | -88.192537 |
| **A28-3** | *Heuchera americana var. americana* | CALYCOSA | brevipetala_brown | *Heuchera americana var. brevipetala* | Folk A28 | MISSA | MISSA036285 | USA | MS | Tishomingo | 34.58071 | -88.192537 |

Table S1: Continued

| | | | | | | | | | | | | |
|---|---|---|---|---|---|---|---|---|---|---|---|---|
| **A28-4** | *Heuchera americana var. americana* | CALYCOSA | brevipetala_brown | *Heuchera americana var. brevipetala* | Folk A28 | MISSA | MISSA036285 | USA | MS | Tishomingo | 34.58071 | -88.192537 |
| **A28-5** | *Heuchera americana var. americana* | CALYCOSA | brevipetala_brown | *Heuchera americana var. brevipetala* | Folk A28 | MISSA | MISSA036285 | USA | MS | Tishomingo | 34.58071 | -88.192537 |
| **A28-6** | *Heuchera americana var. americana* | CALYCOSA | brevipetala_brown | *Heuchera americana var. brevipetala* | Folk A28 | MISSA | MISSA036285 | USA | MS | Tishomingo | 34.58071 | -88.192537 |
| **A28-7** | *Heuchera americana var. americana* | CALYCOSA | brevipetala_brown | *Heuchera americana var. brevipetala* | Folk A28 | MISSA | MISSA036285 | USA | MS | Tishomingo | 34.58071 | -88.192537 |
| **A28-8** | *Heuchera americana var. americana* | CALYCOSA | brevipetala_brown | *Heuchera americana var. brevipetala* | Folk A28 | MISSA | MISSA036285 | USA | MS | Tishomingo | 34.58071 | -88.192537 |
| **A28-9** | *Heuchera americana var. americana* | CALYCOSA | brevipetala_brown | *Heuchera americana var. brevipetala* | Folk A28 | MISSA | MISSA036285 | USA | MS | Tishomingo | 34.58071 | -88.192537 |
| **A29-1** | *Heuchera americana var. americana* | CALYCOSA | brevipetala_brown | *Heuchera americana var. brevipetala* | Folk A29 | MISSA | MISSA036878 | USA | MS | Tishomingo | 34.9294925 | -88.191126 |
| **A29-10** | *Heuchera americana var. americana* | CALYCOSA | brevipetala_brown | *Heuchera americana var. brevipetala* | Folk A29 | MISSA | MISSA036902 | USA | MS | Tishomingo | 34.9294925 | -88.191126 |

Table S1: Continued

| | | | | | | | | | | | | |
|---|---|---|---|---|---|---|---|---|---|---|---|---|
| **A29-2** | *Heuchera americana var. americana* | CALYCOSA | brevipetala_brown | *Heuchera americana var. brevipetala* | Folk A29 | MISSA | MISSA036902 | USA | MS | Tishomingo | 34.9294925 | -88.191126 |
| **A29-3** | *Heuchera americana var. americana* | CALYCOSA | brevipetala_brown | *Heuchera americana var. brevipetala* | Folk A29 | MISSA | MISSA036902 | USA | MS | Tishomingo | 34.9294925 | -88.191126 |
| **A29-4** | *Heuchera americana var. americana* | CALYCOSA | brevipetala_brown | *Heuchera americana var. brevipetala* | Folk A29 | MISSA | MISSA036902 | USA | MS | Tishomingo | 34.9294925 | -88.191126 |
| **A29-5** | *Heuchera americana var. americana* | CALYCOSA | brevipetala_brown | *Heuchera americana var. brevipetala* | Folk A29 | MISSA | MISSA036902 | USA | MS | Tishomingo | 34.9294925 | -88.191126 |
| **A29-6** | *Heuchera americana var. americana* | CALYCOSA | brevipetala_brown | *Heuchera americana var. brevipetala* | Folk A29 | MISSA | MISSA036902 | USA | MS | Tishomingo | 34.9294925 | -88.191126 |
| **A29-7** | *Heuchera americana var. americana* | CALYCOSA | brevipetala_brown | *Heuchera americana var. brevipetala* | Folk A29 | MISSA | MISSA036902 | USA | MS | Tishomingo | 34.9294925 | -88.191126 |
| **A29-8** | *Heuchera americana var. americana* | CALYCOSA | brevipetala_brown | *Heuchera americana var. brevipetala* | Folk A29 | MISSA | MISSA036902 | USA | MS | Tishomingo | 34.9294925 | -88.191126 |
| **A29-9** | *Heuchera americana var. americana* | CALYCOSA | brevipetala_brown | *Heuchera americana var. brevipetala* | Folk A29 | MISSA | MISSA036902 | USA | MS | Tishomingo | 34.9294925 | -88.191126 |

Table S1: Continued

| | | | | | | | | | | | | |
|---|---|---|---|---|---|---|---|---|---|---|---|---|
| **A3-1** | *Heuchera americana var. americana* | CALYCOSA | calycosa_yellow | *Heuchera americana var. alabamense* | Folk A3 | MISSA | MISSA034881 | USA | AL | Coosa | 32.9541667 | -86.447222 |
| **A3-2** | *Heuchera americana var. americana* | CALYCOSA | calycosa_yellow | *Heuchera americana var. alabamense* | Folk A3 | MISSA | MISSA034881 | USA | AL | Coosa | 32.9541667 | -86.447222 |
| **A3-2** | *Heuchera americana var. americana* | CALYCOSA | calycosa_yellow | *Heuchera americana var. alabamense* | Folk A3 | MISSA | MISSA034881 | USA | AL | Coosa | 32.9541667 | -86.447222 |
| **A3-3** | *Heuchera americana var. americana* | CALYCOSA | calycosa_yellow | *Heuchera americana var. alabamense* | Folk A3 | MISSA | MISSA034881 | USA | AL | Coosa | 32.9541667 | -86.447222 |
| **A3-3** | *Heuchera americana var. americana* | CALYCOSA | calycosa_yellow | *Heuchera americana var. alabamense* | Folk A3 | MISSA | MISSA034881 | USA | AL | Coosa | 32.9541667 | -86.447222 |
| **A3-4** | *Heuchera americana var. americana* | CALYCOSA | calycosa_yellow | *Heuchera americana var. alabamense* | Folk A3 | MISSA | MISSA034881 | USA | AL | Coosa | 32.9541667 | -86.447222 |
| **A30-1** | *Heuchera villosa var. macrorhiza* | | | | Folk A30 | MISSA | MISSA036300 | USA | MS | Tishomingo | 34.930459 | -88.189257 |
| **A31** | *Heuchera americana var. hirsuticaulis* | HIRSUTICAULIS | brevipetala_brown | *Heuchera hirsuticaulis* | Engle-Wrye A31 | No voucher | | USA | MS | Panola | 34.408321 | -89.836198 |

Table S1: Continued

| | | | | | | | | | | | | |
|---|---|---|---|---|---|---|---|---|---|---|---|---|
| **A32-1** | *Heuchera americana var. hirsuticaulis* | HIRSUTICAULIS | brevipetala_brown | *Heuchera hirsuticaulis* | Engle-Wrye A32 | MISSA | MISSA036889 | USA | TN | Chester | 35.380564 | -88.832925 |
| **A32-10** | *Heuchera americana var. hirsuticaulis* | HIRSUTICAULIS | brevipetala_brown | *Heuchera hirsuticaulis* | Engle-Wrye A32 | MISSA | MISSA036889 | USA | TN | Chester | 35.380564 | -88.832925 |
| **A32-2** | *Heuchera americana var. hirsuticaulis* | HIRSUTICAULIS | brevipetala_brown | *Heuchera hirsuticaulis* | Engle-Wrye A32 | MISSA | MISSA036882 | USA | TN | Chester | 35.380564 | -88.832925 |
| **A32-3** | *Heuchera americana var. hirsuticaulis* | HIRSUTICAULIS | brevipetala_brown | *Heuchera hirsuticaulis* | Engle-Wrye A32 | MISSA | MISSA036889 | USA | TN | Chester | 35.380564 | -88.832925 |
| **A32-4** | *Heuchera americana var. hirsuticaulis* | HIRSUTICAULIS | brevipetala_brown | *Heuchera hirsuticaulis* | Engle-Wrye A32 | MISSA | MISSA036889 | USA | TN | Chester | 35.380564 | -88.832925 |
| **A32-5** | *Heuchera americana var. hirsuticaulis* | HIRSUTICAULIS | brevipetala_brown | *Heuchera hirsuticaulis* | Engle-Wrye A32 | MISSA | MISSA036889 | USA | TN | Chester | 35.380564 | -88.832925 |
| **A32-6** | *Heuchera americana var. hirsuticaulis* | HIRSUTICAULIS | brevipetala_brown | *Heuchera hirsuticaulis* | Engle-Wrye A32 | MISSA | MISSA036889 | USA | TN | Chester | 35.380564 | -88.832925 |
| **A32-7** | *Heuchera americana var. hirsuticaulis* | HIRSUTICAULIS | brevipetala_brown | *Heuchera hirsuticaulis* | Engle-Wrye A32 | MISSA | MISSA036889 | USA | TN | Chester | 35.380564 | -88.832925 |

Table S1: Continued

| | | | | | | | | | | | | |
|---|---|---|---|---|---|---|---|---|---|---|---|---|
| **A32-8** | *Heuchera americana var. hirsuticaulis* | HIRSUTICAULIS | brevipetala_brown | *Heuchera hirsuticaulis* | Engle-Wrye A32 | MISSA | MISSA036889 | USA | TN | Chester | 35.380564 | -88.832925 |
| **A32-9** | *Heuchera americana var. hirsuticaulis* | HIRSUTICAULIS | brevipetala_brown | *Heuchera hirsuticaulis* | Engle-Wrye A32 | MISSA | MISSA036889 | USA | TN | Chester | 35.380564 | -88.832925 |
| **A33-1** | *Heuchera americana var. americana* | BREVIPETALA | brevipetala_brown | *Heuchera americana var. brevipetala* | Engle-Wrye A33 | MISSA | MISSA036890 | USA | TN | Dickson | 36.036792 | -87.415364 |
| **A33-2** | *Heuchera americana var. americana* | BREVIPETALA | brevipetala_brown | *Heuchera americana var. brevipetala* | Engle-Wrye A33 | MISSA | MISSA036890 | USA | TN | Dickson | 36.036792 | -87.415364 |
| **A33-3** | *Heuchera americana var. americana* | BREVIPETALA | brevipetala_brown | *Heuchera americana var. brevipetala* | Engle-Wrye A33 | MISSA | MISSA036890 | USA | TN | Dickson | 36.036792 | -87.415364 |
| **A33-4** | *Heuchera americana var. americana* | BREVIPETALA | brevipetala_brown | *Heuchera americana var. brevipetala* | Engle-Wrye A33 | MISSA | MISSA036890 | USA | TN | Dickson | 36.036792 | -87.415364 |
| **A33-5** | *Heuchera americana var. americana* | BREVIPETALA | brevipetala_brown | *Heuchera americana var. brevipetala* | Engle-Wrye A33 | MISSA | MISSA036890 | USA | TN | Dickson | 36.036792 | -87.415364 |
| **A33-6** | *Heuchera americana var. americana* | BREVIPETALA | brevipetala_brown | *Heuchera americana var. brevipetala* | Engle-Wrye A33 | MISSA | MISSA036890 | USA | TN | Dickson | 36.036792 | -87.415364 |

Table S1: Continued

| | | | | | | | | | | | | |
|---|---|---|---|---|---|---|---|---|---|---|---|---|
| **A34-1** | *Heuchera americana* | BREVIPETALA | brevipetala_brown | *Heuchera americana var. brevipetala* | Engle-Wrye A34 | MISSA | MISSA036922 | USA | TN | Dickson | 36.036792 | -87.328511 |
| **A34-10** | *Heuchera americana var. americana* | BREVIPETALA | brevipetala_brown | *Heuchera americana var. brevipetala* | Engle-Wrye A34 | MISSA | MISSA036922 | USA | TN | Dickson | 36.036792 | -87.328511 |
| **A34-2** | *Heuchera americana* | BREVIPETALA | brevipetala_brown | *Heuchera americana var. brevipetala* | Engle-Wrye A34 | MISSA | MISSA036922 | USA | TN | Dickson | 36.036792 | -87.328511 |
| **A34-3** | *Heuchera americana var. americana* | BREVIPETALA | brevipetala_brown | *Heuchera americana var. brevipetala* | Engle-Wrye A34 | MISSA | MISSA036922 | USA | TN | Dickson | 36.036792 | -87.328511 |
| **A34-4** | *Heuchera americana var. americana* | BREVIPETALA | brevipetala_brown | *Heuchera americana var. brevipetala* | Engle-Wrye A34 | MISSA | MISSA036922 | USA | TN | Dickson | 36.036792 | -87.328511 |
| **A34-5** | *Heuchera americana var. americana* | BREVIPETALA | brevipetala_brown | *Heuchera americana var. brevipetala* | Engle-Wrye A34 | MISSA | MISSA036922 | USA | TN | Dickson | 36.036792 | -87.328511 |
| **A34-6** | *Heuchera americana var. americana* | BREVIPETALA | brevipetala_brown | *Heuchera americana var. brevipetala* | Engle-Wrye A34 | MISSA | MISSA036922 | USA | TN | Dickson | 36.036792 | -87.328511 |
| **A34-7** | *Heuchera americana var. americana* | BREVIPETALA | brevipetala_brown | *Heuchera americana var. brevipetala* | Engle-Wrye A34 | MISSA | MISSA036922 | USA | TN | Dickson | 36.036792 | -87.328511 |

Table S1: Continued

| | | | | | | | | | | | | |
|---|---|---|---|---|---|---|---|---|---|---|---|---|
| **A34-8** | *Heuchera americana var. americana* | BREVIPETALA | brevipetala_brown | *Heuchera americana var. brevipetala* | Engle-Wrye A34 | MISSA | MISSA036922 | USA | TN | Dickson | 36.036792 | -87.328511 |
| **A34-9** | *Heuchera americana var. americana* | BREVIPETALA | brevipetala_brown | *Heuchera americana var. brevipetala* | Engle-Wrye A34 | MISSA | MISSA036922 | USA | TN | Dickson | 36.036792 | -87.328511 |
| **A35-1** | *Heuchera americana var. americana* | BREVIPETALA | brevipetala_brown | *Heuchera americana var. brevipetala* | Engle-Wrye A35 | MISSA | MISSA036903 | USA | TN | Dickson | 36.311858 | -87.307933 |
| **A35-10** | *Heuchera americana var. americana* | BREVIPETALA | brevipetala_brown | *Heuchera americana var. brevipetala* | Engle-Wrye A35 | MISSA | MISSA036892 | USA | TN | Dickson | 36.311858 | -87.307933 |
| **A35-2** | *Heuchera americana var. americana* | BREVIPETALA | brevipetala_brown | *Heuchera americana var. brevipetala* | Engle-Wrye A35 | MISSA | MISSA036892 | USA | TN | Dickson | 36.311858 | -87.307933 |
| **A35-3** | *Heuchera americana var. americana* | BREVIPETALA | brevipetala_brown | *Heuchera americana var. brevipetala* | Engle-Wrye A35 | MISSA | MISSA036892 | USA | TN | Dickson | 36.311858 | -87.307933 |
| **A35-4** | *Heuchera americana var. americana* | BREVIPETALA | brevipetala_brown | *Heuchera americana var. brevipetala* | Engle-Wrye A35 | MISSA | MISSA036892 | USA | TN | Dickson | 36.311858 | -87.307933 |
| **A35-5** | *Heuchera americana var. americana* | BREVIPETALA | brevipetala_brown | *Heuchera americana var. brevipetala* | Engle-Wrye A35 | MISSA | MISSA036892 | USA | TN | Dickson | 36.311858 | -87.307933 |

Table S1: Continued

| | | | | | | | | | | | | |
|---|---|---|---|---|---|---|---|---|---|---|---|---|
| **A35-6** | *Heuchera americana var. americana* | BREVIPETALA | brevipetala_brown | *Heuchera americana var. brevipetala* | Engle-Wrye A35 | MISSA | MISSA036892 | USA | TN | Dickson | 36.311858 | -87.307933 |
| **A35-7** | *Heuchera americana var. americana* | BREVIPETALA | brevipetala_brown | *Heuchera americana var. brevipetala* | Engle-Wrye A35 | MISSA | MISSA036892 | USA | TN | Dickson | 36.311858 | -87.307933 |
| **A35-8** | *Heuchera americana var. americana* | BREVIPETALA | brevipetala_brown | *Heuchera americana var. brevipetala* | Engle-Wrye A35 | MISSA | MISSA036892 | USA | TN | Dickson | 36.311858 | -87.307933 |
| **A35-9** | *Heuchera americana var. americana* | BREVIPETALA | brevipetala_brown | *Heuchera americana var. brevipetala* | Engle-Wrye A35 | MISSA | MISSA036892 | USA | TN | Dickson | 36.311858 | -87.307933 |
| **A36** | *Heuchera americana var. americana* | BREVIPETALA | brevipetala_brown | *Heuchera americana var. brevipetala* | Engle-Wrye A36 | MISSA | MISSA036927 | USA | KY | Butler | 37.204169 | -86.736069 |
| **A37-5** | *Heuchera missouriensis* | | not included | *Heuchera missouriensis* | Engle-Wrye A37 | MISSA | MISSA036880 | USA | KY | Butler | 37.204169 | -86.736069 |
| **A38-1** | *Heuchera americana var. hirsuticaulis* | HIRSUTICAULIS | brevipetala_brown | *Heuchera hirsuticaulis* | Engle-Wrye A38 | MISSA | MISSA036910 | USA | KY | Logan | 36.888572 | -86.832992 |
| **A38-2** | *Heuchera americana var. hirsuticaulis* | HIRSUTICAULIS | brevipetala_brown | *Heuchera hirsuticaulis* | Engle-Wrye A38 | MISSA | MISSA036909 | USA | KY | Logan | 36.888572 | -86.832992 |

Table S1: Continued

| | | | | | | | | | | | | |
|---|---|---|---|---|---|---|---|---|---|---|---|---|
| **A39-1** | *Heuchera americana var. hirsuticaulis* | HIRSUTICAULIS | brevipetala_brown | *Heuchera hirsuticaulis* | Engle-Wrye A39 | MISSA | MISSA036894 | USA | KY | Trigg | 36.847206 | -88.072117 |
| **A39-10** | *Heuchera americana var. hirsuticaulis* | HIRSUTICAULIS | brevipetala_brown | *Heuchera hirsuticaulis* | Engle-Wrye A39 | MISSA | MISSA036894 | USA | KY | Trigg | 36.847206 | -88.072117 |
| **A39-11** | *Heuchera americana var. hirsuticaulis* | HIRSUTICAULIS | brevipetala_brown | *Heuchera hirsuticaulis* | Engle-Wrye A39 | MISSA | MISSA036894 | USA | KY | Trigg | 36.847206 | -88.072117 |
| **A39-2** | *Heuchera americana var. hirsuticaulis* | HIRSUTICAULIS | brevipetala_brown | *Heuchera hirsuticaulis* | Engle-Wrye A39 | MISSA | MISSA036894 | USA | KY | Trigg | 36.847206 | -88.072117 |
| **A39-3** | *Heuchera americana var. hirsuticaulis* | HIRSUTICAULIS | brevipetala_brown | *Heuchera hirsuticaulis* | Engle-Wrye A39 | MISSA | MISSA036894 | USA | KY | Trigg | 36.847206 | -88.072117 |
| **A39-4** | *Heuchera americana var. hirsuticaulis* | HIRSUTICAULIS | brevipetala_brown | *Heuchera hirsuticaulis* | Engle-Wrye A39 | MISSA | MISSA036894 | USA | KY | Trigg | 36.847206 | -88.072117 |
| **A39-5** | *Heuchera americana var. hirsuticaulis* | HIRSUTICAULIS | brevipetala_brown | *Heuchera hirsuticaulis* | Engle-Wrye A39 | MISSA | MISSA036894 | USA | KY | Trigg | 36.847206 | -88.072117 |
| **A39-6** | *Heuchera americana var. hirsuticaulis* | HIRSUTICAULIS | brevipetala_brown | *Heuchera hirsuticaulis* | Engle-Wrye A39 | MISSA | MISSA036894 | USA | KY | Trigg | 36.847206 | -88.072117 |

Table S1: Continued

| | | | | | | | | | | | | |
|---|---|---|---|---|---|---|---|---|---|---|---|---|
| **A39-7** | *Heuchera americana var. hirsuticaulis* | HIRSUTICAULIS | brevipetala_brown | *Heuchera hirsuticaulis* | Engle-Wrye A39 | MISSA | MISSA036894 | USA | KY | Trigg | 36.847206 | -88.072117 |
| **A39-8** | *Heuchera americana var. hirsuticaulis* | HIRSUTICAULIS | brevipetala_brown | *Heuchera hirsuticaulis* | Engle-Wrye A39 | MISSA | MISSA036894 | USA | KY | Trigg | 36.847206 | -88.072117 |
| **A39-9** | *Heuchera americana var. hirsuticaulis* | HIRSUTICAULIS | brevipetala_brown | *Heuchera hirsuticaulis* | Engle-Wrye A39 | MISSA | MISSA036894 | USA | KY | Trigg | 36.847206 | -88.072117 |
| **A4-1** | *Heuchera americana var. americana* | CALYCOSA | fumosimontana | *Heuchera americana var. americana* | Folk A4 | MISSA | MISSA034507 | USA | GA | Union | 34.7288889 | -84.082778 |
| **A4-2** | *Heuchera americana var. americana* | CALYCOSA | fumosimontana | *Heuchera americana var. americana* | Folk A4 | MISSA | MISSA034507 | USA | GA | Union | 34.7288889 | -84.082778 |
| **A4-2** | *Heuchera americana var. americana* | CALYCOSA | fumosimontana | *Heuchera americana var. americana* | Folk A4 | MISSA | MISSA034507 | USA | GA | Union | 34.7288889 | -84.082778 |
| **A4-3** | *Heuchera americana var. americana* | CALYCOSA | fumosimontana | *Heuchera americana var. americana* | Folk A4 | MISSA | MISSA034507 | USA | GA | Union | 34.7288889 | -84.082778 |
| **A4-4** | *Heuchera americana var. americana* | CALYCOSA | fumosimontana | *Heuchera americana var. americana* | Folk A4 | MISSA | MISSA034507 | USA | GA | Union | 34.7288889 | -84.082778 |

Table S1: Continued

| | | | | | | | | | | | | |
|---|---|---|---|---|---|---|---|---|---|---|---|---|
| **A4-5** | *Heuchera americana var. americana* | CALYCOSA | fumosimontana | *Heuchera americana var. americana* | Folk A4 | MISSA | MISSA034507 | USA | GA | Union | 34.7288889 | -84.082778 |
| **A40** | *Heuchera americana var. hirsuticaulis* | HIRSUTICAULIS | hirsuticaulis_red | *Heuchera hirsuticaulis* | Engle-Wrye A40 | MISSA | MISSA036919 | USA | IL | Union | 37.573553 | -89.439867 |
| **A40-2** | *Heuchera americana var. hirsuticaulis* | HIRSUTICAULIS | hirsuticaulis_red | *Heuchera hirsuticaulis* | Engle-Wrye A40 | MISSA | MISSA036893 | USA | IL | Union | 37.573553 | -89.439867 |
| **A40-3** | *Heuchera americana var. hirsuticaulis* | HIRSUTICAULIS | hirsuticaulis_red | *Heuchera hirsuticaulis* | Engle-Wrye A40 | MISSA | MISSA036919 | USA | IL | Union | 37.573553 | -89.439867 |
| **A41-1** | *Heuchera americana var. hirsuticaulis* | HIRSUTICAULIS | hirsuticaulis_red | *Heuchera hirsuticaulis* | Engle-Wrye A41 | MISSA | MISSA036925 | USA | MO | Wayne | 36.966256 | -90.234139 |
| **A41-2** | *Heuchera americana var. hirsuticaulis* | HIRSUTICAULIS | hirsuticaulis_red | *Heuchera hirsuticaulis* | Engle-Wrye A41 | MISSA | MISSA036888 | USA | MO | Wayne | 36.966256 | -90.234139 |
| **A41-3** | *Heuchera americana var. hirsuticaulis* | HIRSUTICAULIS | hirsuticaulis_red | *Heuchera hirsuticaulis* | Engle-Wrye A41 | MISSA | MISSA036911 | USA | MO | Wayne | 36.966256 | -90.234139 |
| **A42-1** | *Heuchera richardsonii* | RICHARDSONII | hirsuticaulis_red | *Heuchera grayana* | Engle-Wrye A42 | no voucher | | USA | MO | Jefferson | 38.454586 | -90.6237 |

Table S1: Continued

| | | | | | | | | | | | | |
|---|---|---|---|---|---|---|---|---|---|---|---|---|
| **A42-2** | *Heuchera richardsonii* | RICHARDSONII | hirsuticaulis_red | *Heuchera grayana* | Engle-Wrye A42 | no voucher | | USA | MO | Jefferson | 38.454586 | -90.6237 |
| **A42-3** | *Heuchera richardsonii* | RICHARDSONII | hirsuticaulis_red | *Heuchera grayana* | Engle-Wrye A42 | no voucher | | USA | MO | Jefferson | 38.454586 | -90.6237 |
| **A42-4** | *Heuchera richardsonii* | RICHARDSONII | hirsuticaulis_red | *Heuchera grayana* | Engle-Wrye A42 | no voucher | | USA | MO | Jefferson | 38.454586 | -90.6237 |
| **A43-1** | *Heuchera richardsonii* | RICHARDSONII | hirsuticaulis_red | *Heuchera grayana* | Engle-Wrye A43 | no voucher | | USA | IL | St. Louis | 38.630683 | -90.265731 |
| **A43-2** | *Heuchera richardsonii* | RICHARDSONII | hirsuticaulis_red | *Heuchera grayana* | Engle-Wrye A43 | no voucher | | USA | IL | St. Louis | 38.630683 | -90.265731 |
| **A43-3** | *Heuchera richardsonii* | RICHARDSONII | hirsuticaulis_red | *Heuchera grayana* | Engle-Wrye A43 | no voucher | | USA | IL | St. Louis | 38.630683 | -90.265731 |
| **A44-1** | *Heuchera richardsonii* | RICHARDSONII | hirsuticaulis_red | *Heuchera grayana* | Engle-Wrye A44 | MISSA | MISSA036881 | USA | MO | Boone | 38.830369 | -92.283972 |
| **A44-10** | *Heuchera richardsonii* | RICHARDSONII | hirsuticaulis_red | *Heuchera grayana* | Engle-Wrye A44 | MISSA | MISSA036881 | USA | MO | Boone | 38.830369 | -92.283972 |
| **A44-2** | *Heuchera richardsonii* | RICHARDSONII | hirsuticaulis_red | *Heuchera grayana* | Engle-Wrye A44 | MISSA | MISSA036881 | USA | MO | Boone | 38.830369 | -92.283972 |
| **A44-3** | *Heuchera richardsonii* | RICHARDSONII | hirsuticaulis_red | *Heuchera grayana* | Engle-Wrye A44 | MISSA | MISSA036881 | USA | MO | Boone | 38.830369 | -92.283972 |
| **A44-4** | *Heuchera richardsonii* | RICHARDSONII | hirsuticaulis_red | *Heuchera grayana* | Engle-Wrye A44 | MISSA | MISSA036881 | USA | MO | Boone | 38.830369 | -92.283972 |
| **A44-5** | *Heuchera richardsonii* | RICHARDSONII | hirsuticaulis_red | *Heuchera grayana* | Engle-Wrye A44 | MISSA | MISSA036881 | USA | MO | Boone | 38.830369 | -92.283972 |

Table S1: Continued

| | | | | | | | | | | | | |
|---|---|---|---|---|---|---|---|---|---|---|---|---|
| **A44-6** | *Heuchera richardsonii* | RICHARDSONII | richardsonii | *Heuchera richardsonii* | Engle-Wrye A44 | MISSA | MISSA036881 | USA | MO | Boone | 38.830369 | -92.283972 |
| **A44-7** | *Heuchera richardsonii* | RICHARDSONII | richardsonii | *Heuchera richardsonii* | Engle-Wrye A44 | MISSA | MISSA036881 | USA | MO | Boone | 38.830369 | -92.283972 |
| **A44-8** | *Heuchera richardsonii* | RICHARDSONII | richardsonii | *Heuchera richardsonii* | Engle-Wrye A44 | MISSA | MISSA036881 | USA | MO | Boone | 38.830369 | -92.283972 |
| **A44-9** | *Heuchera richardsonii* | RICHARDSONII | richardsonii | *Heuchera richardsonii* | Engle-Wrye A44 | MISSA | MISSA036881 | USA | MO | Boone | 38.830369 | -92.283972 |
| **A45-1** | *Heuchera americana var. hirsuticaulis* | HIRSUTICAULIS | hirsuticaulis_blue | *Heuchera hirsuticaulis* | Engle-Wrye A45 | MISSA | MISSA036897 | USA | AR | Washington | 36.065467 | -94.13855 |
| **A45-10** | *Heuchera americana var. hirsuticaulis* | HIRSUTICAULIS | hirsuticaulis_blue | *Heuchera hirsuticaulis* | Engle-Wrye A45 | MISSA | MISSA036897 | USA | AR | Washington | 36.065467 | -94.13855 |
| **A45-2** | *Heuchera americana var. hirsuticaulis* | HIRSUTICAULIS | hirsuticaulis_blue | *Heuchera hirsuticaulis* | Engle-Wrye A45 | MISSA | MISSA036897 | USA | AR | Washington | 36.065467 | -94.13855 |
| **A45-3** | *Heuchera americana var. hirsuticaulis* | HIRSUTICAULIS | hirsuticaulis_blue | *Heuchera hirsuticaulis* | Engle-Wrye A45 | MISSA | MISSA036897 | USA | AR | Washington | 36.065467 | -94.13855 |
| **A45-4** | *Heuchera americana var. hirsuticaulis* | HIRSUTICAULIS | hirsuticaulis_blue | *Heuchera hirsuticaulis* | Engle-Wrye A45 | MISSA | MISSA036897 | USA | AR | Washington | 36.065467 | -94.13855 |

Table S1: Continued

| | | | | | | | | | | | | |
|---|---|---|---|---|---|---|---|---|---|---|---|---|
| **A45-5** | *Heuchera americana var. hirsuticaulis* | HIRSUTICAULIS | hirsuticaulis_blue | *Heuchera hirsuticaulis* | Engle-Wrye A45 | MISSA | MISSA036897 | USA | AR | Washington | 36.065467 | -94.13855 |
| **A45-6** | *Heuchera americana var. hirsuticaulis* | HIRSUTICAULIS | hirsuticaulis_blue | *Heuchera hirsuticaulis* | Engle-Wrye A45 | MISSA | MISSA036897 | USA | AR | Washington | 36.065467 | -94.13855 |
| **A45-7** | *Heuchera americana var. hirsuticaulis* | HIRSUTICAULIS | hirsuticaulis_blue | *Heuchera hirsuticaulis* | Engle-Wrye A45 | MISSA | MISSA036897 | USA | AR | Washington | 36.065467 | -94.13855 |
| **A45-8** | *Heuchera americana var. hirsuticaulis* | HIRSUTICAULIS | hirsuticaulis_blue | *Heuchera hirsuticaulis* | Engle-Wrye A45 | MISSA | MISSA036897 | USA | AR | Washington | 36.065467 | -94.13855 |
| **A45-9** | *Heuchera americana var. hirsuticaulis* | HIRSUTICAULIS | hirsuticaulis_blue | *Heuchera hirsuticaulis* | Engle-Wrye A45 | MISSA | MISSA036897 | USA | AR | Washington | 36.065467 | -94.13855 |
| **A46-1** | *Heuchera americana var. hirsuticaulis* | HIRSUTICAULIS | hirsuticaulis_blue | *Heuchera hirsuticaulis* | Engle-Wrye A46 | MISSA | MISSA036920 | USA | AR | Washington | 35.994786 | -94.132508 |
| **A46-2** | *Heuchera americana var. hirsuticaulis* | HIRSUTICAULIS | hirsuticaulis_blue | *Heuchera hirsuticaulis* | Engle-Wrye A46 | MISSA | MISSA036924 | USA | AR | Washington | 35.994786 | -94.132508 |
| **A46-3** | *Heuchera americana var. hirsuticaulis* | HIRSUTICAULIS | hirsuticaulis_blue | *Heuchera hirsuticaulis* | Engle-Wrye A46 | MISSA | MISSA036899 | USA | AR | Washington | 35.994786 | -94.132508 |

Table S1: Continued

| | | | | | | | | | | | | |
|---|---|---|---|---|---|---|---|---|---|---|---|---|
| **A46-4** | *Heuchera americana var. hirsuticaulis* | HIRSUTICAULIS | hirsuticaulis_blue | *Heuchera hirsuticaulis* | Engle-Wrye A46 | MISSA | MISSA036899 | USA | AR | Washington | 35.994786 | -94.132508 |
| **A46-5** | *Heuchera americana var. hirsuticaulis* | HIRSUTICAULIS | hirsuticaulis_blue | *Heuchera hirsuticaulis* | Engle-Wrye A46 | MISSA | MISSA036899 | USA | AR | Washington | 35.994786 | -94.132508 |
| **A46-6** | *Heuchera americana var. hirsuticaulis* | HIRSUTICAULIS | hirsuticaulis_blue | *Heuchera hirsuticaulis* | Engle-Wrye A46 | MISSA | MISSA036899 | USA | AR | Washington | 35.994786 | -94.132508 |
| **A46-7** | *Heuchera americana var. hirsuticaulis* | HIRSUTICAULIS | hirsuticaulis_blue | *Heuchera hirsuticaulis* | Engle-Wrye A46 | MISSA | MISSA036899 | USA | AR | Washington | 35.994786 | -94.132508 |
| **A46-8** | *Heuchera americana var. hirsuticaulis* | HIRSUTICAULIS | hirsuticaulis_blue | *Heuchera hirsuticaulis* | Engle-Wrye A46 | MISSA | MISSA036899 | USA | AR | Washington | 35.994786 | -94.132508 |
| **A47-1** | *Heuchera americana var. hirsuticaulis* | HIRSUTICAULIS | hirsuticaulis_blue | *Heuchera hirsuticaulis* | Engle-Wrye A47 | MISSA | MISSA036915 | USA | AR | Washington | 35.996722 | -94.129067 |
| **A47-2** | *Heuchera americana var. hirsuticaulis* | HIRSUTICAULIS | hirsuticaulis_blue | *Heuchera hirsuticaulis* | Engle-Wrye A47 | MISSA | MISSA036914 | USA | AR | Washington | 35.996722 | -94.129067 |
| **A47-3** | *Heuchera americana var. hirsuticaulis* | HIRSUTICAULIS | hirsuticaulis_blue | *Heuchera hirsuticaulis* | Engle-Wrye A47 | MISSA | MISSA036896 | USA | AR | Washington | 35.996722 | -94.129067 |

Table S1: Continued

| | | | | | | | | | | | | |
|---|---|---|---|---|---|---|---|---|---|---|---|---|
| **A47-4** | *Heuchera americana var. hirsuticaulis* | HIRSUTICAULIS | hirsuticaulis_blue | *Heuchera hirsuticaulis* | Engle-Wrye A47 | MISSA | MISSA036915 | USA | AR | Washington | 35.996722 | -94.129067 |
| **A47-5** | *Heuchera americana var. hirsuticaulis* | HIRSUTICAULIS | hirsuticaulis_blue | *Heuchera hirsuticaulis* | Engle-Wrye A47 | MISSA | MISSA036918 | USA | AR | Washington | 35.996722 | -94.129067 |
| **A48** | *Heuchera americana var. hirsuticaulis* | HIRSUTICAULIS | hirsuticaulis_red | *Heuchera hirsuticaulis* | Engle-Wrye A48 | MISSA | MISSA036883 | USA | AR | Pope | 35.304633 | -93.165575 |
| **A49-1** | *Heuchera americana var. hirsuticaulis* | HIRSUTICAULIS | hirsuticaulis_red | *Heuchera hirsuticaulis* | Engle-Wrye A49 | MISSA | MISSA036921 | USA | AR | Faulkner | 35.074283 | -92.538006 |
| **A49-10** | *Heuchera americana var. hirsuticaulis* | HIRSUTICAULIS | hirsuticaulis_red | *Heuchera hirsuticaulis* | Engle-Wrye A49 | MISSA | MISSA036901 | USA | AR | Faulkner | 35.074283 | -92.538006 |
| **A49-11** | *Heuchera americana var. hirsuticaulis* | HIRSUTICAULIS | hirsuticaulis_red | *Heuchera hirsuticaulis* | Engle-Wrye A49 | MISSA | MISSA036895 | USA | AR | Faulkner | 35.074283 | -92.538006 |
| **A49-12** | *Heuchera americana var. hirsuticaulis* | HIRSUTICAULIS | hirsuticaulis_red | *Heuchera hirsuticaulis* | Engle-Wrye A49 | MISSA | MISSA036901 | USA | AR | Faulkner | 35.074283 | -92.538006 |
| **A49-2** | *Heuchera americana var. hirsuticaulis* | HIRSUTICAULIS | hirsuticaulis_red | *Heuchera hirsuticaulis* | Engle-Wrye A49 | MISSA | MISSA036901 | USA | AR | Faulkner | 35.074283 | -92.538006 |

Table S1: Continued

| | | | | | | | | | | | | |
|---|---|---|---|---|---|---|---|---|---|---|---|---|
| **A49-3** | *Heuchera americana var. hirsuticaulis* | HIRSUTICAULIS | hirsuticaulis_red | *Heuchera hirsuticaulis* | Engle-Wrye A49 | MISSA | MISSA036901 | USA | AR | Faulkner | 35.074283 | -92.538006 |
| **A49-4** | *Heuchera americana var. hirsuticaulis* | HIRSUTICAULIS | hirsuticaulis_red | *Heuchera hirsuticaulis* | Engle-Wrye A49 | MISSA | MISSA036901 | USA | AR | Faulkner | 35.074283 | -92.538006 |
| **A49-5** | *Heuchera americana var. hirsuticaulis* | HIRSUTICAULIS | hirsuticaulis_red | *Heuchera hirsuticaulis* | Engle-Wrye A49 | MISSA | MISSA036901 | USA | AR | Faulkner | 35.074283 | -92.538006 |
| **A49-6** | *Heuchera americana var. hirsuticaulis* | HIRSUTICAULIS | hirsuticaulis_red | *Heuchera hirsuticaulis* | Engle-Wrye A49 | MISSA | MISSA036901 | USA | AR | Faulkner | 35.074283 | -92.538006 |
| **A49-7** | *Heuchera americana var. hirsuticaulis* | HIRSUTICAULIS | hirsuticaulis_red | *Heuchera hirsuticaulis* | Engle-Wrye A49 | MISSA | MISSA036901 | USA | AR | Faulkner | 35.074283 | -92.538006 |
| **A49-8** | *Heuchera americana var. hirsuticaulis* | HIRSUTICAULIS | hirsuticaulis_red | *Heuchera hirsuticaulis* | Engle-Wrye A49 | MISSA | MISSA036901 | USA | AR | Faulkner | 35.074283 | -92.538006 |
| **A49-9** | *Heuchera americana var. hirsuticaulis* | HIRSUTICAULIS | hirsuticaulis_red | *Heuchera hirsuticaulis* | Engle-Wrye A49 | MISSA | MISSA036901 | USA | AR | Faulkner | 35.074283 | -92.538006 |
| **A5-1** | *Heuchera villosa var. villosa* | | | | Folk A5 | MISSA | MISSA034894 | USA | GA | Lumpkin | 34.6772222 | -84 |

Table S1: Continued

| | | | | | | | | | | | | |
|---|---|---|---|---|---|---|---|---|---|---|---|---|
| **A50-1** | *Heuchera caroliniana* | caroliniana | | *Heuchera caroliniana* | Folk A50 | MISSA | MISSA036277 | USA | SC | Pickens | 34.993338 | -80.081078 |
| **A50-2** | *Heuchera caroliniana* | caroliniana | | *Heuchera caroliniana* | Folk A50 | MISSA | MISSA036277 | USA | SC | Pickens | 34.993338 | -80.081078 |
| **A50-4** | *Heuchera caroliniana* | caroliniana | | *Heuchera caroliniana* | Folk A50 | MISSA | MISSA036277 | USA | SC | Pickens | 34.993338 | -80.081078 |
| **A53-1** | *Heuchera americana var. americana* | CALYCOSA | calycosa_yellow | *Heuchera americana var. alabamense* | Engle-Wrye A53 | no voucher | | USA | AL | Cleburne | 33.693964 | -85.55944 |
| **A53-10** | *Heuchera americana var. americana* | CALYCOSA | calycosa_yellow | *Heuchera americana var. alabamense* | Engle-Wrye A53 | no voucher | | USA | AL | Cleburne | 33.693964 | -85.55944 |
| **A53-2** | *Heuchera americana var. americana* | CALYCOSA | calycosa_yellow | *Heuchera americana var. alabamense* | Engle-Wrye A53 | no voucher | | USA | AL | Cleburne | 33.693964 | -85.55944 |
| **A53-3** | *Heuchera americana var. americana* | CALYCOSA | calycosa_yellow | *Heuchera americana var. alabamense* | Engle-Wrye A53 | no voucher | | USA | AL | Cleburne | 33.693964 | -85.55944 |

Table S1: Continued

| | | | | | | | | | | | |
|---|---|---|---|---|---|---|---|---|---|---|---|
| **A53-4** | *Heuchera americana var. americana* | CALYCOSA | calycosa_yellow | *Heuchera americana var. alabamense* | Engle-Wrye A53 | no voucher | USA | AL | Cleburne | 33.693964 | -85.55944 |
| **A53-5** | *Heuchera americana var. americana* | CALYCOSA | calycosa_yellow | *Heuchera americana var. alabamense* | Engle-Wrye A53 | no voucher | USA | AL | Cleburne | 33.693964 | -85.55944 |
| **A53-6** | *Heuchera americana var. americana* | CALYCOSA | calycosa_yellow | *Heuchera americana var. alabamense* | Engle-Wrye A53 | no voucher | USA | AL | Cleburne | 33.693964 | -85.55944 |
| **A53-7** | *Heuchera americana var. americana* | CALYCOSA | calycosa_yellow | *Heuchera americana var. alabamense* | Engle-Wrye A53 | no voucher | USA | AL | Cleburne | 33.693964 | -85.55944 |
| **A53-8** | *Heuchera americana var. americana* | CALYCOSA | calycosa_yellow | *Heuchera americana var. alabamense* | Engle-Wrye A53 | no voucher | USA | AL | Cleburne | 33.693964 | -85.55944 |
| **A53-9** | *Heuchera americana var. americana* | CALYCOSA | calycosa_yellow | *Heuchera americana var. alabamense* | Engle-Wrye A53 | no voucher | USA | AL | Cleburne | 33.693964 | -85.55944 |

Table S1: Continued

| | | | | | | | | | | | | |
|---|---|---|---|---|---|---|---|---|---|---|---|---|
| **A54-1** | *Heuchera americana var. americana* | CALYCOSA | calycosa_yellow | *Heuchera americana var. alabamense* | Engle-Wrye A54 | no voucher | | USA | AL | Shelby | 33.369119 | -86.659483 |
| **A54-2** | *Heuchera americana var. americana* | CALYCOSA | calycosa_yellow | *Heuchera americana var. alabamense* | Engle-Wrye A54 | no voucher | | USA | AL | Shelby | 33.369119 | -86.659483 |
| **A54-3** | *Heuchera americana var. americana* | CALYCOSA | calycosa_yellow | *Heuchera americana var. alabamense* | Engle-Wrye A54 | no voucher | | USA | AL | Shelby | 33.369119 | -86.659483 |
| **A55-1** | *Heuchera americana var. heteradenia* | HETERADENIA | fumosimontana | *Heuchera fumosimontana* | Folk A55 | MISSA | MISSA037100 | USA | TN | Polk | 35.0996176 | -84.555909 |
| **A55-2** | *Heuchera americana var. heteradenia* | HETERADENIA | fumosimontana | *Heuchera fumosimontana* | Folk A55 | MISSA | MISSA037100 | USA | TN | Polk | 35.0996176 | -84.555909 |
| **A55-3** | *Heuchera americana var. heteradenia* | HETERADENIA | fumosimontana | *Heuchera fumosimontana* | Folk A55 | MISSA | MISSA037100 | USA | TN | Polk | 35.0996176 | -84.555909 |
| **A55-4** | *Heuchera americana var. heteradenia* | HETERADENIA | fumosimontana | *Heuchera fumosimontana* | Folk A55 | MISSA | MISSA037100 | USA | TN | Polk | 35.0996176 | -84.555909 |

Table S1: Continued

| | | | | | | | | | | | | |
|---|---|---|---|---|---|---|---|---|---|---|---|---|
| **A55-5** | *Heuchera americana var. heteradenia* | HETERADENIA | fumosimontana | *Heuchera fumosimontana* | Folk A55 | MISSA | MISSA037100 | USA | TN | Polk | 35.0996176 | -84.555909 |
| **A55-6** | *Heuchera americana var. heteradenia* | HETERADENIA | fumosimontana | *Heuchera fumosimontana* | Folk A55 | MISSA | MISSA037100 | USA | TN | Polk | 35.0996176 | -84.555909 |
| **A56-1** | *Heuchera americana var. americana* | AMERICANA | fumosimontana | *Heuchera americana var. americana* | Engle-Wrye A56 | no voucher | | USA | NC | Cherokee | 35.1230527 | -83.989823 |
| **A56-2** | *Heuchera americana var. americana* | AMERICANA | fumosimontana | *Heuchera americana var. americana* | Engle-Wrye A56 | no voucher | | USA | NC | Cherokee | 35.1230527 | -83.989823 |
| **A56-3** | *Heuchera americana var. americana* | AMERICANA | fumosimontana | *Heuchera americana var. americana* | Engle-Wrye A56 | no voucher | | USA | NC | Cherokee | 35.1230527 | -83.989823 |
| **A56-4** | *Heuchera americana var. americana* | AMERICANA | fumosimontana | *Heuchera americana var. americana* | Engle-Wrye A56 | no voucher | | USA | NC | Cherokee | 35.1230527 | -83.989823 |

Table S1: Continued

| | | | | | | | | | | | |
|---|---|---|---|---|---|---|---|---|---|---|---|
| **A56-5** | *Heuchera americana var. americana* | AMERICANA | fumosimontana | *Heuchera americana var. americana* | Engle-Wrye A56 | no voucher | USA | NC | Cherokee | 35.1230527 | -83.989823 |
| **A56-6** | *Heuchera americana var. americana* | AMERICANA | fumosimontana | *Heuchera americana var. americana* | Engle-Wrye A56 | no voucher | USA | NC | Cherokee | 35.1230527 | -83.989823 |
| **A56-7** | *Heuchera americana var. americana* | AMERICANA | fumosimontana | *Heuchera americana var. americana* | Engle-Wrye A56 | no voucher | USA | NC | Cherokee | 35.1230527 | -83.989823 |
| **A57** | *Heuchera americana var. americana* | AMERICANA | americana_green | *Heuchera americana var. americana* | Engle-Wrye A57 | no voucher | USA | NC | Swain | 35.4648505 | -83.433531 |
| **A58-1** | *Heuchera americana var. americana* | AMERICANA | americana_green | *Heuchera americana var. americana* | Engle-Wrye A58 | no voucher | USA | NC | Jackson | 35.248278 | -83.086306 |
| **A58-2** | *Heuchera americana var. americana* | AMERICANA | americana_green | *Heuchera americana var. americana* | Engle-Wrye A58 | no voucher | USA | NC | Jackson | 35.248278 | -83.086306 |

| | | | | | | | | | | | | |
|---|---|---|---|---|---|---|---|---|---|---|---|---|
| **A58-3** | *Heuchera americana var. americana* | AMERICANA | americana_green | *Heuchera americana var. americana* | Engle-Wrye A58 | no voucher | | USA | NC | Jackson | 35.248278 | -83.086306 |
| **A58-4** | *Heuchera americana var. americana* | AMERICANA | americana_green | *Heuchera americana var. americana* | Engle-Wrye A58 | no voucher | | USA | NC | Jackson | 35.248278 | -83.086306 |
| **A59-1** | *Heuchera americana var. hispida* | AMERICANA | americana_green | *Heuchera americana var. hispida* | Engle-Wrye A59 | MISSA | MISSA037098 | USA | NC | McDowell | 35.735357 | -82.116329 |
| **A59-2** | *Heuchera americana var. hispida* | AMERICANA | americana_green | *Heuchera americana var. hispida* | Engle-Wrye A59 | MISSA | MISSA037098 | USA | NC | McDowell | 35.735357 | -82.116329 |
| **A59-3** | *Heuchera americana var. hispida* | AMERICANA | americana_green | *Heuchera americana var. hispida* | Engle-Wrye A59 | MISSA | MISSA037098 | USA | NC | McDowell | 35.735357 | -82.116329 |
| **A59-4** | *Heuchera americana var. hispida* | AMERICANA | americana_green | *Heuchera americana var. hispida* | Engle-Wrye A59 | MISSA | MISSA037098 | USA | NC | McDowell | 35.735357 | -82.116329 |
| **A59-5** | *Heuchera americana var. hispida* | AMERICANA | americana_green | *Heuchera americana var. hispida* | Engle-Wrye A59 | MISSA | MISSA037098 | USA | NC | McDowell | 35.735357 | -82.116329 |

Table S1: Continued

| | | | | | | | | | | | | |
|---|---|---|---|---|---|---|---|---|---|---|---|---|
| **A59-6** | *Heuchera americana var. hispida* | AMERICANA | americana_green | *Heuchera americana var. hispida* | Engle-Wrye A59 | MISSA | MISSA037098 | USA | NC | McDowell | 35.735357 | -82.116329 |
| **A59-7** | *Heuchera americana var. hispida* | AMERICANA | americana_green | *Heuchera americana var. hispida* | Engle-Wrye A59 | MISSA | MISSA037098 | USA | NC | McDowell | 35.735357 | -82.116329 |
| **A59-8** | *Heuchera americana var. hispida* | AMERICANA | americana_green | *Heuchera americana var. hispida* | Engle-Wrye A59 | MISSA | MISSA037098 | USA | NC | McDowell | 35.735357 | -82.116329 |
| **A59-9** | *Heuchera americana var. hispida* | AMERICANA | americana_green | *Heuchera americana var. hispida* | Engle-Wrye A59 | MISSA | MISSA037098 | USA | NC | McDowell | 35.735357 | -82.116329 |
| **A6-2** | *Heuchera americana var. americana* | CALYCOSA | fumosimontana | *Heuchera americana var. americana* | Folk A6 | MISSA | MISSA034509 | USA | GA | White | 34.6194444 | -83.792222 |
| **A6-3** | *Heuchera americana var. americana* | CALYCOSA | fumosimontana | *Heuchera americana var. americana* | Folk A6 | MISSA | MISSA034509 | USA | GA | White | 34.6194444 | -83.792222 |
| **A6-3** | *Heuchera americana var. americana* | CALYCOSA | fumosimontana | *Heuchera americana var. americana* | Folk A6 | MISSA | MISSA034509 | USA | GA | White | 34.6194444 | -83.792222 |
| **A60-1** | *Heuchera americana var. americana* | AMERICANA | americana_green | *Heuchera americana var. americana* | Engle-Wrye A60 | MISSA | no voucher | USA | NC | Alexander | 35.966166 | -81.114492 |

Table S1: Continued

| | | | | | | | | | | | | |
|---|---|---|---|---|---|---|---|---|---|---|---|---|
| **A60-2** | *Heuchera americana var. americana* | AMERICANA | americana_green | *Heuchera americana var. americana* | Engle-Wrye A60 | MISSA | no voucher | USA | NC | Alexander | 35.966166 | -81.114492 |
| **A60-3** | *Heuchera americana var. americana* | AMERICANA | americana_green | *Heuchera americana var. americana* | Engle-Wrye A60 | MISSA | no voucher | USA | NC | Alexander | 35.966166 | -81.114492 |
| **A60-4** | *Heuchera americana var. americana* | AMERICANA | americana_green | *Heuchera americana var. americana* | Engle-Wrye A60 | MISSA | no voucher | USA | NC | Alexander | 35.966166 | -81.114492 |
| **A62-1** | *Heuchera americana var. heteradenia* | HETERADENIA | fumosimontana | *Heuchera fumosimontana* | Engle-Wrye A62 | MISSA | MISSA037106 | USA | TN | Sevier | 35.695778 | -83.389556 |
| **A62-2** | *Heuchera americana var. heteradenia* | HETERADENIA | fumosimontana | *Heuchera fumosimontana* | Engle-Wrye A62 | MISSA | MISSA037106 | USA | TN | Sevier | 35.695778 | -83.389556 |
| **A62-3** | *Heuchera americana var. heteradenia* | HETERADENIA | fumosimontana | *Heuchera fumosimontana* | Engle-Wrye A62 | MISSA | MISSA037106 | USA | TN | Sevier | 35.695778 | -83.389556 |
| **A63-1** | *Heuchera americana var. heteradenia* | HETERADENIA | fumosimontana | *Heuchera fumosimontana* | Engle-Wrye A63 | MISSA | MISSA037105 | USA | TN | Sevier | 35.695778 | -83.389556 |
| **A63-10** | *Heuchera americana var. heteradenia* | HETERADENIA | fumosimontana | *Heuchera fumosimontana* | Engle-Wrye A63 | MISSA | MISSA037105 | USA | TN | Sevier | 35.695778 | -83.389556 |

Table S1: Continued

| | | | | | | | | | | | | |
|---|---|---|---|---|---|---|---|---|---|---|---|---|
| **A63-2** | *Heuchera americana var. heteradenia* | HETERADENIA | fumosimontana | *Heuchera fumosimontana* | Engle-Wrye A63 | MISSA | MISSA037105 | USA | TN | Sevier | 35.695778 | -83.389556 |
| **A63-3** | *Heuchera americana var. heteradenia* | HETERADENIA | fumosimontana | *Heuchera fumosimontana* | Engle-Wrye A63 | MISSA | MISSA037105 | USA | TN | Sevier | 35.695778 | -83.389556 |
| **A63-4** | *Heuchera americana var. heteradenia* | HETERADENIA | fumosimontana | *Heuchera fumosimontana* | Engle-Wrye A63 | MISSA | MISSA037105 | USA | TN | Sevier | 35.695778 | -83.389556 |
| **A63-5** | *Heuchera americana var. heteradenia* | HETERADENIA | fumosimontana | *Heuchera fumosimontana* | Engle-Wrye A63 | MISSA | MISSA037105 | USA | TN | Sevier | 35.695778 | -83.389556 |
| **A63-6** | *Heuchera americana var. heteradenia* | HETERADENIA | fumosimontana | *Heuchera fumosimontana* | Engle-Wrye A63 | MISSA | MISSA037105 | USA | TN | Sevier | 35.695778 | -83.389556 |
| **A63-7** | *Heuchera americana var. heteradenia* | HETERADENIA | fumosimontana | *Heuchera fumosimontana* | Engle-Wrye A63 | MISSA | MISSA037105 | USA | TN | Sevier | 35.695778 | -83.389556 |
| **A63-8** | *Heuchera americana var. heteradenia* | HETERADENIA | fumosimontana | *Heuchera fumosimontana* | Engle-Wrye A63 | MISSA | MISSA037105 | USA | TN | Sevier | 35.695778 | -83.389556 |
| **A63-9** | *Heuchera americana var. heteradenia* | HETERADENIA | fumosimontana | *Heuchera fumosimontana* | Engle-Wrye A63 | MISSA | MISSA037105 | USA | TN | Sevier | 35.695778 | -83.389556 |

Table S1: Continued

| | | | | | | | | | | | | |
|---|---|---|---|---|---|---|---|---|---|---|---|---|
| **A7-1** | *Heuchera americana var. americana* | AMERICANA | calycosa_yellow | *Heuchera americana var. alabamense* | Folk A7 | MISSA | To be processed | USA | SC | Abbeville | 34.0972222 | -82.351389 |
| **A7-2** | *Heuchera americana var. americana* | AMERICANA | calycosa_yellow | *Heuchera americana var. alabamense* | Folk A7 | MISSA | To be processed | USA | SC | Abbeville | 34.0972222 | -82.351389 |
| **A7-3** | *Heuchera americana var. americana* | AMERICANA | calycosa_yellow | *Heuchera americana var. alabamense* | Folk A7 | MISSA | To be processed | USA | SC | Abbeville | 34.0972222 | -82.351389 |
| **A7-4** | *Heuchera americana var. americana* | AMERICANA | calycosa_yellow | *Heuchera americana var. alabamense* | Folk A7 | MISSA | To be processed | USA | SC | Abbeville | 34.0972222 | -82.351389 |
| **A7-4** | *Heuchera americana var. americana* | AMERICANA | calycosa_yellow | *Heuchera americana var. alabamense* | Folk A7 | MISSA | To be processed | USA | SC | Abbeville | 34.0972222 | -82.351389 |
| **A8** | *Heuchera americana var. americana* | CALYCOSA | calycosa_yellow | *Heuchera americana var. alabamense* | Folk A8 | MISSA | MISSA036643 | USA | GA | Jasper | 33.2555556 | -83.6825 |
| **A9-1** | *Heuchera americana var. hirsuticaulis* | HIRSUTICAULIS | grayana_orange | *Heuchera grayana* | Folk A9 | MISSA | MISSA034515 | USA | IN | Warren | 40.3380556 | -87.316389 |
| **A9-3** | *Heuchera americana var. hirsuticaulis* | HIRSUTICAULIS | grayana_orange | *Heuchera grayana* | Folk A9 | MISSA | MISSA034515 | USA | IN | Warren | 40.3380556 | -87.316389 |

Table S1: Continued

| | | | | | | | | | | | | |
|---|---|---|---|---|---|---|---|---|---|---|---|---|
| **A9-4** | *Heuchera americana var. hirsuticaulis* | HIRSUTICAULIS | grayana_orange | *Heuchera grayana* | Folk A9 | MISSA | MISSA034515 | USA | IN | Warren | 40.3380556 | -87.316389 |
| **E1** | *Heuchera americana var. americana* | CALYCOSA | calycosa_yellow | *Heuchera americana var. alabamense* | Bowers 13612 | UNA | UNA00034487 | USA | AL | Shelby | 33.183991 | -86.999559 |
| **E12** | *Heuchera americana var. americana* | CALYCOSA | brevipetala_brown | *Heuchera americana var. brevipetala* | Williams s.n. | UNA | UNA00027468 | USA | AL | Jefferson | 33.5228184 | -86.916451 |
| **E1415** | *Heuchera americana var. americana* | AMERICANA | calycosa_yellow | *Heuchera americana var. alabamense* | Radford 220808 | NCU | NCU00180859 | USA | SC | Abbeville | 34.09611 | -82.344593 |
| **E1416** | *Heuchera americana var. americana* | AMERICANA | caroliniana | *Heuchera americana var. americana* | Ahles 56273 | NCU | NCU00180860 | USA | SC | Aiken | 33.542909 | -81.99475 |
| **E1419** | *Heuchera americana var. americana* | AMERICANA | caroliniana | *Heuchera americana var. americana* | Nelson 7394 | NCU | NCU00180863 | USA | SC | Cherokee | 33.1595951 | -79.907046 |
| **E1420** | *Heuchera americana var. americana* | AMERICANA | caroliniana | *Heuchera americana var. americana* | Haesloop 26792 | NCU | NCU00180864 | USA | SC | Cherokee | 35.13592 | -81.593086 |
| **E1421** | *Heuchera americana var. americana* | AMERICANA | caroliniana | *Heuchera americana var. americana* | Leonard 4250 | NCU | NCU00180865 | USA | SC | Dorchester | 35.0372626 | -81.648076 |

Table S1: Continued

| | | | | | | | | | | | | |
|---|---|---|---|---|---|---|---|---|---|---|---|---|
| **E1422** | *Heuchera americana var. americana* | AMERICANA | calycosa_yellow | *Heuchera americana var. alabamense* | Radford 22536 | NCU | NCU00180866 | USA | SC | Edgefield | 33.0756291 | -80.343113 |
| **E1423** | *Heuchera americana var. americana* | AMERICANA | caroliniana | *Heuchera americana var. americana* | Bell 7189 | NCU | NCU00180867 | USA | SC | Fairfield | 34.51148 | -81.297873 |
| **E1426** | *Heuchera americana var. americana* | AMERICANA | calycosa_yellow | *Heuchera americana var. alabamense* | Radford 22388 | NCU | NCU00180851 | USA | SC | McCormick | 34.4646451 | -81.994404 |
| **E1427** | *Heuchera americana var. americana* | AMERICANA | caroliniana | *Heuchera americana var. americana* | Bell 7024 | NCU | NCU00180852 | USA | SC | Newberry | 33.8795325 | -82.279772 |
| **E1429** | *Heuchera americana var. americana* | AMERICANA | calycosa_yellow | *Heuchera americana var. alabamense* | Knox 100 | NCU | NCU00180854 | USA | SC | Pickens | 34.8882592 | -82.719382 |
| **E1430** | *Heuchera americana var. americana* | AMERICANA | fumosimontana | *Heuchera americana var. americana* | Freeman 57187 | NCU | NCU00180855 | USA | SC | Pickens | 34.8882592 | -82.719382 |
| **E1432** | *Heuchera americana var. americana* | AMERICANA | caroliniana | *Heuchera americana var. americana* | Bell 8451 | NCU | NCU00180857 | USA | SC | Union | 34.9498007 | -81.932016 |
| **E1435** | *Heuchera caroliniana* | | caroliniana | *Heuchera caroliniana* | Ahles 27366 | NCU | NCU00053744 | USA | SC | Lancaster | 34.6859896 | -81.154507 |

Table S1: Continued

| | | | | | | | | | | | | |
|---|---|---|---|---|---|---|---|---|---|---|---|---|
| **E1436** | *Heuchera caroliniana* | | caroliniana | *Heuchera caroliniana* | Wells 3310 | NCU | NCU00053745 | USA | SC | Lancaster | 34.6628067 | -80.700546 |
| **E1438** | *Heuchera caroliniana* | | caroliniana | *Heuchera caroliniana* | Smith 67 | NCU | NCU00053743 | USA | SC | Darlington | 34.7622221 | -83.109965 |
| **E1457** | *Heuchera americana var. americana* | CALYCOSA | fumosimontana | *Heuchera americana var. americana* | Furr 582 | NCU | NCU00180880 | USA | KY | Bell | 33.762884 | -83.740416 |
| **E1459** | *Heuchera americana var. hirsuticaulis* | HIRSUTICAULIS | brevipetala_brown | *Heuchera hirsuticaulis* | Wilson 234 | NCU | NCU00180881 | USA | KY | Carlisle | 36.848 | -89.085 |
| **E1469** | *Heuchera americana var. americana* | BREVIPETALA | fumosimontana | *Heuchera americana var. americana* | Furr 586 | NCU | NCU00180898 | USA | TN | Claiborne | 35.6719722 | -83.931447 |
| **E1470** | *Heuchera americana var. americana* | BREVIPETALA | brevipetala_brown | *Heuchera americana var. brevipetala* | Sharp 23198 | NCU | NCU00180897 | USA | TN | Coffee | 36.4673161 | -83.678638 |
| **E1472** | *Heuchera americana var. americana* | BREVIPETALA | fumosimontana | *Heuchera americana var. americana* | Shaw s.n. | NCU | NCU00439719 | USA | TN | Fentress | 36.188076 | -85.002043 |
| **E1473** | *Heuchera americana var. americana* | BREVIPETALA | calycosa_yellow | *Heuchera americana var. brevipetala* | Lazell s.n. | NCU | NCU00180894 | USA | TN | Franklin | 36.360098 | -84.926231 |

Table S1: Continued

| | | | | | | | | | | | | |
|---|---|---|---|---|---|---|---|---|---|---|---|---|
| **E1474** | *Heuchera americana var. americana* | BREVIPETALA | fumosimontana | *Heuchera americana var. americana* | Furr 337 | NCU | NCU00180900 | USA | TN | Grainger | 35.925206 | -86.868942 |
| **E1475** | *Heuchera americana var. americana* | BREVIPETALA | fumosimontana | *Heuchera americana var. americana* | Clark 1760 | NCU | NCU00181081 | USA | TN | Grundy | 36.2810821 | -83.510702 |
| **E1476** | *Heuchera americana var. americana* | BREVIPETALA | fumosimontana | *Heuchera americana var. americana* | Furr 366 | NCU | NCU00181082 | USA | TN | Hawkins | 35.3697894 | -85.710661 |
| **E1477** | *Heuchera americana var. americana* | AMERICANA | fumosimontana | *Heuchera americana var. americana* | Wells 3286 | NCU | NCU00181083 | USA | TN | Jefferson | 36.429878 | -82.95689 |
| **E1482** | *Heuchera americana var. americana* | BREVIPETALA | fumosimontana | *Heuchera americana var. americana* | Hess 1150 | NCU | NCU00181090 | USA | TN | Monroe | 35.4397238 | -84.239964 |
| **E1486** | *Heuchera americana var. americana* | BREVIPETALA | americana_green | *Heuchera americana var. americana* | Furr 292 | NCU | NCU00181093 | USA | TN | Swain | 35.84857 | -84.522552 |
| **E1490** | *Heuchera americana var. americana* | AMERICANA | alba_lavender | *Heuchera americana var. americana* | Wells 3292 | NCU | NCU00181097 | USA | VA | Botetourt | 37.5520825 | -79.802957 |
| **E1491** | *Heuchera americana var. americana* | AMERICANA | caroliniana | *Heuchera americana var. americana* | Kirkland 23220 | NCU | NCU00181098 | USA | VA | Brunswick | 37.5520825 | -79.802957 |

Table S1: Continued

| | | | | | | | | | | | | |
|---|---|---|---|---|---|---|---|---|---|---|---|---|
| **E1492** | *Heuchera americana var. americana* | AMERICANA | alba_lavender | *Heuchera americana var. americana* | Ramsey 7535 | NCU | NCU00181099 | USA | VA | Buckingham | 36.7788996 | -77.867088 |
| **E1493** | *Heuchera americana var. americana* | AMERICANA | alba_lavender | *Heuchera americana var. americana* | Howell 16501 | NCU | NCU00181100 | USA | VA | Buckingham | 37.5558826 | -78.554717 |
| **E1494** | *Heuchera americana var. americana* | AMERICANA | alba_lavender | *Heuchera americana var. americana* | Ware 4210 | NCU | NCU00181101 | USA | VA | Charles City | 37.5558826 | -78.554717 |
| **E1495** | *Heuchera americana var. americana* | AMERICANA | alba_lavender | *Heuchera americana var. americana* | Bradley 6826 | NCU | NCU00181104 | USA | VA | Fairfax | 37.3705777 | -77.06051 |
| **E1496** | *Heuchera americana var. americana* | AMERICANA | alba_lavender | *Heuchera americana var. americana* | Diggs 226 | NCU | NCU00181105 | USA | VA | Fluvanna | 38.8462236 | -77.306373 |
| **E1497** | *Heuchera pubescens* | | americana_green | *Heuchera pubescens* | Weiboldt 8150 | NCU | NCU00181106 | USA | VA | Franklin | 36.838161 | -79.727771 |
| **E1498** | *Heuchera americana var. americana* | AMERICANA | alba_lavender | *Heuchera americana var. americana* | Wells 3363 | NCU | NCU00181107 | USA | VA | Giles | 37.0527719 | -79.881309 |
| **E1499** | *Heuchera americana var. americana* | AMERICANA | alba_lavender | *Heuchera americana var. americana* | Kimsey 122 | NCU | NCU00181108 | USA | VA | Goochland | 37.3088528 | -80.709878 |

Table S1: Continued

| | | | | | | | | | | | | |
|---|---|---|---|---|---|---|---|---|---|---|---|---|
| **E1500** | *Heuchera americana var. americana* | AMERICANA | caroliniana | *Heuchera americana var. americana* | Massey 4014 | NCU | NCU00181111 | USA | VA | Halifax | 37.7204342 | -77.883798 |
| **E1501** | *Heuchera americana var. americana* | AMERICANA | alba_lavender | *Heuchera americana var. americana* | Ramsey 4173 | NCU | NCU00181109 | USA | VA | Halifax | 36.747244 | -78.942563 |
| **E1502** | *Heuchera americana var. americana* | AMERICANA | caroliniana | *Heuchera americana var. americana* | Boufford 13886 | NCU | NCU00181110 | USA | VA | Halifax | 36.747244 | -78.942563 |
| **E1503** | *Heuchera americana var. americana* | AMERICANA | alba_lavender | *Heuchera americana var. americana* | Wells 3291 | NCU | NCU00181113 | USA | VA | Hanover | 36.747244 | -78.942563 |
| **E1504** | *Heuchera americana var. americana* | AMERICANA | caroliniana | *Heuchera americana var. americana* | Dougherty s.n. | NCU | NCU00181115 | USA | VA | Richmond City | 37.744783 | -77.446417 |
| **E1505** | *Heuchera americana var. americana* | AMERICANA | alba_lavender | *Heuchera americana var. americana* | Wells 3294 | NCU | NCU00181116 | USA | VA | Isle of Wright | 37.034167 | -76.619444 |
| **E1506** | *Heuchera americana var. americana* | AMERICANA | caroliniana | *Heuchera americana var. americana* | Borans 201 | NCU | NCU00181118 | USA | VA | James City | 37.271705 | -76.714104 |
| **E1507** | *Heuchera americana var. americana* | AMERICANA | alba_lavender | *Heuchera americana var. americana* | Gillespie 635 | NCU | NCU00181122 | USA | VA | New Kent | 37.581311 | -77.062158 |

Table S1: Continued

| | | | | | | | | | | | | |
|---|---|---|---|---|---|---|---|---|---|---|---|---|
| **E1508** | *Heuchera americana var. americana* | AMERICANA | alba_lavender | *Heuchera americana var. americana* | Mikula 5348 | NCU | NCU00181123 | USA | VA | Page | 37.4974701 | -76.999045 |
| **E1510** | *Heuchera americana var. americana* | AMERICANA | alba_lavender | *Heuchera americana var. americana* | Corcoran 296 | NCU | NCU00181126 | USA | VA | Powhatan | 36.6818706 | -80.284636 |
| **E1512** | *Heuchera americana var. americana* | AMERICANA | alba_lavender | *Heuchera americana var. americana* | Wells 3295 | NCU | NCU00181128 | USA | VA | Surry | 37.1118778 | -76.895924 |
| **E1513** | *Heuchera americana var. americana* | AMERICANA | americana_green | *Heuchera americana var. americana* | Fur 439 | NCU | NCU00181129 | USA | VA | Washington | 37.1118778 | -76.895924 |
| **E1514** | *Heuchera americana var. americana* | AMERICANA | alba_lavender | *Heuchera americana var. americana* | Salle 324 | NCU | NCU00181130 | USA | VA | York | 36.740236 | -81.942167 |
| **E1515** | *Heuchera americana var. americana* | CALYCOSA | brevipetala_brown | *Heuchera americana var. brevipetala* | Massey 4627 | NCU | NCU00181131 | USA | AL | Jackson | 34.744036 | -86.30427 |
| **E1516** | *Heuchera americana var. americana* | CALYCOSA | brevipetala_brown | *Heuchera americana var. brevipetala* | Lelong 4564 | NCU | NCU00181132 | USA | AL | Marion | 34.7692447 | -85.986794 |
| **E1517** | *Heuchera americana var. americana* | BREVIPETALA | hirsuticaulis_blue | *Heuchera americana var. brevipetala* | Tucker 4003 | NCU | NCU00128865 | USA | AR | Johnson | 35.624429 | -93.291523 |

Table S1: Continued

| | | | | | | | | | | | | |
|---|---|---|---|---|---|---|---|---|---|---|---|---|
| **E1518** | *Heuchera americana var. americana* | BREVIPETALA | hirsuticaulis_red | *Heuchera americana var. brevipetala* | Merrill 1844 | NCU | NCU00128866 | USA | AR | Pulaski | 35.5600697 | -93.455011 |
| **E1519** | *Heuchera americana var. americana* | CALYCOSA | caroliniana | *Heuchera americana var. americana* | Younors s.n. | NCU | NCU00181133 | USA | GA | Bulloch | 32.393408 | -81.74381 |
| **E1520** | *Heuchera americana var. americana* | BREVIPETALA | hirsuticaulis_red | *Heuchera americana var. brevipetala* | Thieret 22647 | NCU | NCU00181134 | USA | LA | Caddo | 32.577195 | -93.882423 |
| **E1521** | *Heuchera americana var. americana* | AMERICANA | brevipetala_pink | *Heuchera americana var. brevipetala* | Windler 3822 | NCU | NCU00181135 | USA | MD | Baltimore | 39.417425 | -76.541639 |
| **E1522** | *Heuchera americana var. americana* | AMERICANA | alba_lavender | *Heuchera americana var. americana* | Hickey 348 | NCU | NCU00181136 | USA | MD | Frederick | 39.2908816 | -76.610759 |
| **E1523** | *Heuchera americana var. americana* | HIRSUTICAULIS | hirsuticaulis_red | *Heuchera americana var. brevipetala* | Richards 3790 | NCU | NCU00142730 | USA | AR | Craighead | 36.142026 | -90.575279 |
| **E1524** | *Heuchera americana var. hirsuticaulis* | HIRSUTICAULIS | hirsuticaulis_red | *Heuchera hirsuticaulis* | Demaree 33248 | NCU | NCU00092970 | USA | AR | Craighead | 35.8348413 | -90.644114 |
| **E1525** | *Heuchera americana var. hirsuticaulis* | HIRSUTICAULIS | hirsuticaulis_blue | *Heuchera hirsuticaulis* | Demaree 5138 | NCU | NCU00092973 | USA | AR | Franklin | 35.8348413 | -90.644114 |

Table S1: Continued

| | | | | | | | | | | | | |
|---|---|---|---|---|---|---|---|---|---|---|---|---|
| **E1526** | *Heuchera americana var. hirsuticaulis* | HIRSUTICAULIS | hirsuticaulis_blue | *Heuchera hirsuticaulis* | Stephens 10575 | NCU | NCU00092972 | USA | AR | Franklin | 35.5110075 | -93.886485 |
| **E1527** | *Heuchera americana var. hirsuticaulis* | HIRSUTICAULIS | hirsuticaulis_red | *Heuchera hirsuticaulis* | Hardin 634 | NCU | NCU00092971 | USA | AR | Franklin | 35.5110075 | -93.886485 |
| **E1529** | *Heuchera americana var. hirsuticaulis* | HIRSUTICAULIS | hirsuticaulis_red | *Heuchera hirsuticaulis* | Iltis 5477 | NCU | NCU00092974 | USA | AR | Newton | 35.943319 | -93.070165 |
| **E1531** | *Heuchera americana var. hirsuticaulis* | HIRSUTICAULIS | hirsuticaulis_red | *Heuchera hirsuticaulis* | Hawkins 385 | NCU | NCU00142731 | USA | AR | Randolph | 36.306896 | -91.085753 |
| **E1532** | *Heuchera americana var. hirsuticaulis* | HIRSUTICAULIS | hirsuticaulis_red | *Heuchera hirsuticaulis* | Furr 649 | NCU | NCU00092977 | USA | AR | Searcy | 36.3412315 | -91.038702 |
| **E1533** | *Heuchera americana var. hirsuticaulis* | HIRSUTICAULIS | hirsuticaulis_red | *Heuchera hirsuticaulis* | Redfearn 31682 | NCU | NCU00128867 | USA | AR | Stone | 35.995665 | -92.297486 |
| **E1534** | *Heuchera americana var. hirsuticaulis* | HIRSUTICAULIS | hirsuticaulis_red | *Heuchera hirsuticaulis* | Demaree 49855 | NCU | NCU00092978 | USA | AR | Van Buren | 35.8317159 | -92.174763 |
| **E1535** | *Heuchera americana var. hirsuticaulis* | HIRSUTICAULIS | hirsuticaulis_blue | *Heuchera hirsuticaulis* | Hughes 76 | NCU | NCU00092981 | USA | AR | Washington | 35.5759086 | -92.491614 |

Table S1: Continued

| | | | | | | | | | | | | |
|---|---|---|---|---|---|---|---|---|---|---|---|---|
| **E1536** | *Heuchera americana var. hirsuticaulis* | HIRSUTICAULIS | brevipetala_brown | *Heuchera hirsuticaulis* | Wells 3301 | NCU | NCU00092986 | USA | KY | Edmonson | 35.9672625 | -94.228066 |
| **E1537** | *Heuchera americana var. hirsuticaulis* | HIRSUTICAULIS | brevipetala_brown | *Heuchera hirsuticaulis* | Windler 2497 | NCU | NCU00092991 | USA | KY | Livingston | 37.1999181 | -86.220674 |
| **E1538** | *Heuchera americana var. hirsuticaulis* | HIRSUTICAULIS | brevipetala_brown | *Heuchera hirsuticaulis* | Ellis 01089 | NCU | NCU00092982 | USA | KY | Lyon | 37.1898627 | -88.334383 |
| **E1539** | *Heuchera americana var. hirsuticaulis* | HIRSUTICAULIS | brevipetala_brown | *Heuchera hirsuticaulis* | Ellis 01338 | NCU | NCU00092987 | USA | KY | Trigg | 36.9981232 | -88.065815 |
| **E1540** | *Heuchera americana var. hirsuticaulis* | HIRSUTICAULIS | brevipetala_brown | *Heuchera hirsuticaulis* | Ellis 01393 | NCU | NCU00092983 | USA | KY | Trigg | 36.7949221 | -87.878243 |
| **E1541** | *Heuchera americana var. heteradenia* | HETERADENIA | calycosa_yellow | *Heuchera americana var. alabamense* | Parrish 40 | NCU | NCU00092996 | USA | GA | Screven | 32.744751 | -81.617585 |
| **E1542** | *Heuchera americana var. americana* | CALYCOSA | calycosa_yellow | *Heuchera americana var. alabamense* | Park s.n. | NCU | NCU00092992 | USA | GA | Screven | 32.744751 | -81.617585 |
| **E1543** | *Heuchera americana var. hirsuticaulis* | HIRSUTICAULIS | brevipetala_brown | *Heuchera hirsuticaulis* | Clebsch s.n. | NCU | NCU00092988 | USA | TN | Montgomery | 36.439722 | -87.296667 |

Table S1: Continued

| | | | | | | | | | | | | |
|---|---|---|---|---|---|---|---|---|---|---|---|---|
| **E1544** | *Heuchera americana var. americana* | BREVIPETALA | fumosimontana | *Heuchera americana var. americana* | Shaw s.n. | NCU | NCU00439730 | USA | TN | Polk | 35.103928 | -84.55709 |
| **E1546** | *Heuchera americana var. hispida* | HISPIDA | alba_lavender | *Heuchera americana var. hispida* | Sharp 319 | NCU | NCU00045132 | USA | VA | Augusta | | |
| **E1547** | *Heuchera americana var. hispida* | HISPIDA | alba_lavender | *Heuchera americana var. hispida* | Freer 3943 | NCU | NCU00045133 | USA | VA | Augusta | 38.1793809 | -79.152576 |
| **E1548** | *Heuchera americana var. hispida* | HISPIDA | alba_lavender | *Heuchera americana var. hispida* | Freer 1564 | NCU | NCU00045134 | USA | VA | Bedford | 38.1793809 | -79.152576 |
| **E1549** | *Heuchera americana var. hispida* | HISPIDA | alba_lavender | *Heuchera americana var. hispida* | Furr 497 | NCU | NCU00045135 | USA | VA | Botetourt | 37.545183 | -79.971503 |
| **E1550** | *Heuchera americana var. hispida* | HISPIDA | alba_lavender | *Heuchera americana var. hispida* | Furr 484 | NCU | NCU00045136 | USA | VA | Botetourt | 37.544884 | -79.971284 |
| **E1551** | *Heuchera americana var. hispida* | HISPIDA | alba_lavender | *Heuchera americana var. hispida* | Fernald 1596 | NCU | NCU00045137 | USA | VA | Botetourt | 37.482975 | -79.668629 |
| **E1552** | *Heuchera americana var. hispida* | HISPIDA | alba_lavender | *Heuchera americana var. hispida* | Wells 3313 | NCU | NCU00045138 | USA | VA | Craig | 37.534587 | -80.235484 |

Table S1: Continued

| | | | | | | | | | | | | |
|---|---|---|---|---|---|---|---|---|---|---|---|---|
| **E1553** | *Heuchera americana var. hispida* | HISPIDA | alba_lavender | *Heuchera americana var. hispida* | Furr 480 | NCU | NCU00045139 | USA | VA | Craig | 37.542294 | -79.969289 |
| **E1554** | *Heuchera americana var. hispida* | HISPIDA | alba_lavender | *Heuchera americana var. hispida* | Furr 474 | NCU | NCU00045140 | USA | VA | Craig | 37.48337 | -80.130283 |
| **E1556** | *Heuchera americana var. hispida* | HISPIDA | alba_lavender | *Heuchera americana var. hispida* | Hawill 14127 | NCU | NCU00045142 | USA | VA | Craig | 37.357343 | -80.439593 |
| **E1557** | *Heuchera americana var. hispida* | HISPIDA | alba_lavender | *Heuchera americana var. hispida* | Wells 3293 | NCU | NCU00045143 | USA | VA | Fauquier | 38.825515 | -77.719505 |
| **E1558** | *Heuchera americana var. hispida* | HISPIDA | alba_lavender | *Heuchera americana var. hispida* | Massey W3315 | NCU | NCU00045144 | USA | VA | Giles | 37.605918 | -80.240697 |
| **E1559** | *Heuchera americana var. hispida* | HISPIDA | americana_green | *Heuchera americana var. hispida* | Cooperrider 4456 | NCU | NCU00045145 | USA | VA | Giles | 37.270794 | -80.694765 |
| **E1560** | *Heuchera americana var. hispida* | HISPIDA | americana_green | *Heuchera americana var. hispida* | Furr 464 | NCU | NCU00045146 | USA | VA | Giles | 37.266857 | -80.656989 |
| **E1561** | *Heuchera americana var. hispida* | HISPIDA | americana_green | *Heuchera americana var. hispida* | Harvill 14272 | NCU | NCU00045147 | USA | VA | Giles | 37.605918 | -80.240697 |

Table S1: Continued

| | | | | | | | | | | | | |
|---|---|---|---|---|---|---|---|---|---|---|---|---|
| **E1562** | *Heuchera americana var. hispida* | HISPIDA | americana_green | *Heuchera americana var. hispida* | Harvill 16432 | NCU | NCU00045148 | USA | VA | Grayson | 36.576455 | -81.158355 |
| **E1563** | *Heuchera americana var. hispida* | HISPIDA | alba_lavender | *Heuchera americana var. hispida* | Harvill 19416 | NCU | NCU00045149 | USA | VA | Loudoun | 39.319112 | -77.710924 |
| **E1564** | *Heuchera americana var. hispida* | HISPIDA | americana_green | *Heuchera americana var. hispida* | Kral 10351 | NCU | NCU00045150 | USA | VA | Montgomery | 37.244208 | -80.604519 |
| **E1565** | *Heuchera americana var. hispida* | HISPIDA | alba_lavender | *Heuchera americana var. hispida* | Smyth 1055 | NCU | NCU00045151 | USA | VA | Montgomery | 38.1803833 | -81.328445 |
| **E1566** | *Heuchera americana var. hispida* | HISPIDA | alba_lavender | *Heuchera americana var. hispida* | Ahles 62433 | NCU | NCU00045153 | USA | VA | Pittsylvania | 37.067349 | -79.105487 |
| **E1567** | *Heuchera americana var. hispida* | HISPIDA | americana_green | *Heuchera americana var. hispida* | Furr 454 | NCU | NCU00045154 | USA | VA | Pulaski | 37.208469 | -80.738294 |
| **E1568** | *Heuchera americana var. hispida* | HISPIDA | alba_lavender | *Heuchera americana var. hispida* | Wood 5819 | NCU | NCU00045155 | USA | VA | Roanoke | 37.140928 | -80.118981 |
| **E1569** | *Heuchera americana var. hispida* | HISPIDA | fumosimontana | *Heuchera americana var. hispida* | Furr 393 | NCU | NCU00045157 | USA | VA | Scott | 37.270973 | -79.941431 |

Table S1: Continued

| | | | | | | | | | | | | |
|---|---|---|---|---|---|---|---|---|---|---|---|---|
| **E1570** | *Heuchera americana var. hispida* | HISPIDA | americana_green | *Heuchera americana var. hispida* | Uttal 10455 | NCU | NCU00045158 | USA | VA | Tazewell | 36.7117473 | -82.589305 |
| **E1571** | *Heuchera americana var. hispida* | HISPIDA | fumosimontana | *Heuchera americana var. hispida* | Furr 434 | NCU | NCU00045159 | USA | VA | Wise | 36.90057 | -82.310079 |
| **E1572** | *Heuchera americana var. hispida* | HISPIDA | americana_green | *Heuchera americana var. hispida* | Furr 449 | NCU | NCU00045160 | USA | VA | Wythe | 36.886036 | -81.192015 |
| **E1573** | *Heuchera americana var. hispida* | HISPIDA | alba_lavender | *Heuchera americana var. hispida* | Downs 3488 | NCU | NCU00196809 | USA | MD | Washington | 39.346451 | -77.726568 |
| **E1574** | *Heuchera americana var. hispida* | HISPIDA | alba_lavender | *Heuchera americana var. hispida* | Wells 3314 | NCU | NCU00053746 | USA | WV | Greenbrier | 37.806692 | 80.06763 |
| **E1575** | *Heuchera americana var. hispida* | HISPIDA | alba_lavender | *Heuchera americana var. hispida* | Wells 3312 | NCU | NCU00053747 | USA | WV | Monroe | 37.961175 | -80.450934 |
| **E1576** | *Heuchera americana var. hispida* | HISPIDA | alba_lavender | *Heuchera americana var. hispida* | Southern Appalachian Botanical Club 320 | NCU | NCU00053748 | USA | WV | Pocahontas | 37.5585821 | -80.519223 |

Table S1: Continued

| | | | | | | | | | | | | |
|---|---|---|---|---|---|---|---|---|---|---|---|---|
| **E1578** | *Heuchera americana var. americana* | CALYCOSA | calycosa_yellow | *Heuchera americana var. alabamense* | Bussey 614 | NCU | NCU00181137 | USA | AL | Clay | 37.5969073 | -81.536493 |
| **E1579** | *Heuchera americana var. americana* | CALYCOSA | calycosa_yellow | *Heuchera americana var. alabamense* | Orzell 9505 | NCU | NCU00181138 | USA | AL | Cleburne | 33.721111 | -85.604444 |
| **E1580** | *Heuchera americana var. hirsuticaulis* | HIRSUTICAULIS | hirsuticaulis_red | *Heuchera hirsuticaulis* | Lipscomb 1564 | NCU | NCU00128864 | USA | AR | Izard | 36.118644 | -92.153699 |
| **E1581** | *Heuchera americana var. americana* | CALYCOSA | calycosa_yellow | *Heuchera americana var. alabamense* | Hill 349 | NCU | NCU00181139 | USA | GA | Morgan | 33.661916 | -83.593355 |
| **E1596** | *Heuchera pubescens* | | alba_lavender | *Heuchera pubescens* | Gupton 3549 | NCU | NCU00190348 | USA | VA | Bath | 38.038103 | 79.763563 |
| **E1598** | *Heuchera pubescens* | | alba_lavender | *Heuchera pubescens* | Wells 3316 | NCU | NCU00190350 | USA | VA | Craig | 37.546354 | -79.976945 |
| **E1599** | *Heuchera pubescens* | | alba_lavender | *Heuchera pubescens* | Ramsey 4693 | NCU | NCU00190352 | USA | VA | Franklin | 36.999573 | -79.878086 |
| **E1600** | *Heuchera pubescens* | | alba_lavender | *Heuchera pubescens* | Mitchell 4107 | NCU | NCU00190353 | USA | VA | Franklin | 37.007717 | -79.889332 |
| **E1601** | *Heuchera pubescens* | | alba_lavender | *Heuchera pubescens* | Kral 10306 | NCU | NCU00190354 | USA | VA | Montgomery | 37.215641 | -80.268486 |
| **E1602** | *Heuchera pubescens* | | alba_lavender | *Heuchera pubescens* | Johnson 4565 | NCU | NCU00190355 | USA | VA | Patrick | 36.720388 | -80.327535 |

Table S1: Continued

| | | | | | | | | | | | | |
|---|---|---|---|---|---|---|---|---|---|---|---|---|
| **E1603** | *Heuchera pubescens* | | alba_lavender | *Heuchera pubescens* | Ahles 60159 | NCU | NCU00190356 | USA | VA | Patrick | 36.717451 | -80.323062 |
| **E1677** | *Heuchera americana var. hispida* | HISPIDA | alba_lavender | *Heuchera americana var. hispida* | Kral 63868 | NCU | NCU00386386 | USA | VA | Montgomery | 37.191354 | -80.36668 |
| **E1679** | *Heuchera americana var. americana* | BREVIPETALA | brevipetala_pink | *Heuchera americana var. brevipetala* | Cusick 28987 | NCU | NCU00190481 | USA | KY | Greenup | 38.658386 | -83.04535 |
| **E1680** | *Heuchera americana var. hirsuticaulis* | HIRSUTICAULIS | hirsuticaulis_red | *Heuchera hirsuticaulis* | Ugent 81-70 | NCU | NCU00196305 | USA | IL | Calhoun | 38.999907 | -90.588731 |
| **E1681** | *Heuchera americana var. hirsuticaulis* | HIRSUTICAULIS | hirsuticaulis_red | *Heuchera hirsuticaulis* | Raven 27489 | NCU | NCU00196306 | USA | MO | Jefferson | 38.125 | -90.675 |
| **E1682** | *Heuchera americana var. americana* | BREVIPETALA | brevipetala_pink | *Heuchera americana var. brevipetala* | Wells 3298 | NCU | NCU00196308 | USA | NJ | Hunterdon | 40.475307 | -75.05958 |
| **E1683** | *Heuchera americana var. americana* | BREVIPETALA | brevipetala_pink | *Heuchera americana var. brevipetala* | Cusick 596 | NCU | NCU00196316 | USA | OH | Jefferson | 40.353147 | -80.691019 |
| **E1684** | *Heuchera americana var. americana* | BREVIPETALA | brevipetala_pink | *Heuchera americana var. brevipetala* | Cusick 7568 | NCU | NCU00196317 | USA | OH | Monroe | 39.65929 | -81.06818 |

Table S1: Continued

| | | | | | | | | | | | | |
|---|---|---|---|---|---|---|---|---|---|---|---|---|
| **E1685** | *Heuchera americana var. americana* | BREVIPETALA | brevipetala_pink | *Heuchera americana var. brevipetala* | Cusick 7961 | NCU | NCU00196318 | USA | OH | Noble | 39.7074 | -81.583942 |
| **E1686** | *Heuchera americana var. americana* | BREVIPETALA | brevipetala_pink | *Heuchera americana var. brevipetala* | Cooperrider 6745 | NCU | NCU00196319 | USA | OH | Portage | 41.216555 | -81.301325 |
| **E1687** | *Heuchera americana var. americana* | BREVIPETALA | hirsuticaulis_red | *Heuchera americana var. brevipetala* | Ziegler 371 | NCU | NCU00196320 | USA | OK | McCurtain | 34.09695 | -94.7043 |
| **E1688** | *Heuchera americana var. americana* | BREVIPETALA | brevipetala_pink | *Heuchera americana var. brevipetala* | Wells 3299 | NCU | NCU00196310 | USA | PA | Bucks | 40.563889 | -75.097729 |
| **E1689** | *Heuchera americana var. americana* | BREVIPETALA | brevipetala_pink | *Heuchera americana var. brevipetala* | Krouse s.n. | NCU | NCU00196312 | USA | PA | Fayette | 39.871742 | -79.492261 |
| **E1690** | *Heuchera americana var. americana* | BREVIPETALA | brevipetala_pink | *Heuchera americana var. brevipetala* | Wells 3297 | NCU | NCU00196313 | USA | PA | Lancaster | 39.940464 | -75.993633 |
| **E1691** | *Heuchera americana var. americana* | BREVIPETALA | brevipetala_pink | *Heuchera americana var. brevipetala* | Utech 82-188 | NCU | NCU00196315 | USA | PA | Westmoreland | 40.129167 | -79.291667 |
| **E1692** | *Heuchera americana var. americana* | BREVIPETALA | hirsuticaulis_red | *Heuchera americana var. brevipetala* | Correll 37142 | NCU | NCU00196493 | USA | TX | Bowie | 33.516333 | -94.147 |

Table S1: Continued

| | | | | | | | | | | | | |
|---|---|---|---|---|---|---|---|---|---|---|---|---|
| **E1693** | *Heuchera americana var. hirsuticaulis* | HIRSUTICAULIS | hirsuticaulis_red | *Heuchera hirsuticaulis* | Furr 592 | NCU | NCU00196495 | USA | IL | Alexander | 37.208057 | -89.428705 |
| **E1694** | *Heuchera americana var. hirsuticaulis* | HIRSUTICAULIS | brevipetala_pink | *Heuchera hirsuticaulis* | Winterringer 901 | NCU | NCU00196496 | USA | IL | Hardin | 37.4999684 | -88.237835 |
| **E1697** | *Heuchera americana var. hirsuticaulis* | HIRSUTICAULIS | brevipetala_brown | *Heuchera hirsuticaulis* | Bollwinkel FC 124 | NCU | NCU00196500 | USA | IL | Johnson | 37.4501818 | -88.884405 |
| **E1698** | *Heuchera americana var. hirsuticaulis* | HIRSUTICAULIS | hirsuticaulis_red | *Heuchera hirsuticaulis* | Neill 15228 | NCU | NCU00196550 | USA | IL | Saint Clair | 38.4616972 | -89.932435 |
| **E1699** | *Heuchera americana var. hirsuticaulis* | HIRSUTICAULIS | brevipetala_pink | *Heuchera hirsuticaulis* | Bartlett 2109 | NCU | NCU00196551 | USA | IN | Brown | 39.1682855 | -86.2297 |
| **E1702** | *Heuchera americana var. hirsuticaulis* | HIRSUTICAULIS | brevipetala_pink | *Heuchera hirsuticaulis* | Wells 3306 | NCU | NCU00196555 | USA | IN | Howard | 40.4787061 | -86.135034 |
| **E1703** | *Heuchera americana var. hirsuticaulis* | HIRSUTICAULIS | brevipetala_pink | *Heuchera hirsuticaulis* | Wells 3303 | NCU | NCU00196556 | USA | IN | Montgomery | 40.0361447 | -86.900708 |
| **E1704** | *Heuchera americana var. hirsuticaulis* | GRAYANA | richardsonii | *Heuchera richardsonii* | Furr 594 | NCU | NCU00196559 | USA | MO | Boone | 39.0153926 | -92.330753 |

Table S1: Continued

| | | | | | | | | | | | | |
|---|---|---|---|---|---|---|---|---|---|---|---|---|
| **E1705** | *Heuchera americana var. hirsuticaulis* | HIRSUTICAULIS | hirsuticaulis_red | *Heuchera hirsuticaulis* | Furr 668 | NCU | NCU00196801 | USA | MO | Shannon | 37.1498052 | -91.432825 |
| **E1706** | *Heuchera americana var. hirsuticaulis* | HIRSUTICAULIS | hirsuticaulis_red | *Heuchera hirsuticaulis* | Redfearn 27414 | NCU | NCU00196802 | USA | MO | Shannon | 37.093791 | -91.209527 |
| **E1707** | *Heuchera americana var. hirsuticaulis* | HIRSUTICAULIS | hirsuticaulis_red | *Heuchera hirsuticaulis* | Dorr 387 | NCU | NCU00196803 | USA | MO | Saint Louis | 38.6319657 | -90.242876 |
| **E1708** | *Heuchera americana var. hirsuticaulis* | HIRSUTICAULIS | hirsuticaulis_red | *Heuchera hirsuticaulis* | Furr 665 | NCU | NCU00196804 | USA | MO | Taney | 36.6563729 | -93.066578 |
| **E1709** | *Heuchera americana var. americana* | BREVIPETALA | brevipetala_pink | *Heuchera americana var. brevipetala* | O'Dell 276 | NCU | NCU00196806 | USA | OH | Vinton | 39.2744355 | -82.47403 |
| **E1711** | *Heuchera americana var. hirsuticaulis* | HIRSUTICAULIS | hirsuticaulis_blue | *Heuchera hirsuticaulis* | Wallis 6790-1 | NCU | NCU00196808 | USA | OK | Sequoyah | 35.5035863 | -94.736058 |
| **E1780** | *Heuchera richardsonii* | GRAYANA | grayana_orange | *Heuchera grayana* | Huang 3045 | NCU | NCU00196884 | USA | IA | Cedar | 41.7608608 | -91.12632 |
| **E1781** | *Heuchera richardsonii* | GRAYANA | grayana_orange | *Heuchera grayana* | Cooperrider 1120 | NCU | NCU00196885 | USA | IA | Jackson | 42.1420382 | -90.548007 |
| **E1782** | *Heuchera richardsonii* | GRAYANA | grayana_orange | *Heuchera grayana* | Walker 231 | NCU | NCU00196894 | USA | IA | Jackson | 42.1420382 | -90.548007 |

Table S1: Continued

| | | | | | | | | | | | | |
|---|---|---|---|---|---|---|---|---|---|---|---|---|
| **E1783** | *Heuchera richardsonii* | GRAYANA | richardsonii | *Heuchera richardsonii* | Witlake 905 | NCU | NCU00196887 | USA | IA | Lyon | 43.3747071 | -96.208192 |
| **E1784** | *Heuchera richardsonii* | GRAYANA | richardsonii | *Heuchera richardsonii* | DeBurh 1184 | NCU | NCU00196888 | USA | IA | Lyon | 43.3747071 | -96.208192 |
| **E1785** | *Heuchera richardsonii* | GRAYANA | richardsonii | *Heuchera richardsonii* | Myron 13 | NCU | NCU00196889 | USA | IA | Pocahontas | 42.7262681 | -94.647738 |
| **E1786** | *Heuchera richardsonii* | GRAYANA | richardsonii | *Heuchera richardsonii* | Schwab 209 | NCU | NCU00196891 | USA | IA | Story | 42.040106 | -93.634508 |
| **E1787** | *Heuchera richardsonii* | GRAYANA | richardsonii | *Heuchera richardsonii* | Freckmann 1882 | NCU | NCU00196892 | USA | IA | Story | 42.0161612 | -93.489194 |
| **E1788** | *Heuchera richardsonii* | GRAYANA | grayana_orange | *Heuchera grayana* | Wagenknecht 287 | NCU | NCU00196893 | USA | IA | Washington | 41.3160082 | -91.734033 |
| **E1789** | *Heuchera richardsonii* | GRAYANA | grayana_orange | *Heuchera grayana* | Utech 1925 | NCU | NCU00196912 | USA | WI | Crawford | 43.19824 | -90.874083 |
| **E1790** | *Heuchera richardsonii* | GRAYANA | grayana_orange | *Heuchera grayana* | Rice 1902 | NCU | NCU00196913 | USA | WI | Rock | 42.606119 | -89.02125 |
| **E1791** | *Heuchera richardsonii* | GRAYANA | grayana_orange | *Heuchera grayana* | Rice W3351 | NCU | NCU00196915 | USA | WI | Rock | 42.606119 | -89.02125 |
| **E1792** | *Heuchera richardsonii* | GRAYANA | grayana_orange | *Heuchera grayana* | Rice W3349 | NCU | NCU00196917 | USA | WI | Rock | 42.824114 | -89.318987 |
| **E1793** | *Heuchera richardsonii* | GRAYANA | grayana_orange | *Heuchera grayana* | Rice W3350 | NCU | NCU00196918 | USA | WI | Rock | 42.59156 | -89.021029 |
| **E1794** | *Heuchera richardsonii* | RICHARDSONII | richardsonii | *Heuchera richardsonii* | Jackson 628 | NCU | NCU00196898 | USA | CO | El Paso | 38.827383 | -104.52747 |

Table S1: Continued

| | | | | | | | | | | | | |
|---|---|---|---|---|---|---|---|---|---|---|---|---|
| **E1795** | *Heuchera richardsonii* | RICHARDSONII | grayana_orange | *Heuchera grayana* | Wells 3347 | NCU | NCU00196899 | USA | IN | Fulton | 41.0421416 | -86.287521 |
| **E1796** | *Heuchera richardsonii* | GRAYANA | grayana_orange | *Heuchera grayana* | Wells 3346 | NCU | NCU00196900 | USA | IN | Porter | 41.4457032 | -87.072497 |
| **E1797** | *Heuchera richardsonii* | GRAYANA | richardsonii | *Heuchera richardsonii* | McGregor 15586 | NCU | NCU00196901 | USA | KS | Cherokee | 37.1718068 | -94.848207 |
| **E1798** | *Heuchera richardsonii* | RICHARDSONII | richardsonii | *Heuchera richardsonii* | Kukla 91 | NCU | NCU00196902 | USA | MN | Clay | 46.8994904 | -96.50882 |
| **E1799** | *Heuchera richardsonii* | GRAYANA | richardsonii | *Heuchera richardsonii* | Steyermark 84538 | NCU | NCU00196904 | USA | MO | Andrew | 39.905814 | -94.730975 |
| **E1800** | *Heuchera richardsonii* | GRAYANA | richardsonii | *Heuchera richardsonii* | Palmer 51751 | NCU | NCU00196905 | USA | MO | Dade | 37.4323036 | -93.84063 |
| **E1801** | *Heuchera richardsonii* | RICHARDSONII | hirsuticaulis_red | *Heuchera grayana* | Redfearn 14465 | NCU | NCU00196906 | USA | MO | Sainte Geneveive | 37.781647 | -90.284423 |
| **E1802** | *Heuchera richardsonii* | RICHARDSONII | richardsonii | *Heuchera richardsonii* | Stephens 23426 | NCU | NCU00196908 | USA | ND | Golden Valley | 46.9317944 | -96.947075 |
| **E1804** | *Heuchera richardsonii* | GRAYANA | richardsonii | *Heuchera richardsonii* | Wallis 6866 | NCU | NCU00196910 | USA | OK | Ottawa | 36.835764 | -94.802681 |
| **E1805** | *Heuchera richardsonii* | RICHARDSONII | richardsonii | *Heuchera richardsonii* | Uttal 9346 | NCU | NCU00196911 | USA | SD | Custer | 43.684943 | -103.46225 |
| **E1852** | *Heuchera americana var. americana* | AMERICANA | caroliniana | *Heuchera americana var. americana* | Radford 6853 | NCU | NCU00181741 | USA | NC | Bladen | 34.627971 | -78.562771 |

Table S1: Continued

| | | | | | | | | | | | | |
|---|---|---|---|---|---|---|---|---|---|---|---|---|
| **E1853** | *Heuchera americana var. americana* | AMERICANA | americana_green | *Heuchera americana var. americana* | McCormick s.n. | NCU | NCU00075922 | USA | NC | Buncombe | 35.64484 | -82.282619 |
| **E1854** | *Heuchera americana var. americana* | AMERICANA | americana_green | *Heuchera americana var. americana* | Bradford 0011 | NCU | NCU00181746 | USA | NC | Burke | 35.834001 | -81.711678 |
| **E1855** | *Heuchera americana var. americana* | AMERICANA | caroliniana | *Heuchera americana var. americana* | Bell 11904 | NCU | NCU00180302 | USA | NC | Caswell | 36.343309 | -79.438356 |
| **E1856** | *Heuchera americana var. americana* | AMERICANA | caroliniana | *Heuchera americana var. americana* | Crutchfield 1304 | NCU | NCU00180304 | USA | NC | Chatham | 35.757001 | -79.088078 |
| **E1857** | *Heuchera americana var. americana* | AMERICANA | caroliniana | *Heuchera americana var. americana* | Ahles 57961 | NCU | NCU00180308 | USA | NC | Durham | 36.073411 | -78.873341 |
| **E1858** | *Heuchera americana var. americana* | BREVIPETALA | fumosimontana | *Heuchera americana var. americana* | Radford 13285 | NCU | NCU00180310 | USA | NC | Graham | 35.44307 | -83.937459 |
| **E1859** | *Heuchera americana var. americana* | AMERICANA | americana_green | *Heuchera americana var. americana* | Radford 13212 | NCU | NCU00180311 | USA | NC | Graham | 36.0690258 | -79.400576 |
| **E1860** | *Heuchera americana var. americana* | AMERICANA | caroliniana | *Heuchera americana var. americana* | Radford 43889 | NCU | NCU00180312 | USA | NC | Granville | 36.084312 | -78.746671 |

Table S1: Continued

| | | | | | | | | | | | | |
|---|---|---|---|---|---|---|---|---|---|---|---|---|
| **E1862** | *Heuchera americana var. americana* | AMERICANA | caroliniana | *Heuchera americana var. americana* | Downs 13390 | NCU | NCU00014279 | USA | NC | Harnett | 35.468214 | -78.898074 |
| **E1863** | *Heuchera americana var. americana* | AMERICANA | americana_green | *Heuchera americana var. americana* | Wells 3284 | NCU | NCU00181750 | USA | NC | Haywood | 35.645284 | -82.941249 |
| **E1864** | *Heuchera americana var. americana* | AMERICANA | americana_green | *Heuchera americana var. americana* | Ramseur 3329 | NCU | NCU00181749 | USA | NC | Haywood | 35.409551 | -82.856243 |
| **E1866** | *Heuchera americana var. americana* | AMERICANA | caroliniana | *Heuchera americana var. americana* | Radford 5793 | NCU | NCU00181757 | USA | NC | Hertford | 36.468816 | -77.095399 |
| **E1868** | *Heuchera americana var. americana* | AMERICANA | caroliniana | *Heuchera americana var. americana* | Stewart 405 | NCU | NCU00180318 | USA | NC | Lee | 35.4691746 | -79.154764 |
| **E1869** | *Heuchera americana var. americana* | AMERICANA | caroliniana | *Heuchera americana var. americana* | Kessler 222 | NCU | NCU00180319 | USA | NC | Lee | 35.5386 | -79.246693 |
| **E1870** | *Heuchera americana var. americana* | AMERICANA | caroliniana | *Heuchera americana var. americana* | Wells 3282 | NCU | NCU00180320 | USA | NC | Lee | 35.4691746 | -79.154764 |
| **E1871** | *Heuchera americana var. americana* | AMERICANA | caroliniana | *Heuchera americana var. americana* | Kessler 242 | NCU | NCU00180321 | USA | NC | Lee | 35.536819 | -79.252642 |

Table S1: Continued

| | | | | | | | | | | | | |
|---|---|---|---|---|---|---|---|---|---|---|---|---|
| **E1872** | *Heuchera americana var. americana* | AMERICANA | americana_green | *Heuchera americana var. americana* | Wells 3283 | NCU | NCU00180322 | USA | NC | Macon | 35.1436639 | -83.39773 |
| **E1873** | *Heuchera americana var. americana* | AMERICANA | fumosimontana | *Heuchera americana var. americana* | Radford s.n. | NCU | NCU00180323 | USA | NC | Macon | 35.148812 | -83.300076 |
| **E1874** | *Heuchera americana var. americana* | AMERICANA | caroliniana | *Heuchera americana var. americana* | Wells 3287 | NCU | NCU00180326 | USA | NC | Madison | 35.8482034 | -82.693106 |
| **E1875** | *Heuchera americana var. americana* | AMERICANA | americana_green | *Heuchera americana var. americana* | Wells 3285 | NCU | NCU00180328 | USA | NC | Madison | 35.737329 | -82.869151 |
| **E1876** | *Heuchera americana var. americana* | AMERICANA | caroliniana | *Heuchera americana var. americana* | Treiber 425 | NCU | NCU00180331 | USA | NC | Martin | 35.966359 | -77.209684 |
| **E1877** | *Heuchera americana var. americana* | AMERICANA | americana_green | *Heuchera americana var. americana* | Leonard 4793 | NCU | NCU00180332 | USA | NC | McDowell | 35.6608869 | -82.048217 |
| **E1879** | *Heuchera americana var. americana* | AMERICANA | americana_green | *Heuchera americana var. americana* | Wells 3288 | NCU | NCU00180334 | USA | NC | Mitchell | 36.0001181 | -82.134903 |
| **E1880** | *Heuchera americana var. americana* | AMERICANA | americana_green | *Heuchera americana var. americana* | Ahles 43133 | NCU | NCU00180335 | USA | NC | Mitchell | 36.0001181 | -82.134903 |

Table S1: Continued

| | | | | | | | | | | | | |
|---|---|---|---|---|---|---|---|---|---|---|---|---|
| **E1881** | *Heuchera americana var. americana* | AMERICANA | caroliniana | *Heuchera americana var. americana* | Radford 42974 | NCU | NCU00180339 | USA | NC | Montgomery | 35.3299572 | -79.897902 |
| **E1882** | *Heuchera americana var. americana* | AMERICANA | caroliniana | *Heuchera americana var. americana* | Kessler 272 | NCU | NCU00078066 | USA | NC | Moore | 35.3054614 | -79.476124 |
| **E1883** | *Heuchera americana var. americana* | AMERICANA | caroliniana | *Heuchera americana var. americana* | Ahles 11797 | NCU | NCU00180340 | USA | NC | Nash | 35.992983 | -77.975531 |
| **E1884** | *Heuchera americana var. americana* | AMERICANA | caroliniana | *Heuchera americana var. americana* | Ahles 41790 | NCU | NCU00180341 | USA | NC | Northampton | 36.4168078 | -77.364223 |
| **E1885** | *Heuchera americana var. americana* | AMERICANA | caroliniana | *Heuchera americana var. americana* | Munch s.n. | NCU | NCU00061747 | USA | NC | Orange | 35.914781 | -79.039 |
| **E1887** | *Heuchera americana var. americana* | AMERICANA | caroliniana | *Heuchera americana var. americana* | Wells 3281 | NCU | NCU00180343 | USA | NC | Orange | 36.0605095 | -79.117268 |
| **E1888** | *Heuchera americana var. americana* | AMERICANA | caroliniana | *Heuchera americana var. americana* | Radford 7569 | NCU | NCU00180344 | USA | NC | Orange | 36.0605095 | -79.117268 |
| **E1889** | *Heuchera americana var. americana* | AMERICANA | caroliniana | *Heuchera americana var. americana* | Larke 1193 | NCU | NCU00180345 | USA | NC | Orange | 35.891961 | -79.037875 |

Table S1: Continued

| | | | | | | | | | | | | |
|---|---|---|---|---|---|---|---|---|---|---|---|---|
| **E1893** | *Heuchera americana var. americana* | AMERICANA | caroliniana | *Heuchera americana var. americana* | White s.n. | NCU | NCU00180353 | USA | NC | Orange | 36.0605095 | -79.117268 |
| **E1894** | *Heuchera americana var. americana* | AMERICANA | caroliniana | *Heuchera americana var. americana* | Wickland 904 | NCU | NCU00180357 | USA | NC | Randolph | 35.738333 | -80.020833 |
| **E1895** | *Heuchera americana var. americana* | AMERICANA | caroliniana | *Heuchera americana var. americana* | Sorrie 9714 | NCU | NCU00180358 | USA | NC | Richmond | 35.0288383 | -79.733326 |
| **E1896** | *Heuchera americana var. americana* | AMERICANA | caroliniana | *Heuchera americana var. americana* | Radford 11479 | NCU | NCU00180359 | USA | NC | Richmond | 35.0288383 | -79.733326 |
| **E1899** | *Heuchera americana var. americana* | AMERICANA | fumosimontana | *Heuchera americana var. americana* | Boufford 13648 | NCU | NCU00180482 | USA | NC | Swain | 35.45819 | -83.466275 |
| **E1900** | *Heuchera americana var. americana* | AMERICANA | caroliniana | *Heuchera americana var. americana* | Ahles 56581 | NCU | NCU00180483 | USA | NC | Wake | 35.7979355 | -78.611831 |
| **E1901** | *Heuchera americana var. americana* | AMERICANA | caroliniana | *Heuchera americana var. americana* | Bell 2911 | NCU | NCU00180484 | USA | NC | Warren | 36.3901694 | -78.105212 |
| **E1902** | *Heuchera americana var. americana* | AMERICANA | americana_green | *Heuchera americana var. americana* | Downs 13696 | NCU | NCU00169493 | USA | NC | Wilkes | 36.17371 | -81.169751 |

Table S1: Continued

| | | | | | | | | | | | | |
|---|---|---|---|---|---|---|---|---|---|---|---|---|
| **E1903** | *Heuchera americana var. americana* | AMERICANA | americana_green | *Heuchera americana var. americana* | Stewart s.n. | NCU | NCU00180488 | USA | NC | Wilkes | 36.1998247 | -81.134135 |
| **E1904** | *Heuchera americana var. americana* | AMERICANA | americana_green | *Heuchera americana var. americana* | Ahles 42660 | NCU | NCU00180490 | USA | NC | Yancey | 35.913356 | -82.455335 |
| **E1905** | *Heuchera caroliniana* | | americana_green | *Heuchera caroliniana* | Radford 1351 | NCU | NCU00053715 | USA | NC | Alexander | 35.972354 | -81.108692 |
| **E1911** | *Heuchera caroliniana* | | caroliniana | *Heuchera caroliniana* | Ahles 41083 | NCU | NCU00053722 | USA | NC | Iredell | 35.604503 | -80.905932 |
| **E1915** | *Heuchera caroliniana* | | caroliniana | *Heuchera caroliniana* | Williams s.n. | NCU | NCU00053725 | USA | NC | Mecklenburg | 35.2356385 | -80.813949 |
| **E1916** | *Heuchera americana var. americana* | AMERICANA | caroliniana | *Heuchera americana var. americana* | Sorrie 13301 | NCU | NCU00439687 | USA | NC | Moore | 35.3054614 | -79.476124 |
| **E1921** | *Heuchera caroliniana* | | caroliniana | *Heuchera caroliniana* | Radford 11873 | NCU | NCU00053736 | USA | NC | Stanly | 35.3235477 | -80.239137 |
| **E1924** | *Heuchera caroliniana* | | caroliniana | *Heuchera caroliniana* | Radford 12001 | NCU | NCU00053739 | USA | NC | Union | 34.9795158 | -80.512821 |
| **E1926** | *Heuchera americana var. hispida* | HISPIDA | americana_green | *Heuchera americana var. hispida* | Poindexter 08-460 | NCU | NCU00128800 | USA | NC | Alleghany | 36.5691944 | 81.1784167 |

Table S1: Continued

| | | | | | | | | | | | | |
|---|---|---|---|---|---|---|---|---|---|---|---|---|
| **E1927** | *Heuchera americana var. hispida* | HISPIDA | americana_green | *Heuchera americana var. hispida* | Poindexter 08-350 | NCU | NCU00128901 | USA | NC | Alleghany | 36.5703611 | 81.4518889 |
| **E1928** | *Heuchera americana var. hispida* | HISPIDA | americana_green | *Heuchera americana var. hispida* | Radford 13141 | NCU | NCU00180492 | USA | NC | Surry | 36.551292 | -80.909589 |
| **E1936** | *Heuchera pubescens* | | americana_green | *Heuchera pubescens* | McCurdy 449 | NCU | NCU00064407 | USA | NC | Stokes | 36.4120995 | -80.228809 |
| **E306-2** | *Heuchera alba* | | alba_lavender | *Heuchera alba* | USDA GRIN: Ames 34945 | Sent directly | | USA | WV | Grant | 39.0034 | -79.2204 |
| **E35** | *Heuchera caroliniana* | | caroliniana | *Heuchera caroliniana* | Haesloop 504 | UNA | UNA00014676 | USA | NC | Stanly | 35.3235477 | -80.239137 |
| **E36** | *Heuchera americana var. americana* | AMERICANA | caroliniana | *Heuchera americana var. americana* | Horn 1644 | UNA | UNA00014678 | USA | SC | Newberry | 34.3266879 | -81.583009 |
| **E438** | *Heuchera americana var. americana* | BREVIPETALA | brevipetala_pink | *Heuchera americana var. brevipetala* | Browne 70K14.5 | MEM | | USA | KY | Pulaski | 37.1124781 | -84.593893 |
| **E439** | *Heuchera americana var. americana* | BREVIPETALA | brevipetala_pink | *Heuchera americana var. brevipetala* | Gentry 926 | MEM | | USA | KY | Henry | 38.447983 | -85.119117 |

Table S1: Continued

| | | | | | | | | | | | | |
|---|---|---|---|---|---|---|---|---|---|---|---|---|
| **E440** | *Heuchera americana var. americana* | HIRSUTICAULIS | brevipetala_brown | *Heuchera americana var. brevipetala* | Bates 2064 | MEM | | USA | TN | Decatur | 35.6067875 | -88.108398 |
| **E441** | *Heuchera americana var. hirsuticaulis* | HIRSUTICAULIS | brevipetala_brown | *Heuchera hirsuticaulis* | Bates 1758 | MEM | | USA | TN | Hardeman | 35.1743058 | -88.996336 |
| **E442** | *Heuchera americana var. hirsuticaulis* | HIRSUTICAULIS | brevipetala_brown | *Heuchera hirsuticaulis* | Bates 2205 | MEM | | USA | TN | McNairy | 35.1663375 | -88.576617 |
| **E443** | *Heuchera americana var. americana* | BREVIPETALA | fumosimontana | *Heuchera americana var. americana* | Athey 3076 | MEM | | USA | KY | Bell | 36.7370344 | -83.64917 |
| **E490** | *Heuchera americana var. americana* | BREVIPETALA | hirsuticaulis_red | *Heuchera americana var. brevipetala* | Holmes 10831 | TEX | 211832 | USA | TX | Bowie | 33.4198886 | -94.447963 |
| **E491** | *Heuchera americana var. americana* | BREVIPETALA | hirsuticaulis_red | *Heuchera americana var. brevipetala* | Correll 37142 | TEX | 459919 | USA | TX | Bowie | 33.4198886 | -94.447963 |
| **E492** | *Heuchera americana var. americana* | BREVIPETALA | hirsuticaulis_red | *Heuchera americana var. brevipetala* | Correll 31281 | TEX | 352864 | USA | TX | Bowie | 33.4198886 | -94.447963 |
| **E498** | *Heuchera americana var. hirsuticaulis* | HIRSUTICAULIS | hirsuticaulis_blue | *Heuchera hirsuticaulis* | D'Arcy 4497 | TEX | | USA | AR | Yell | 35.221944 | -93.243611 |

Table S1: Continued

| | | | | | | | | | | | | |
|---|---|---|---|---|---|---|---|---|---|---|---|---|
| **E499** | *Heuchera americana var. hirsuticaulis* | HIRSUTICAULIS | hirsuticaulis_red | *Heuchera hirsuticaulis* | Kral 67083 | TEX | | USA | AR | Garland | 34.5488944 | -93.183854 |
| **E500** | *Heuchera americana var. hirsuticaulis* | HIRSUTICAULIS | hirsuticaulis_red | *Heuchera hirsuticaulis* | Krall 61777 | TEX | | USA | AR | Howard | 34.0744088 | -93.974578 |
| **E501** | *Heuchera americana var. hirsuticaulis* | HIRSUTICAULIS | hirsuticaulis_red | *Heuchera hirsuticaulis* | Kral 59876 | TEX | | USA | AR | Faulkner | 35.1470851 | -92.321905 |
| **E502** | *Heuchera americana var. americana* | CALYCOSA | calycosa_yellow | *Heuchera americana var. alabamense* | Orzell 9505 | TEX | | USA | AL | Cleburne | 33.721111 | -85.604444 |
| **E503** | *Heuchera americana var. hirsuticaulis* | HIRSUTICAULIS | hirsuticaulis_red | *Heuchera hirsuticaulis* | Raven 27489 | TEX | | USA | MO | Jefferson | 38.125 | -90.675 |
| **E505** | *Heuchera americana var. americana* | CALYCOSA | calycosa_yellow | *Heuchera americana var. alabamense* | Pyron 2492 | TEX | | USA | GA | Burke | 33.0482247 | -81.957598 |
| **E506** | *Heuchera americana var. americana* | AMERICANA | caroliniana | *Heuchera americana var. americana* | Moldenke 30033 | TEX | | USA | VA | Amelia | 37.3319664 | -78.008448 |
| **E507** | *Heuchera americana var. americana* | AMERICANA | americana_green | *Heuchera americana var. americana* | Woodbury s.n. | TEX | | USA | NC | Rutherford | 35.4833 | -81.95 |

Table S1: Continued

| | | | | | | | | | | | | |
|---|---|---|---|---|---|---|---|---|---|---|---|---|
| **E509** | *Heuchera americana var. americana* | BREVIPETALA | hirsuticaulis_red | *Heuchera americana var. brevipetala* | Shoals 30506 | TEX | | USA | AR | Cleburne | 35.5298937 | -92.031282 |
| **E510** | *Heuchera americana var. americana* | AMERICANA | caroliniana | *Heuchera americana var. americana* | Brown 2135 | TEX | | USA | NC | Durham | 35.996653 | -78.901805 |
| **E511** | *Heuchera americana var. americana* | CALYCOSA | brevipetala_brown | *Heuchera americana var. brevipetala* | Crawford 223 | TEX | | USA | AL | Fayette | 33.7303493 | -87.741909 |
| **E513** | *Heuchera americana var. hirsuticaulis* | HIRSUTICAULIS | hirsuticaulis_red | *Heuchera hirsuticaulis* | Randrianaivo 423 | TEX | | USA | MO | Saint Genevieve | 37.828889 | -90.225556 |
| **E514** | *Heuchera americana var. americana* | CALYCOSA | calycosa_yellow | *Heuchera americana var. alabamense* | Kral 63437 | TEX | | USA | GA | Floyd | 34.2421984 | -85.222925 |
| **E515** | *Heuchera americana var. americana* | BREVIPETALA | fumosimontana | *Heuchera americana var. americana* | Wofford 96-38 | TEX | | USA | TN | Grainger | 36.2810821 | -83.510702 |
| **E516** | *Heuchera americana var. americana* | AMERICANA | caroliniana | *Heuchera americana var. americana* | Haesloop 504 | TEX | | USA | NC | Stanly | 35.3235477 | -80.239137 |
| **E517** | *Heuchera americana var. americana* | BREVIPETALA | brevipetala_pink | *Heuchera americana var. brevipetala* | Roller 1/60 | TEX | | USA | KY | Madison | 37.7298081 | -84.297474 |

Table S1: Continued

| | | | | | | | | | | | | |
|---|---|---|---|---|---|---|---|---|---|---|---|---|
| **E518** | *Heuchera americana var. americana* | BREVIPETALA | fumosimontana | *Heuchera americana var. americana* | Moldenke 27133 | TEX | | USA | TN | Blount | 35.6719722 | -83.931447 |
| **E519** | *Heuchera americana var. americana* | AMERICANA | caroliniana | *Heuchera americana var. americana* | Moldenke 27000 | TEX | | USA | SC | McCormick | 33.911441 | -82.295433 |
| **E521** | *Heuchera americana var. hirsuticaulis* | HIRSUTICAULIS | hirsuticaulis_blue | *Heuchera hirsuticaulis* | Hardin 634 | TEX | | USA | AR | Franklin | 35.5110075 | -93.886485 |
| **E522** | *Heuchera americana var. americana* | CALYCOSA | calycosa_yellow | *Heuchera americana var. alabamense* | McVaugh 8610 | TEX | | USA | AL | Randolph | 33.2651763 | -85.469396 |
| **E523** | *Heuchera americana var. americana* | BREVIPETALA | brevipetala_pink | *Heuchera americana var. brevipetala* | Bright 15210 | TEX | | USA | PA | Allegheny | 40.4597204 | -79.976041 |
| **E524** | *Heuchera americana var. americana* | BREVIPETALA | fumosimontana | *Heuchera americana var. americana* | McNeilus 97-384 | TEX | | USA | TN | Campbell | 36.3865389 | -84.135162 |
| **E542** | *Heuchera richardsonii* | GRAYANA | grayana_orange | *Heuchera grayana* | Chase 9546 | TEX | | USA | IL | McHenry | 42.3294391 | -88.460571 |
| **E543** | *Heuchera richardsonii* | GRAYANA | richardsonii | *Heuchera richardsonii* | Welch 9839 | TEX | | USA | IA | Dickinson | 43.3753018 | -95.165428 |
| **E639** | *Heuchera richardsonii* | RICHARDSONII | richardsonii | *Heuchera richardsonii* | Breitung 1139 | TEX | | Canada | Saskatchewan | Wallwort | 52.57045 | -104.01686 |

Table S1: Continued

| | | | | | | | | | | | | |
|---|---|---|---|---|---|---|---|---|---|---|---|---|
| **E640** | *Heuchera richardsonii* | GRAYANA | richardsonii | *Heuchera richardsonii* | Chase 14460 | TEX | | USA | IL | Peoria | 41.4194058 | -89.589101 |
| **E641** | *Heuchera richardsonii* | RICHARDSONII | grayana_orange | *Heuchera grayana* | Chase 11914 | TEX | | USA | IL | Will | 45.9009106 | -87.999475 |
| **FL-25** | *Heuchera longiflora var. longiflora* | | fumosimontana | *Heuchera longiflora var. longiflora* | Floden 3208 | OS | | USA | VA | Lee | 36.6679833 | -83.230925 |
| **FL-26** | *Heuchera longiflora var. longiflora* | | fumosimontana | *Heuchera longiflora var. longiflora* | Floden s.n. | OS | | USA | AL | Talladega | 33.1415972 | -86.258378 |
| **H105-1** | *Heuchera richardsonii* | RICHARDSONII | richardsonii | *Heuchera richardsonii* | Folk 105 | OS | | USA | SD | Pennington | 43.8438361 | -102.43763 |
| **H106-1** | *Heuchera richardsonii* | RICHARDSONII | richardsonii | *Heuchera richardsonii* | Folk 106 | OS | | USA | SD | Pennington | 44.4666667 | -102.0525 |
| **H106-2** | *Heuchera richardsonii* | RICHARDSONII | richardsonii | *Heuchera richardsonii* | Folk 106 | OS | | USA | SD | Pennington | 44.4666667 | -102.0525 |
| **H166-2** | *Heuchera richardsonii* | GRAYANA | richardsonii | *Heuchera richardsonii* | Folk 166 | OS | | USA | IA | Boone | 41.9966667 | -93.883611 |
| **H167-1** | *Heuchera richardsonii* | GRAYANA | grayana_orange | *Heuchera grayana* | Folk 167 | OS | | USA | IL | Porter | 41.3213889 | -87.006389 |
| **H167-2** | *Heuchera richardsonii* | GRAYANA | grayana_orange | *Heuchera grayana* | Folk 167 | OS | | USA | IL | Porter | 41.3213889 | -87.006389 |

Table S1: Continued

| | | | | | | | | | | | | |
|---|---|---|---|---|---|---|---|---|---|---|---|---|
| **H177-2** | *Heuchera americana var. hirsuticaulis* | HIRSUTICAULIS | hirsuticaulis_red | *Heuchera hirsuticaulis* | Folk 177 | OS | | USA | IL | Union | 37.5415333 | -89.426917 |
| **H246** | *Heuchera americana var. americana* | BREVIPETALA | brevipetala_pink | *Heuchera americana var. brevipetala* | Folk 246 | OS | | USA | OH | Delaware | 40.267491 | -82.951414 |
| **H59-2** | *Heuchera americana var. hirsuticaulis* | HIRSUTICAULIS | hirsuticaulis_red | *Heuchera hirsuticaulis* | Folk 59 | OS | | USA | MO | Shannon | 37.39023 | -91.45272 |
| **H59-2** | *Heuchera americana var. hirsuticaulis* | HIRSUTICAULIS | hirsuticaulis_red | *Heuchera hirsuticaulis* | Folk 59 | OS | | USA | MO | Shannon | 37.39023 | -91.45272 |
| **H62-1** | *Heuchera americana var. hirsuticaulis* | HIRSUTICAULIS | hirsuticaulis_red | *Heuchera hirsuticaulis* | Folk 62 | OS | | USA | MO | Scott | 37.19118 | 89.63458 |
| **H62-2** | *Heuchera americana var. hirsuticaulis* | HIRSUTICAULIS | hirsuticaulis_red | *Heuchera hirsuticaulis* | Folk 62 | OS | | USA | MO | Scott | 37.19118 | 89.63458 |
| **H62-2** | *Heuchera americana var. hirsuticaulis* | HIRSUTICAULIS | hirsuticaulis_red | *Heuchera hirsuticaulis* | Folk 62 | OS | | USA | MO | Scott | 37.19118 | -89.63458 |
| **H62-4** | *Heuchera americana var. hirsuticaulis* | HIRSUTICAULIS | hirsuticaulis_red | *Heuchera hirsuticaulis* | Folk 62 | OS | | USA | MO | Scott | 37.19118 | 89.63458 |

Table S1: Continued

| | | | | | | | | | | | |
|---|---|---|---|---|---|---|---|---|---|---|---|
| **H62-4** | *Heuchera americana var. hirsuticaulis* | HIRSUTICAULIS | hirsuticaulis_red | *Heuchera hirsuticaulis* | Folk 62 | OS | USA | MO | Scott | 37.19118 | -89.63458 |
| **H62-5** | *Heuchera americana var. hirsuticaulis* | HIRSUTICAULIS | hirsuticaulis_red | *Heuchera hirsuticaulis* | Folk 62 | OS | USA | MO | Scott | 37.19118 | 89.63458 |
| **H62-5** | *Heuchera americana var. hirsuticaulis* | HIRSUTICAULIS | hirsuticaulis_red | *Heuchera hirsuticaulis* | Folk 62 | OS | USA | MO | Scott | 37.19118 | -89.63458 |
| **H62-6** | *Heuchera americana var. hirsuticaulis* | HIRSUTICAULIS | hirsuticaulis_red | *Heuchera hirsuticaulis* | Folk 62 | OS | USA | MO | Scott | 37.19118 | 89.63458 |
| **H62-6** | *Heuchera americana var. hirsuticaulis* | HIRSUTICAULIS | hirsuticaulis_red | *Heuchera hirsuticaulis* | Folk 62 | OS | USA | MO | Scott | 37.19118 | -89.63458 |
| **H71** | *Heuchera americana var. americana* | BREVIPETALA | brevipetala_pink | *Heuchera americana var. brevipetala* | Folk 71 | OS | USA | OH | Lawrence | 38.576956 | -82.335822 |
| **H95** | *Heuchera americana var. americana* | AMERICANA | americana_green | *Heuchera americana var. americana* | Folk 95 | OS | USA | NC | Buncombe | 35.625553 | -82.518968 |
| **I173** | *Heuchera alba* | | alba_lavender | *Heuchera alba* | Vanderhorst 7485 | MISSA | USA | WV | Grant | 38.927459 | -79.237163 |
| **I174** | *Heuchera alba* | | alba_lavender | *Heuchera alba* | Byers 1570 | MISSA | USA | WV | Grant | 39.000504 | -79.081646 |

Table S1: Continued

| | | | | | | | | | | | |
|---|---|---|---|---|---|---|---|---|---|---|---|
| **I175** | *Heuchera alba* | | alba_lavender | *Heuchera alba* | Streets 3790 | MISSA | USA | WV | Pocahontas | 38.202504 | -80.194893 |
| **I176** | *Heuchera alba* | | alba_lavender | *Heuchera alba* | Streets 3004 | MISSA | USA | WV | Pendleton | 38.505711 | -79.209617 |
| **I177** | *Heuchera alba* | | alba_lavender | *Heuchera alba* | Streets 5990 | MISSA | USA | WV | Pocahontas | 38.365892 | -79.93746 |
| **I178** | *Heuchera alba* | | alba_lavender | *Heuchera alba* | Streets 4090 | MISSA | USA | WV | Grant | 38.984892 | -79.218757 |
| **I179** | *Heuchera alba* | | alba_lavender | *Heuchera alba* | Streets 4768 | MISSA | USA | WV | Pendleton | 38.636815 | -79.394064 |
| **I180** | *Heuchera alba* | | alba_lavender | *Heuchera alba* | Streets 4778 | MISSA | USA | WV | Pendleton | 38.666659 | -79.235843 |
| **I2** | *Heuchera americana var. americana* | BREVIPETALA | brevipetala_pink | *Heuchera americana var. brevipetala* | Folk I2 | OS | USA | OH | Defiance | 41.174476 | -84.256819 |
| **I20** | *Heuchera americana var. americana* | AMERICANA | filtered out | *Heuchera americana var. americana* | Folk I20 | OS | USA | VA | Albemarle | 38.094964 | -78.489373 |
| **I65** | *Heuchera richardsonii* | GRAYANA | grayana_orange | *Heuchera grayana* | Folk I65 | OS | USA | MI | Berrien | 41.868479 | -86.348772 |
| **I67** | *Heuchera alba* | | alba_lavender | *Heuchera alba* | Folk I67 | OS | USA | WV | Highland | 38.594238 | -79.18181 |
| **I82** | *Heuchera richardsonii* | RICHARDSONII | richardsonii | *Heuchera richardsonii* | Folk I82 | OS | Canada | Alberta | | 49.949883 | -113.97907 |
| **I90-C** | *Heuchera inconstans* | | not included | *Heuchera inconstans* | Folk I90 | OS | USA | AZ | Coconino | 34.99126 | -111.73521 |
| **L1-1** | *Heuchera longiflora var. aceroides* | | americana_green | *Heuchera longiflora var. aceroides* | Folk L-1 | OS | USA | NC | Madison | 35.7467139 | -82.872478 |

Table S1: Continued

| | | | | | | | | | | | |
|---|---|---|---|---|---|---|---|---|---|---|---|
| **L1-3** | *Heuchera longiflora var. aceroides* | americana_green | *Heuchera longiflora var. aceroides* | Folk L-1 | OS | | USA | NC | Madison | 35.7467139 | -82.872478 |
| **L14-1** | *Heuchera longiflora var. longiflora* | fumosimontana | *Heuchera longiflora var. longiflora* | Folk L-14 | OS | | USA | KY | Floyd | 37.6698389 | -82.913272 |
| **L14-2** | *Heuchera longiflora var. longiflora* | fumosimontana | *Heuchera longiflora var. longiflora* | Folk L-14 | OS | | USA | KY | Floyd | 37.6698389 | -82.913272 |
| **L20-1** | *Heuchera longiflora var. longiflora* | fumosimontana | *Heuchera longiflora var. longiflora* | Folk L-20 | OS | | USA | KY | Lee | 37.6304667 | -83.770767 |
| **L20-2** | *Heuchera longiflora var. longiflora* | fumosimontana | *Heuchera longiflora var. longiflora* | Folk L-20 | OS | | USA | KY | Lee | 37.6304667 | -83.770767 |
| **L3-1** | *Heuchera longiflora var. aceroides* | americana_green | *Heuchera longiflora var. aceroides* | Folk L-3 | OS | | USA | NC | Madison | 35.92865 | -82.750078 |
| **L3-2** | *Heuchera longiflora var. aceroides* | americana_green | *Heuchera longiflora var. aceroides* | Folk L-3 | OS | | USA | NC | Madison | 35.92865 | -82.750078 |
| **L6-1** | *Heuchera longiflora var. longiflora* | fumosimontana | *Heuchera longiflora var. longiflora* | Folk L-6 | OS | | USA | VA | Wise | 36.8558167 | -82.747758 |

Table S1: Continued

| | | | | | | | | | | | |
|---|---|---|---|---|---|---|---|---|---|---|---|
| **L6-2** | *Heuchera longiflora var. longiflora* | fumosimontana | *Heuchera longiflora var. longiflora* | Folk L-6 | OS | | USA | VA | Wise | 36.8558167 | -82.747758 |
| **L9-2** | *Heuchera longiflora var. longiflora* | fumosimontana | Heuchera longiflora var. longiflora | Folk L-9 | OS | | USA | VA | Wise | 37.1037417 | -82.668386 |

Table S1: Continued

**Figures**

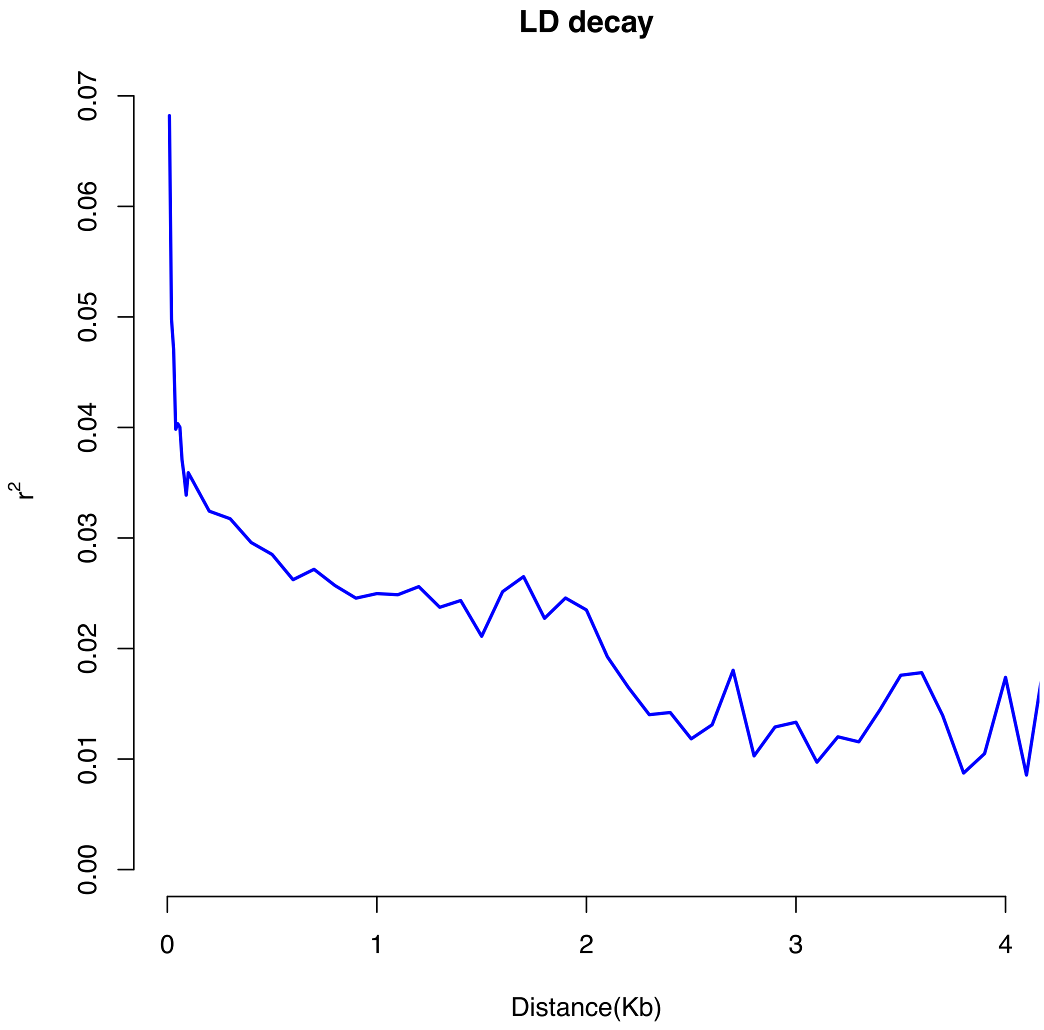

**Figure S1** | Linkage disequilibrium decay graph.

Graph showing linkage disequilibrium (LD) decay with base pair distances on the x-axis and $R^2$ on the y-axis.
